# Chemically induced partial unfolding of the multifunctional Apurinic/apyrimidinic endonuclease 1

**DOI:** 10.1101/2023.06.29.547112

**Authors:** Ratan Rai, Olabode I. Dawodu, Jingwei Meng, Steven M. Johnson, Jonah Z. Vilseck, Mark R. Kelley, Joshua J. Ziarek, Millie M. Georgiadis

## Abstract

Apurinic/apyrimidinic endonuclease I (APE1) acts as both an endonuclease and a redox factor to ensure cell survival. The two activities require different conformations of APE1. As an endonuclease, APE1 is fully folded. As a redox factor, APE1 must be partially unfolded to expose the buried residue Cys65, which reduces transcription factors including AP-1, NF-κB, and HIF-1α and thereby enables them to bind DNA. To determine a molecular basis for partial unfolding associated with APE1’s redox activity, we characterized specific interactions of a known redox inhibitor APX3330 with APE1 through waterLOGSY and ^1^H-^15^N HSQC NMR approaches using ethanol and acetonitrile as co-solvents. We find that APX3330 binds to the endonuclease active site in both co-solvents and to a distant small pocket in acetonitrile. Prolonged exposure of APE1 with APX3330 in acetonitrile resulted in a time-dependent loss of ^1^H-^15^N HSQC chemical shifts (∼35%), consistent with partial unfolding. Regions that are partially unfolded include adjacent N- and C-terminal beta strands within one of the two sheets comprising the core, which converge within the small binding pocket defined by the CSPs. Removal of APX3330 via dialysis resulted in a slow reappearance of the ^1^H-^15^N HSQC chemical shifts suggesting that the effect of APX3330 is reversible. APX3330 significantly decreases the melting temperature of APE1 but has no effect on endonuclease activity using a standard assay in either co-solvent. Our results provide insights on reversible partial unfolding of APE1 relevant for its redox function as well as the mechanism of redox inhibition by APX3330.

**TOC graphic:** 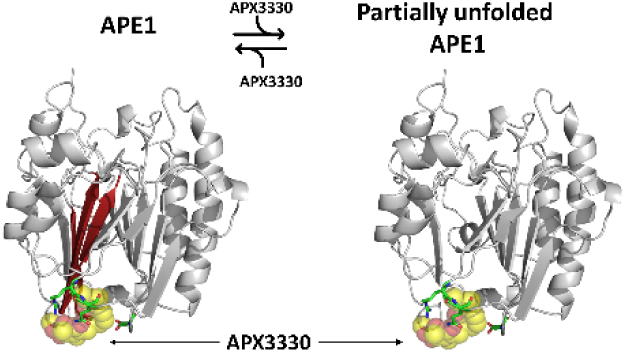

## Introduction

The reliance of cancer cells on robust DNA repair has spurred the development of small molecule inhibitors targeting DNA repair proteins including the multifunctional apurinic/apyrimidinic endonuclease 1/redox effector factor 1 (referred to here as APE1)(32; 34). APE1 is critical for the repair of damaged bases in the base excision repair pathway. Damage to nucleobases from oxidation, alkylation, and spontaneous nucleobase loss in cells is chronic and extensive with estimates of at least 10,000 events within the genome of each cell per day; thus, repair is critical for maintaining genomic stability (37). APE1 is dually targeted; under normal conditions, it is primarily localized in the nucleus with small amounts seen in the cytoplasm and mitochondria,(16) but under oxidative stress, APE1 is relocalized from the nucleus to the cytoplasm and mitochondria (23; 35). In both the nucleus and mitochondria, APE1 functions as an endonuclease to repair damaged DNA. The nuclear localization sequence (NLS) is located on the unstructured N-terminus (1–20) of the protein (29) and the mitochondrial targeting sequence (MTS) on the structured C-terminus (289–318)(35). Exposure of the MTS would necessarily require at least partial unfolding of the protein. Other nuclease functions attributed to APE1 include 3’-5’ exonuclease (11), 3’-phosdiesterase (50), nucleotide incision repair activity (28), and repair of damaged RNA (15; 41).

The redox activity of APE1 was first identified in the search for the factor regulating the redox state of AP-1(66). The proposed mechanism was a thiol-exchange mechanism involving Cys65 as the nucleophile in APE1, which reduced disulfide bonds in AP-1 that prevented it from binding to DNA(64; 21). Subsequently, APE1 was identified as the redox factor responsible for reducing several transcription factors including HIF-1α, NF-kB, STAT3, and others that are critical for cell growth and survival (33; 9; 44). Reduction of TFs is relevant for many diseases; specifically, reduction of HIF-1α, NF-kB, and STAT3 impacts angiogenesis/inflammation associated with retinal vascular diseases (25). Cys65 has also been implicated in mitochondrial trafficking (63).

Knock-down of APE1 in a xenograft mouse model resulted in reduced tumor growth (18) supporting development of small molecule inhibitors targeting this protein. Since knock-down results in loss of both endonuclease and redox functions, development of small molecule inhibitors targeting one or the other function of APE1 has been important in defining the roles of these functions in cancer (44) and more recently in vascular retinal diseases (25; 45). The first small molecule, (2E)-2-[(4,5-dimethoxy-2-methyl-3,6-dioxo-1,4-cyclohexadien-1-yl)methylene]-undecanoic acid (E3330 or APX3330) (Fig. 1), targeting APE1 was identified in 2000 and found to inhibit the redox function of APE1 (26; 54).

**Figure 1.**
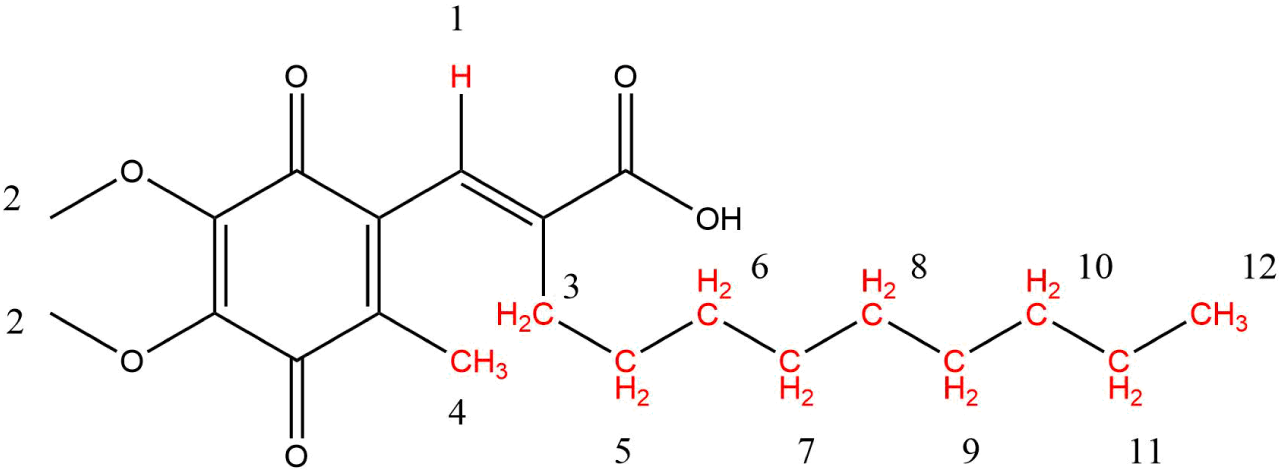
Chemical structure of APX3330 (E3330). Protons with altered solvent accessibility in a 1D WaterLOGSY NMR experiment are highlighted in red. Numbers refer to assignments on the NMR spectra (SI Figure S9).

The mechanism of action of APX3330 remains controversial. In one study, APX3330 dissolved in D6-ethanol was reported through characterization by ^1^H-^15^N HSQC/TROSY NMR experiments to bind to the endonuclease active site and inhibit endonuclease activity (43). In other studies, APX3330 showed no inhibition of APE1 endonuclease activity at concentrations up to 160 μM (39; 67) and was shown to lower the melting temperature of the APE1 (67). Partial unfolding of APE1 was observed by coupling N-ethylmaleimide (NEM) footprinting and intact mass spectrometry. In this study, we observed a time-dependent labeling of buried Cys residues with NEM consistent with partial unfolding of APE1 in the presence of APX3330 (57). The extent of partial unfolding was confirmed through HDX mass spectrometry experiments (67).

There are 7 Cys residues in APE1 and no disulfide bonds; the two solvent accessible Cys residues (99 and 138) were readily modified by NEM in the absence of APX3330. No covalent adducts between APX3330 and APE1 were observed by intact mass spectrometry (57). In the presence of APX3330, we suggested that once Cys residues 65, 93, 208, 296, and 310 (normally buried within the interior of fully folded APE1) were modified with NEM, it would not be possible for APE1 to fully refold. Thus, partially unfolded APE1 molecules were trapped through NEM modification. While incubation of APX3330 resulted in a population of partially unfolded APE1 molecules in these studies (57; 67), the mechanism of partial unfolding has yet to be determined.

In the present study, we explored the effects of APX3330 interactions with APE1 in two different co-solvents, ethanol-D6 and acetonitrile-D3, by ^1^H-^15^N HSQC spectroscopy, ligand-based waterLOGSY NMR, CD, and endonuclease activity assays. Our studies provide a mechanism by which Cys65, which is essential for redox activity and subcellular trafficking of APE1 to the mitochondria, is exposed through partial unfolding of a beta sheet within the core of APE1. Further, our studies provide a basis for classifying redox inhibitors based on mechanism of action, which may help guide future drug development.

## Results

### Solvent interactions with APE1

To assess interactions of solvents with APE1, we first assigned 234 backbone ^1^H-^15^N resonances, representing 90% of the possible 261 backbone resonances in the well-resolved HSQC spectra of ^15^N-labelled APE1 (39-318/C138A), hereafter referred to as APE1, in buffer containing 20 mM sodium phosphate, 0.1 M sodium chloride at pH 6.5 on a 600 MHz Bruker NMR spectrometer at 25 °C (SI Fig. S1). Assignments were made based on previously reported chemical shift assignments for APE1 (42). It is difficult to overstate the similarity between the respective ^1^H-^15^N HSQC/TROSY spectra obtained for [*U*-^15^N] APE1; visual inspection resulted in 234 NH assignments (SI Fig. S2). In the previous report, 235 out of 261 resonances were assigned; the following resonances were not assigned (G80-V84, E87, D90, E101, L104, S120, S123-G130, Q153-R156, G178, D210, T268, and G279). The C138A variant used for our studies was produced for crystallographic studies to avoid disulfide bond formation between Cys138 residues in two APE1 molecules in close proximity within the crystal lattice. We were unable to assign Ala138 since this differs in sequence from the reported work (42). Equally well-resolved ^1^H-^15^N HSQC spectra were obtained in a buffer including 20 mM sodium phosphate, pH 6.5, 0.1M sodium chloride, 0.5 mM DTT, and 0.2 mM EDTA as used in the previous NMR study (42) (SI Fig. S3). We further demonstrated that the APE1 sample was stable for up to 20 days (SI Fig. S4) and produced equally well-resolved spectra at 30 °C and 35 °C (SI Fig. S5).

We next explored interactions of acetonitrile-D3 or ethanol-D6 with APE1 in ^1^H-^15^N HSQC NMR experiments. As a control, interactions of DMSO-D3 with APE1 were also mapped. Chemical shift perturbations were calculated as previously described (43). DMSO interacts extensively with APE1 (SI Fig. S6). Residues with significant CSPs obtained for DMSO added to APE1 include the following: Gly113, Ser135, Arg136, Gln137, Asp152, Val172, Arg187, Asn222, Thr233, Arg237, Tyr262, Tyr264, Asp283, Ala304, and Leu305 (SI Fig. S6). Separately, in crystallographic studies, we had identified two specific binding sites for DMSO on APE1 (58); one in the endonuclease active site and the second in a smaller binding pocket on the opposite face of APE1 (SI Fig. S6). CSPs obtained for DMSO interactions with APE1 are consistent with binding in the vicinity of the active site (Asn222, Ala304, and Leu305) and to the smaller pocket defined by Ser135, Arg136, and Gln137. The smaller pocket was identified as the binding site for a naphthoquinone (43) and in a recent report investigating APE1 inhibitors (52).

Other residues with significant CSPs include the buried residue Asp283, which along with two other buried residues, Ser66 and Gln95, are all hydrogen-bonded to a single water molecule in the crystal structure, PDB ID 4QHD. This water mediated hydrogen-bonding network effectively links the first and second strands of β sheet 1 and fifth strand of β sheet 2 within the β sandwich core of APE1 (SI Fig. S7). Due to the extensive CSPs obtained for DMSO, we excluded DMSO as a usable solvent for mapping compound interactions with APE1.

In contrast to DMSO, ethanol perturbs the chemical environment of Asn174, located in the endonuclease active site (SI Fig. S8), and Ser66, noted in the hydrogen-bonding network above (SI Fig. S7). While it is possible to subtract the CSPs for interactions of ethanol with APE1 as was done in a previous study(43), the physical reality is that ethanol perturbs the chemical environment of N174 within the endonuclease active site of APE1 and that of Ser66 within the β sheet core. No significant CSPs were observed for interactions of acetonitrile with APE1 (SI Fig. S9).

### Interactions of APX3330 with APE1

Interactions with APX3330 (Fig. 1) can be assessed from the perspective of APX3330 as a ligand as well as the protein. To map interactions of APX3330 from the protein perspective, we performed ^1^H-^15^N HSQC experiments for APX3330 dissolved in ethanol-D6 or acetonitrile-D3 using previously reported buffering conditions (43) with a 10-fold molar excess of APX3330:APE1. Consistent with the previous study (43), APE1 CSPs for APX3330 dissolved in ethanol map largely to the endonuclease active site (Fig. 2). Residues with significant CSPs include Asn212, Gly231, Met270, Met271, Asn272, Ala273, Val278, Trp280, and Tyr315. In the previous study, the following residues exhibited significant CSPs: Gly231, Met270, Met271, Asn272, Ala273, Val278, Trp280, and Asp308. Results from our experiment agree well with those of the previous study with good overlap of residues within or near the endonuclease active site (Fig. 2).

**Figure 2.**
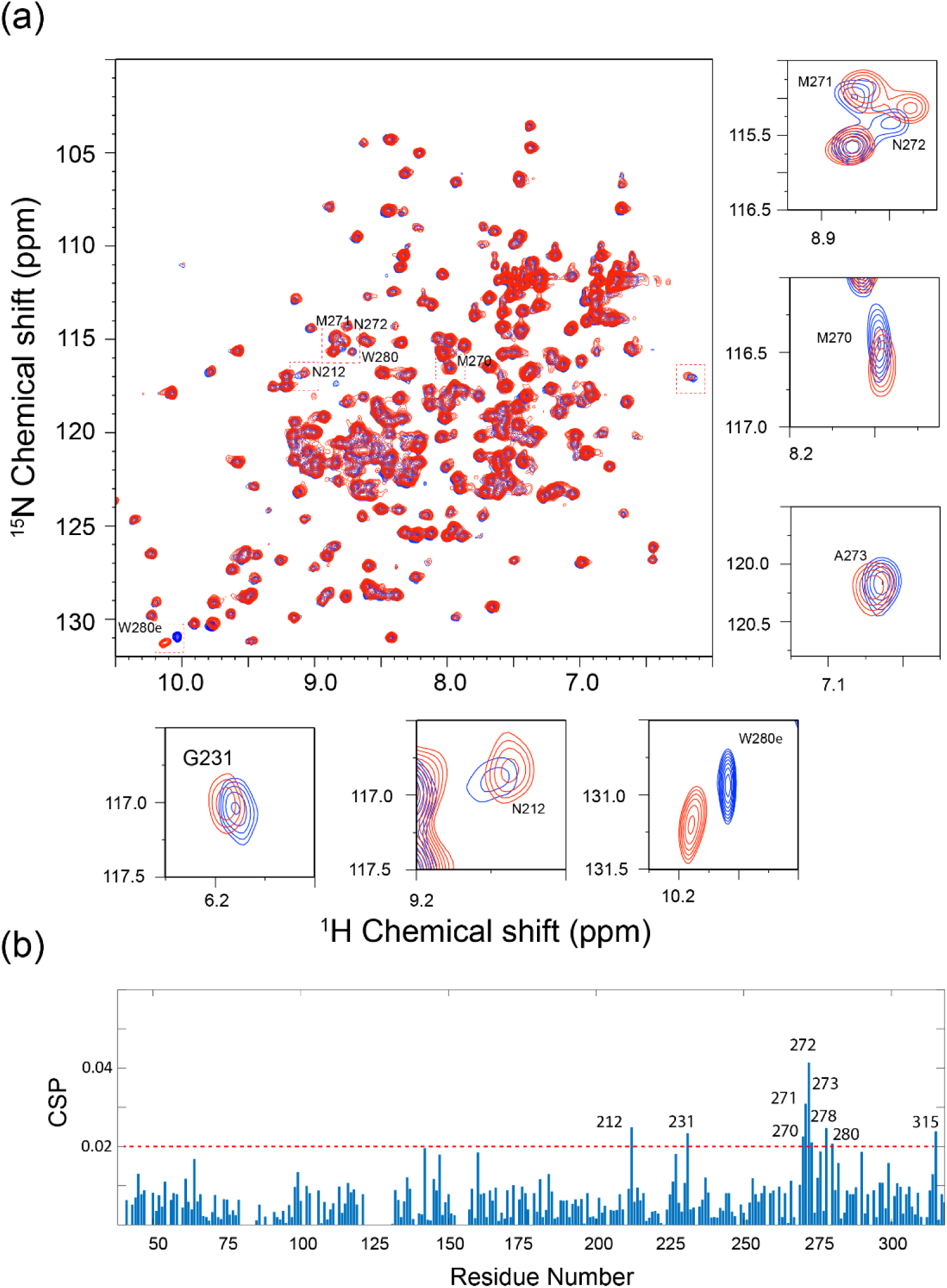
Interaction of APX3330/ethanol results in CSPs within the endonuclease active site of APE1. (a) 2D ^1^H-^15^N HSQC spectral overlay of the APE1 (85 μM) spectrum (black) and with 850 µM APX3330 (red). Each cross peak corresponds to one backbone or side chain amide. Specific chemical shift perturbations are shown in small boxes. (b) The chemical shift perturbations versus residue number reveal widespread interactions with APE1.

CSPs for APX3330 dissolved in D3-acetonitrile define interactions with both the small pocket and the endonuclease active site (Fig. 3). Significant CSPs were observed for the following residues: Ile64, Ser66, Asp70, Arg136, Gln137, Ile146, Ile158, Gly231, Ser252, Tyr264, Met270, Met271, Asn272, Ala273, Trp280, Lys303, and Leu305. Of these, Arg136 and Gln137 are surface residues that define the small pocket on APE1 discussed above, while Asp70, Met270, Met271, Asn272, Ala273, and Trp280 define the endonuclease active site pocket on the opposite face of the protein (Fig. 4a, b). The only other surface residues with significant CSPs are Ile146, Lys303, and Leu305. The remaining residues with CSPs above 0.02 are located within the interior of the protein. Notably, residues Ile64 and Ser66 (as noted above involved in a water-mediated hydrogen-bonding network) reside within a central strand in one of the two beta sheets that comprise the β sandwich core of the protein, followed by a connecting loop that includes residue Asp70. Residues Met270, Met271, and Asn272 reside within a loop adjacent to the endonuclease active site (Fig. 4c-f).

**Figure 3.**
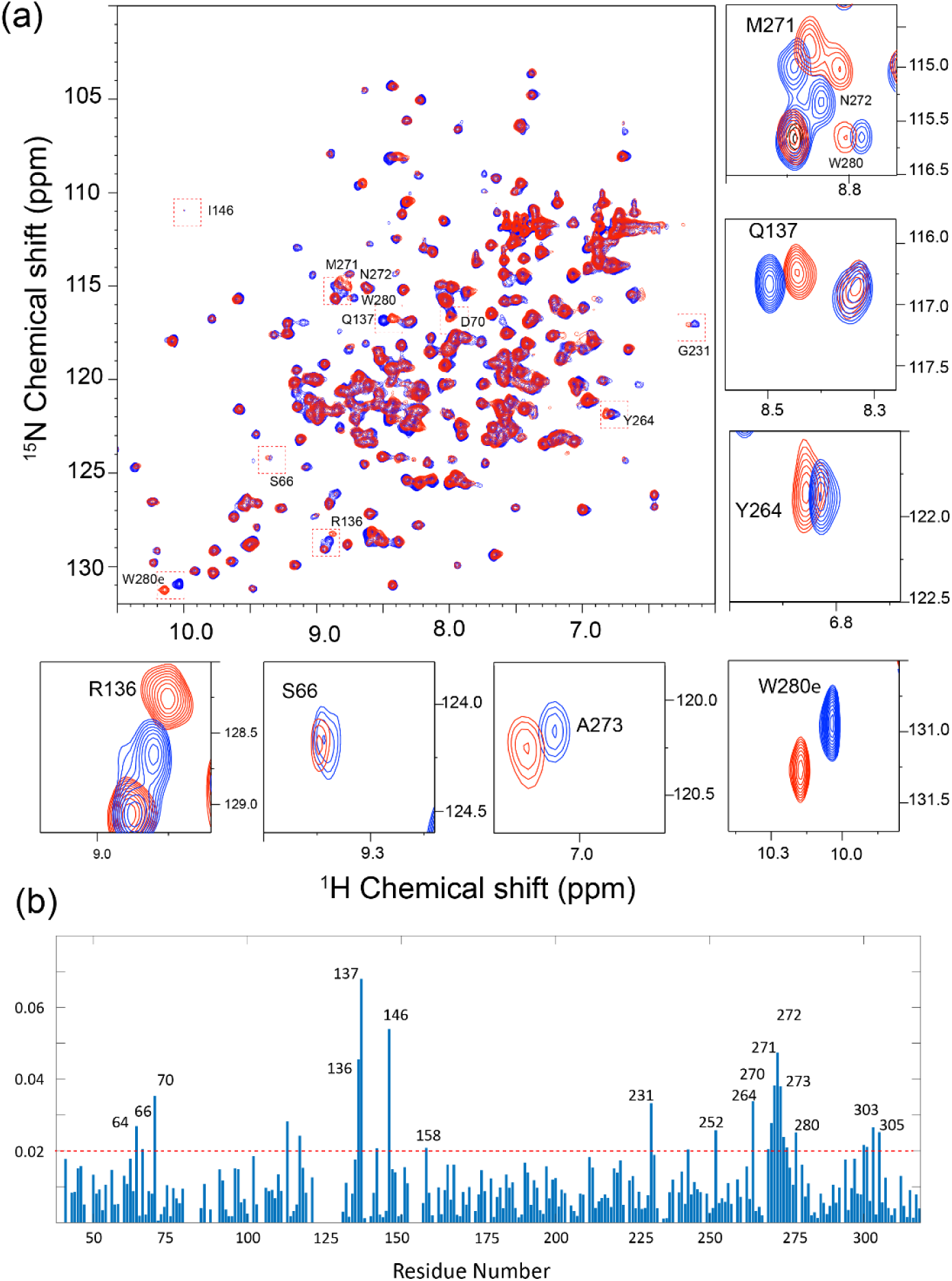
Interaction of APX3330/acetonitrile result in CSPs defining two distinct pockets in APE1. (a) 2D ^1^H-^15^N HSQC spectral overlay of the APE1 (85 μM) spectrum (black) and with 850 µM APX3330 (red). Each cross peak corresponds to one backbone or side chain amide. Specific chemical shift perturbations are shown in small boxes. (b) The chemical shift perturbations versus residue number reveal widespread interactions with APE1.

**Figure 4.**
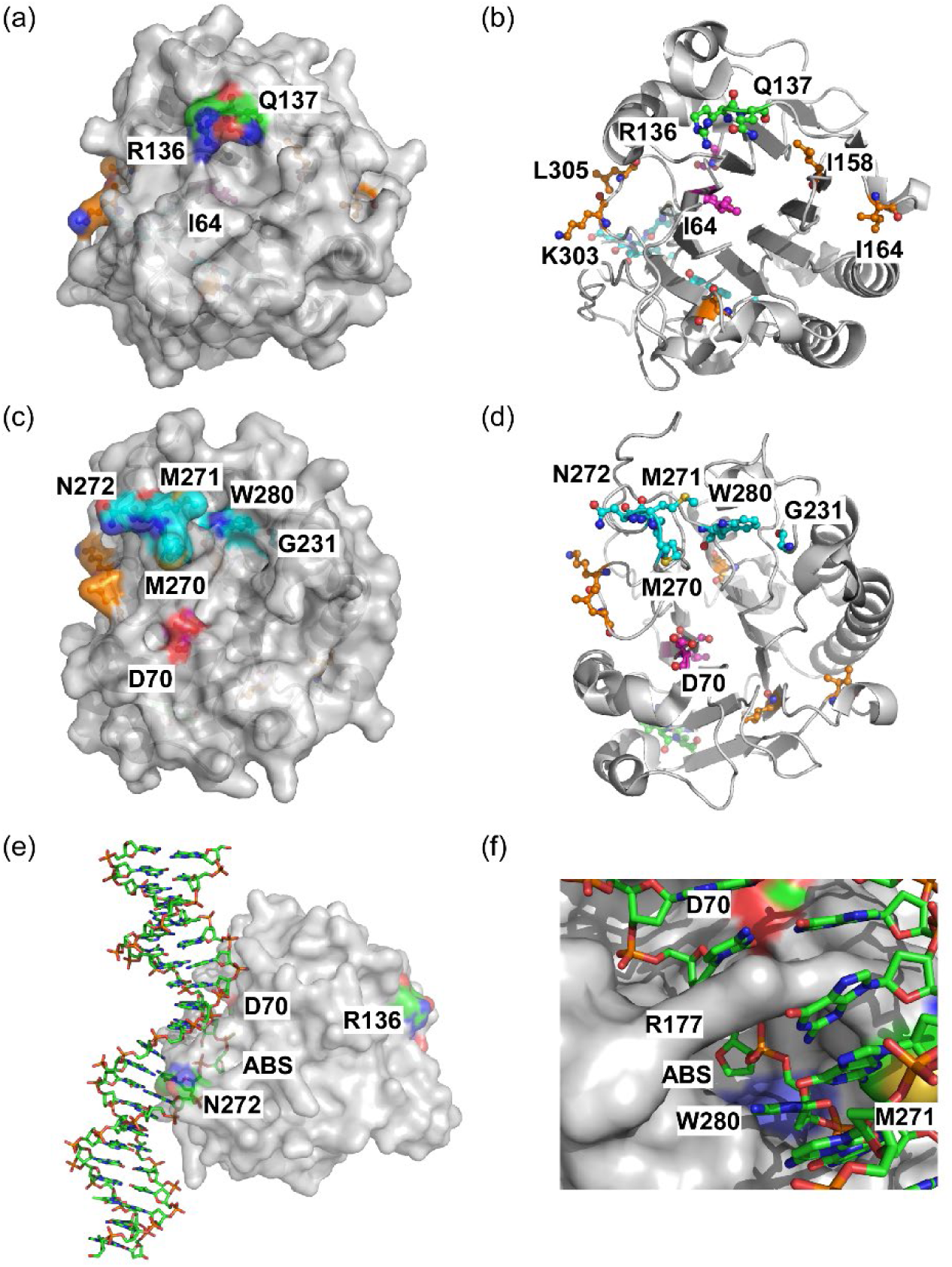
Residues with CSPs greater than 0.02 are shown mapped on the structure of APE1 (PDB ID: 4QHD) for the single point, 10-fold molar excess of APX3330 dissolved in acetonitrile. APE1 is shown as a semi-transparent gray surface rendering in (a) and cartoon rendering with residues as ball-and-stick models (C, green for small pocket, cyan for endonuclease active site, magenta for internal residues on N-terminal strand, or orange for other residues; O, red; N, blue; and S, yellow) in (b). In (a), a small pocket is defined by interactions of Ile64 (C, magenta), Arg136, and Gln137 with APX3330 (C, green). This pocket is located on the opposite face of APE1 from the endonuclease active site. (b) The same as in (a) is shown as a cartoon rendering. (c) A semi-transparent surface rendering for a view of APE1 rotated approximately 180° highlights residues within the endonuclease active site (C, cyan) with significant CSPs. (d) A cartoon rendering of the same view as in (c) is shown. (e) A surface rendering of APE1 with a stick model of bound DNA substrate mimic (PDB ID: 5DFI) is shown. Highlighted in the color scheme indicated above are a subset of residues with significant CSPs that are found in either active site (Asp 70 and Asn 272) or the small pocket (Arg136) and the abasic site (ABS). (f) A close-up of the endonuclease active site is shown with the abasic site (ABS) flipped out into the shallow pocket defined by Trp280 (light blue).

To map protons within APX3330 that are affected by its interaction with APE1, we explored ligand-based waterLOGSY experiments for APX3330 dissolved in ethanol-D6 or acetonitrile-D3. The waterLOGSY experiment examines solvent accessibility of ligands bound to proteins through magnetization transfer from the water through the protein to the ligand. Water molecules bound to the protein, those experiencing chemical exchange with protein labile protons, or water molecules involved in the protein-ligand complex contribute to the magnetization transfer (13; 12; 27; 53). The differences in waterLOGSY peak intensities obtained for the ligand in the presence and absence of the protein indicate an interaction. As previously noted, the difference is dependent on the conditions used including the ratio of protein to ligand (12). In acetonitrile, methylene 3, 5-11, and methyl 12 protons within the nonyl hydrophobic tail, methyl 4 protons on the quinone ring, and the olefinic proton 1 (Fig. 1) exhibit peaks in the waterLOGSY difference spectrum (SI Fig. S10) suggesting that binding involves the hydrophobic tail of APX3330. In ethanol, smaller difference peaks were observed in the waterLOGSY spectrum for the same protons (SI Fig. S10).

We modeled APX3330 bound to the small pocket and to endonuclease active site using molecular docking, taking into consideration information obtained from the waterLOGSY as well as the ^1^H-^15^N HSQC analyses. Using both CDOCKER and AutoDock Vina programs,(65; 59), a consensus pose for APX3330 bound to the small pocket was identified (Fig. 5). In this pose, the hydrophobic tail (9 carbons) of APX3330 extends into the pocket to sit atop Leu62 and Ile91, while the methoxy and carbonyl oxygens of the dimethoxyquinone ring form hydrogen bonds to Arg136 and Gln137. When aligned with the crystal structure of CRT0044876,^19^ many atoms in APX3330 overlap well with atoms in the indole ring of CRT0044876, and the carboxylic acid group in APX3330 sits near a water molecule in the crystal structure and within hydrogen bonding distance of Thr61. In this manner, APX3330 contacts residues Arg136 and Gln137 that display high CSPs, as well as potential contacts to β-strands that comprise parts of the N- and C-termini of APE1, respectively.

**Figure 5.**
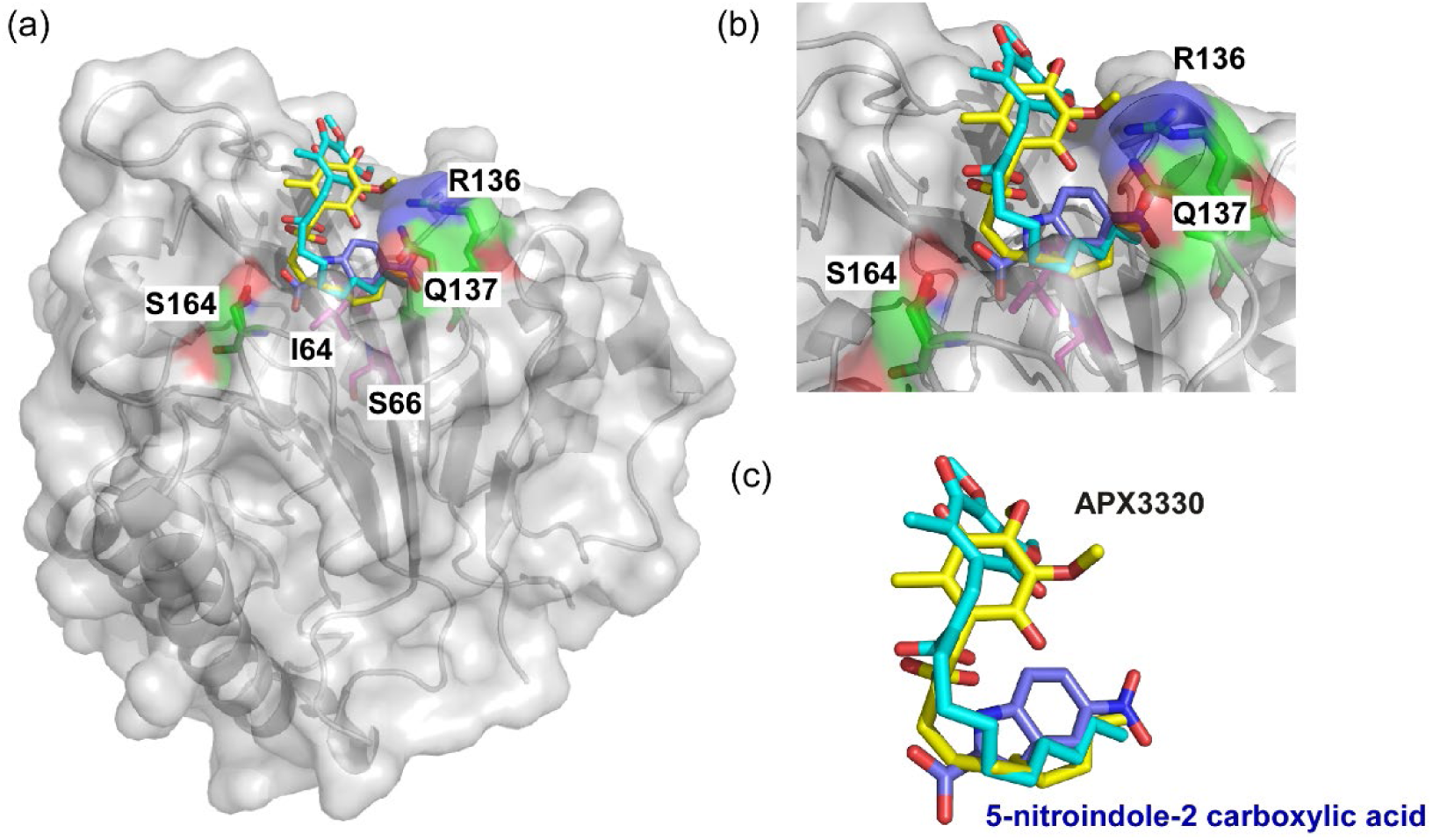
Docking of APX3330 based on crystal structure of CRT0044876 (7TC2). (a) A semi-transparent surface of APE1 (4QHD) is shown with residues Arg136, Gln137, and Ser164 (stick model with C, green, O red, and N blue), which define the small pocket identified in the crystal structure and our CSP analysis. Ser164 exhibited a significant CSP in the 10-fold molar excess point in our titration experiment for APX3330 dissolved in acetonitrile. For reference, Ile64 and Ser66 are shown as stick models with C atoms in magenta. Consensus poses for APX3330 obtained from AutoDock Vina in yellow and CDOCKER in cyan are shown superimposed on the pose for CRT0004876 from the crystal structure. (b) A closeup view of the consensus poses shown in (a). (c) The consensus APX3330 models shown in yellow and cyan as indicated in (a) with CRT0044876 in blue.

Modeling of APX3330 to the endonuclease active site of APE1 was done using AutoDock Vina and AutoDock 4.2(59; 47; 19). One of several comparable low energy poses for APX3330 bound in the endonuclease active site pocket 1 (SI Fig. S11) is similar to the single pose that was previously reported for E3330 bound to the endonuclease active site (43). However, several alternative poses were also identified including those that expand into a second potential binding pocket adjacent to the endonuclease active site (pocket 2). At this site, the nonyl tail can contact residues Met270, Met271, Asn272, and Ala273, which showed shifted CSPs above, and the dimethoxyquinone ring sits adjacent to W280 and M270. The modeling results suggest that there are potentially many poses of APX3330 bound to or near the endonuclease active site that would be consistent with the observed CSPs and waterLOGSY results (SI Fig. S11).

### Effects of APX3330 co-solvent effects on APE1 endonuclease activity

Due to the limited solubility of APX3330 in water, it is necessary to use a co-solvent. For interactions of APX3330 with [*U*-^15^N] APE1, the co-solvent clearly has a significant effect on the nature of the interactions including binding to either the endonuclease active site alone with ethanol as a co-solvent or binding to the endonuclease active site and the small pocket with acetonitrile as the co-solvent.

Small molecules tested for inhibition of APE1 endonuclease activity are typically dissolved in DMSO (final concentration 0.1-0.2%). In this case given the striking differences for acetonitrile versus ethanol as a co-solvent in interactions of APX3330 with [*U*-^15^N] APE1, it was of interest to determine the effects of co-solvent on inhibition of endonuclease activity by APX3330. The reactions were started by addition of Mg^2+^ using an injector and synchronous plate reading mode on the BioTek Synergy Neo2 instrument in a modified version of a previously reported assay (3). Briefly, a kinetic fluorescent signal from the fluorescein-labeled product results from cleavage of one strand of DNA including tetrahydrofuran as a substrate mimic for an abasic site in a duplex substrate. Prior to cleavage, fluorescence from fluorescein on the 5’ end of the strand including the AP site mimic is quenched by the presence of dabcyl on the 3’ end of the complementary strand.

In the first set of measurements, APX3330 was added to reactions including APE1 and substrate DNA. In this case, no significant reduction in the steady state reaction rate for APE1 (final concentration 0.6 nM) endonuclease activity was observed for addition of 10 μM APX3330 dissolved in acetonitrile, ethanol, or DMSO (final concentrations of 0.2%) with reaction rates of 80, 77, and 74% of the control reaction, respectively. However, if APX3330 (50 μM) dissolved in acetonitrile, ethanol, or DMSO is preincubated with APE1 (3 nM) prior to addition of substrate DNA, the reaction rate for APX3330 dissolved in ethanol is 32% of the control rate, while reaction rates were 87% for acetonitrile, and 110% for DMSO (final concentrations 0.6 nM APE1, 10 uM APX3330, 0.1% solvent) (Fig. 6). For acetonitrile or DMSO as a co-solvent, the endonuclease activity is not significantly increased or decreased when preincubated with APE1.

**Figure 6.**
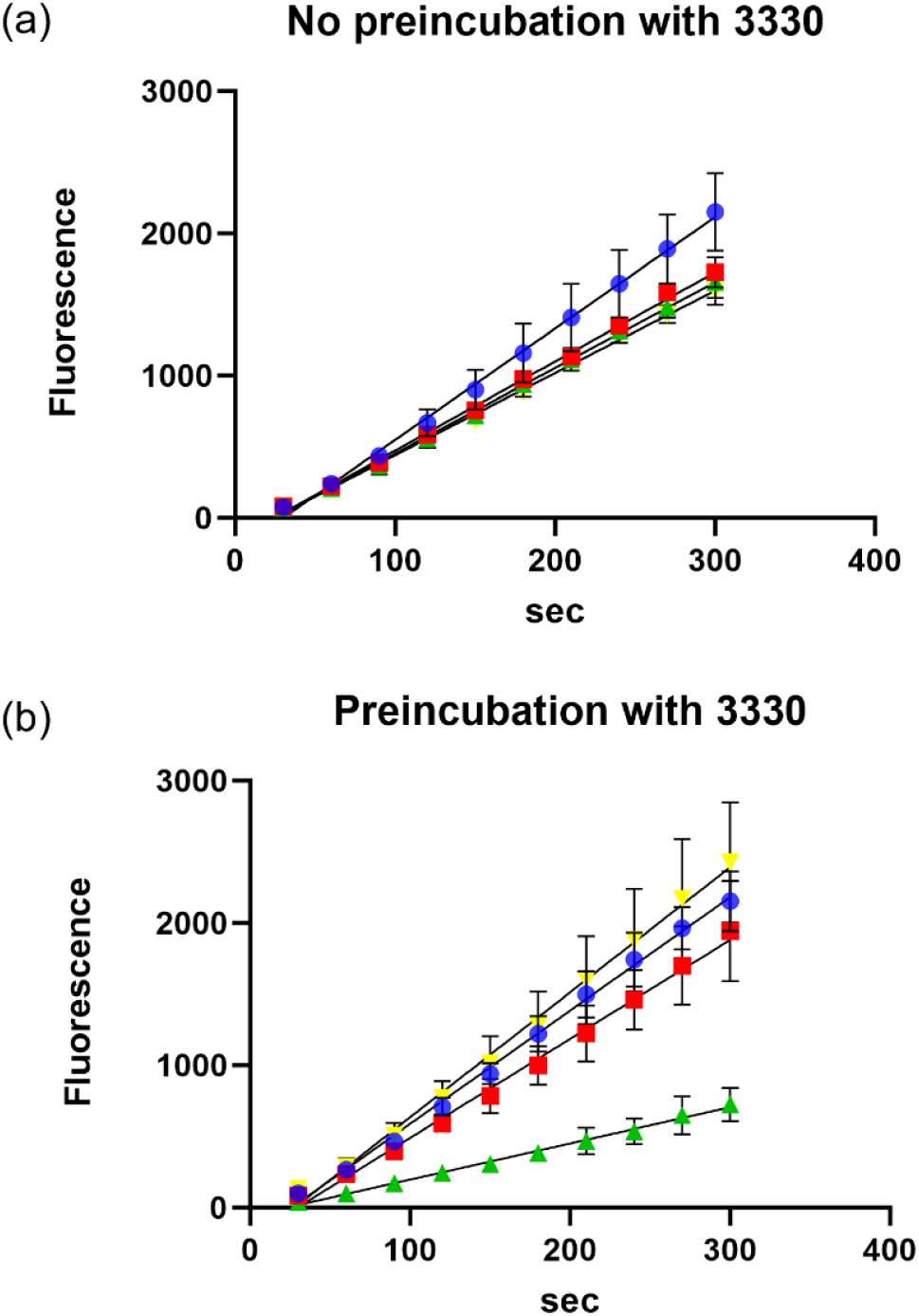
APX3330 inhibits endonuclease activity only when preincubated with APE1 and when ethanol is used as a co-solvent. (a) APX3330 (10 μM) dissolved in acetonitrile (red square), ethanol (green triangle), or DMSO (yellow cross) has little effect on the rate of endonuclease activity (control blue circle) when added to reactions containing APE1 (0.6 nM) and DNA substrate (50 nM). (b) Preincubation of APX3330 (final concentration of 10 μM) dissolved in acetonitrile, ethanol, or DMSO (same symbols and colors as for (a)) shows differential effects, with the co-solvent ethanol resulting in a significant reduction in the rate of endonuclease activity. Triplicate measurements were made for endonuclease activity.

### The melting temperature of APE1 is not sensitive to co-solvent

In past studies, DMSO was used as a co-solvent to determine the effect of APX3330 on melting temperature and endonuclease activity of APE1. As previously reported, the melting temperature measured by CD decreased by 2 °C (67). The melting temperature of APE1 in the presence of acetonitrile alone is 0.5 °C lower and 1 °C lower in ethanol as compared to the control with a melting temperature of 47.3 °C. APX3330 dissolved in either ethanol or acetonitrile lowered the melting temperature of APE1 by 2 and 2.5 °C, respectively (SI Fig. S12) at a 10-fold molar excess as compared to the control with the same solvent, consistent with results obtained using DMSO as a co-solvent. Thus, with increased temperature, effects of APX3330 on the stability of APE1 are similar independent of the co-solvent used in the experiment.

### Prolonged exposure of APX3330/acetonitrile with APE1 results in loss of chemical shifts

Using a single [*U*-^15^N] APE1 sample, we acquired ^1^H-^15^N HSQC spectra following additions of APX3330 dissolved in acetonitrile to yield a 1, 2, 4, 6, 10, and 12-fold molar excess (Fig. 7). We observed significant CSPs for the 10-fold molar excess sample. CSPs above a threshold of 0.02 in this experiment include the following residues: Ile64, Ser66, Asp70, Arg136, Gln137, Ile146, Ser164, Ala175, Gly231, Ser252, Tyr264, Met271, Asn272, and Ser307 (SI Fig. S13). There is significant overlap of the CSPs mapped in this experiment buffered in the absence of added DTT or EDTA with those mapped in the single 10-fold molar excess point experiment (Fig. 3), although we note that CSPs are larger for the single point experiment. An exception is the absence of significant CSPs for Gly279 and Trp280. Based on selected CSPs, we determined a K_d_ of 0.5 mM for binding of APX3330 to APE1 (SI Fig. S14), similar to the previously reported values of 0.4-0.6 mM (43). In contrast to the previous study, the ^1^H-^15^N HSQC spectrum for the 12-fold molar excess in our titration was collected after 24 hours incubation and revealed a significant loss in the number of backbone resonances (Fig. 7). This result was consistent with NEM-footprinting mass spectrometry experiments that we had previously conducted in which prolonged exposure of APE1 with APX3330 resulted in partial unfolding of the protein (57).

**Figure 7.**
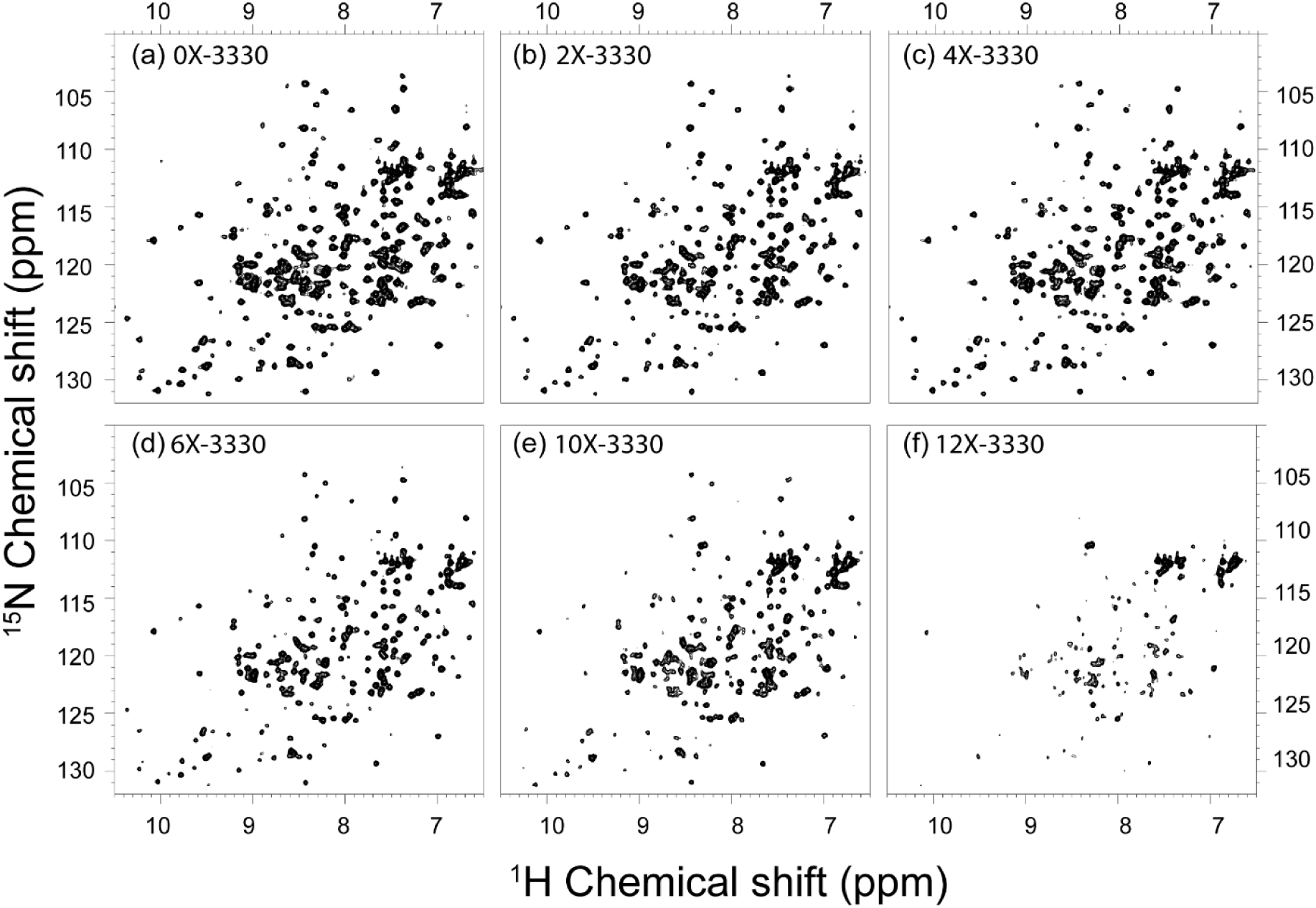
Concentration dependent effects of APX3330 on [*U*-^15^N]-APE1. All ^1^H-^15^N HSQC spectra were recorded on a single APE1 sample. (a) 2D ^1^H-^15^N HSQC spectra of (85 μM) free protein. ^1^H-^15^N HSQC spectra with increasing APX3330 concentration: (b) 2-fold APX3330, (c) 4-fold APX3330, (d) 6-fold APX3330, (e) 10-fold APX3330, and (f) 12-fold APX3330. It took a total of 24 hours to record all spectra. The time points for APX3330 titration are as follows: (b) 2.0 hours, (c) 4.2 hours, (d) 6.3 hours, (e) 8.3 hours, and (f) 24 hours at 25 °C.

Acquisition of the ^1^H-^15^N HSQC spectra for the APX3330 titration of [*U*-^15^N]-APE1 (approximately 2-3 hours per spectrum) necessarily imposes time *and* concentration effects in the experiment. To investigate the time component, we incubated [*U*-^15^N]-APE1 with an 8-fold molar excess of APX3330 as an intermediate concentration between 6-fold, which had few CSPs, and 10-fold molar excess, which had substantially more CSPs, and performed the study at 30 °C (Fig. 8). In this experiment, we observed a clear loss of backbone resonances after incubation longer than 20 hours.

**Figure 8.**
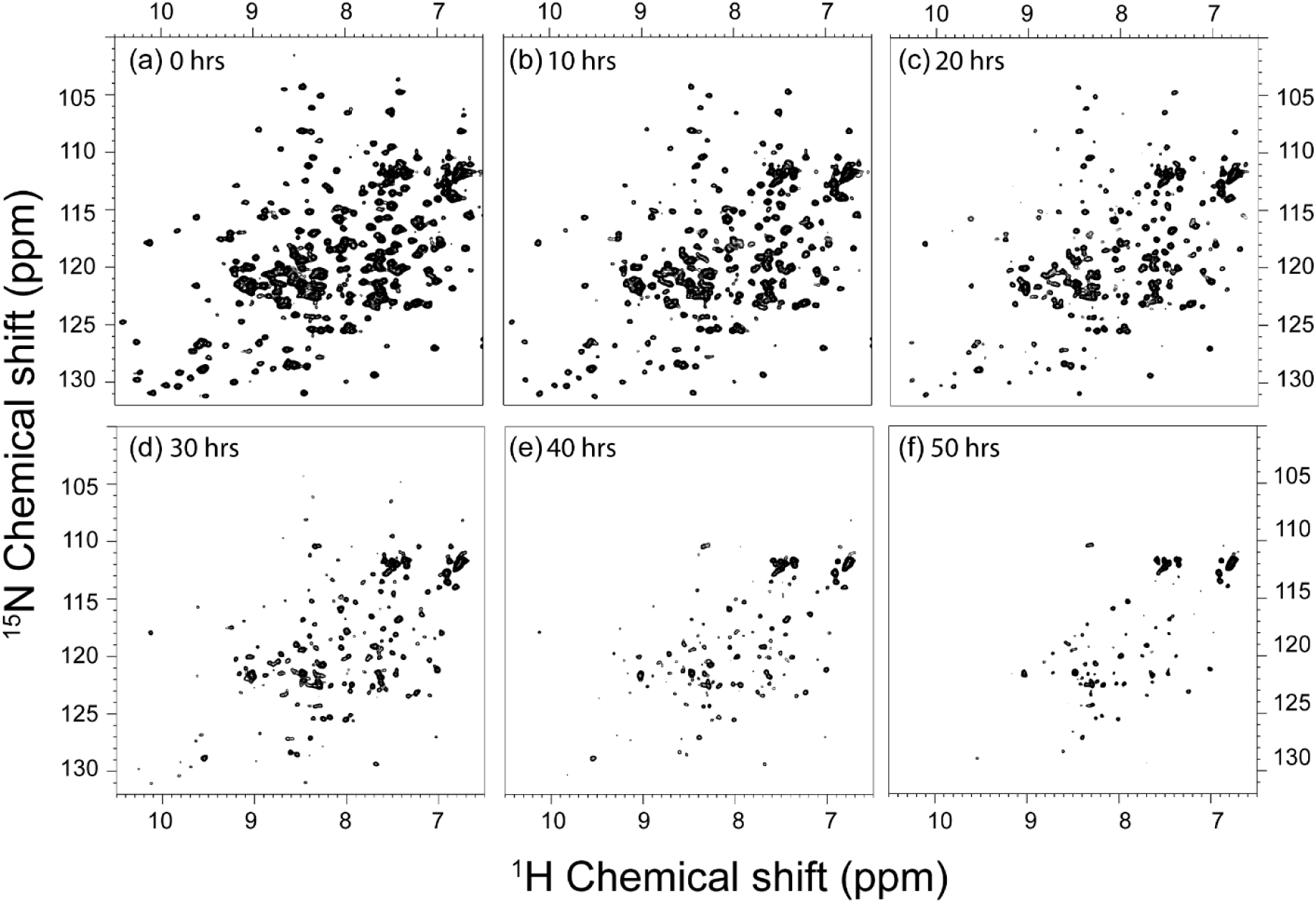
Time dependent effects of APX3330 on [*U*-^15^N]-APE1. A time course of ^1^H-^15^N HSQC spectra were collected at 30 °C following addition of an 8-fold molar excess of APX3330 at (a) 0 hours, (b) 10 hours, (c) 20 hours, (d) 30 hours, (e) 40 hours, and (f) 50 hours. The peak intensities decrease as time progresses and approximately 35% of the backbone resonances are lost after 40 hours; this spectrum appears similar to that of the 12-fold APX3330.

For comparison, we acquired ^1^H-^15^N HSQC spectra for [*U*-^15^N]-APE1 samples with a 10-fold molar excess of APX3330 dissolved in ethanol-D6 over time using the buffer containing DTT and EDTA as previously reported. There was no evidence of unfolding for the ethanol experiment (SI Fig. S15). However, in attempts to prepare a sample with a 12-fold molar excess of APX3330 for direct comparison with the previously study, it was necessary to reduce the protein concentration to approximately 75 μM as previously reported (43). At 100 μM APE1, addition of a 12-fold molar excess of APX3330 resulted in precipitation of the protein and a ^1^H-^15^N HSQC spectrum consistent with random coil (SI Fig. S16). While use of a concentration of 76 μM protein for the 12-fold molar excess of APX3330 was indicated in the previous report, there was no mention of precipitation at higher concentrations of protein.

To determine whether acetonitrile might contribute to the observed loss of chemical shifts in our time-dependent experiment with an 8-fold molar excess of APX3330, we measured ^1^H-^15^N HSQC data for APE1 in the presence of acetonitrile-D3. Over a time period of 15 days, there was no loss of chemical shifts. However, after 48 hours, we observed CSPs for a few residues, notably R136 and Q137 within the small pocket (SI Fig. S17). Using acetonitrile as a co-solvent, the redox inhibitor APX2009, which binds to the same small pocket as APX3330 as reported (48), exhibited no loss of chemical shifts over time (SI Fig. S18) suggesting that loss of chemical shifts is dependent upon the presence of APX3330.

Assignment of the resonances remaining in the ^1^H-^15^N HSQC spectrum for the 40 h time point revealed a loss of 81 backbone resonances (∼35% of the assigned resonances). These residues were mapped on the structure of APE1 along with regions that were not assigned in the native spectrum (Fig. 9a, b). APE1 has a core beta sandwich structure comprising a six-stranded beta sheet (sheet 1) and a five-stranded beta sheet (sheet 2). The N-terminal residues (39–62), which lack secondary structure, lead to a beta strand including residues 62-68 (strand 1) located within sheet 1 (Fig. 9c, d). The C-terminal region of the protein comprises strand 6 (residues 312-316), which is adjacent to strand 1 arising from the N-terminal region of the protein. Backbone resonances lost upon treatment with APX3330 include most of the residues within strands 1 (63–68) and 6 (312–314) along with residues in adjacent strands 2 (92–93) and 5 (299–301) all within sheet 1. Other secondary structural elements for which backbone resonances were lost include alpha helices 2, 4, 5, and 6. Among these, alpha helix 2 also lacks backbone resonance assignments for much of the helix. The remaining residues for which backbone resonances were lost in a time-dependent manner reside within loops.

**Figure 9.**
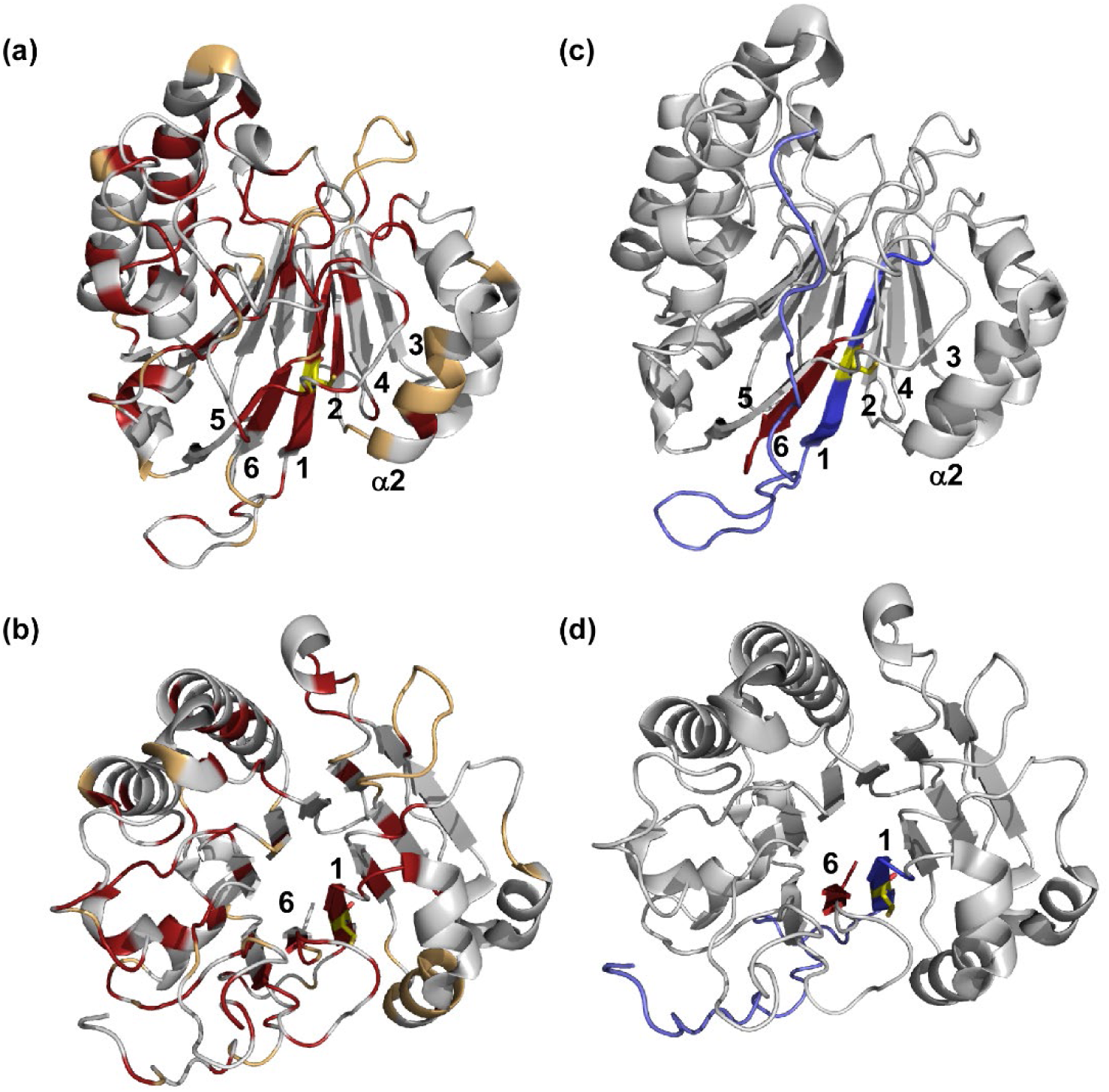
Time-dependent effects of APX3330 on APE1. (a) Highlighted in red on a cartoon rendering of APE1 (4QHD) are the residues for which backbone resonances were lost in the 40 hour time point in the HSQC spectrum, while in tan are the residues that were initially unassigned. Cys65 located in strand 1 is shown as a yellow stick model. (b) A view of APE1 with the same residues highlighted looking down the beta sandwich. Numbering is shown for the strands comprising sheet 1 of the beta sandwich core. In (c) and (d), the same views of APE1 are shown with the N-terminal region highlighted in blue including strand 1 and the C-terminal region in red, including strand 6, highlighting the juxtaposition of the N- and C-terminal regions of the protein within sheet 1.

Consistent with the NMR experiments, treatment of APE1 with APX3330/acetonitrile revealed a time-dependent loss of secondary structure in CD experiments. Far-UV spectra for (260-190 nm) were measured at 25 and 30 °C. Prominent features include negative bands at 210 nm and 230 nm. The spectra for untreated APE1 are stable up to 5 days at 25 °C and 3 days at 30 °C, while treatment with APX3330 results in a loss in the intensity and a shift in the negative band at 210 nm and complete loss of the negative band at 230 nm (SI Fig. S19). These results are consistent with the time-dependent loss of backbone resonances for some residues found in secondary structural elements in the NMR experiments (Fig. 9).

### APE1 does not aggregate in the presence of APX3330

We next considered potential underlying causes for the loss of chemical shifts in the ^1^H-^15^N HSQC spectrum. One possibility is that the protein is aggregating in the presence of APX3330, which could potentially introduce heterogeneity in the environment of the amide nuclei (8). To address this concern, the effects of APX3330/acetonitrile on APE1 were analyzed by diffusion ordered spectroscopy (DOSY) NMR and size exclusion chromatography (SEC). In the DOSY experiment, we determined the diffusion coefficient of APE1 alone and then following 2 days incubation with an 8-fold molar excess of APX3330. The diffusion coefficient calculated for APE1 alone was 1.123 x 10^-10^ m^2^/s, which corresponds to a molecular weight of 32.4 kDa (SI Fig. S20. With an 8-fold molar excess of APX3330, the diffusion coefficient was 1.124 x 10^-10^ m^2^/s, which corresponds to a molecular weight of 31 kDa. As the calculated molecular weight of APE1 used in these experiments is 32 kDa, we conclude that APE1 remains monomeric and does not dimerize through disulfide bond formation or otherwise aggregate to form large multimers following exposure to APX3330 over time. To determine whether large higher order aggregates that would not be detected by DOSY formed in the course of the incubation with APX3330, we assessed the integral area for the DOSY signal; the signal is reduced ∼20% in signal intensity following APX3330 incubation suggesting that most of the sample remains monomeric (SI Fig. S20). In the characterization by SEC, we observed a large peak for APE1 incubated with APX3330/acetonitrile that eluted at the same volume as the control APE1 sample (SI Fig. S21). Thus, we conclude that APE1 does not dimerize through disulfide bond formation or otherwise aggregate to form large multimers following exposure to APX3330 over time.

Another possibility is that dynamical changes within specific regions of the protein have resulted in the loss of amide cross peaks (8). Although in theory, partial unfolding could lead to the appearance of new resonances consistent with random coil for locally unfolded regions, this would be dependent on the motional timescale. The absence of new resonances is consistent with motions that are on a different timescale and not observable under the conditions of our experiment. There are two criteria for assessing interaction of a compound with a protein by NMR. One is loss of peak intensity with increasing concentrations. The second is CSPs. We observed both in the concentration dependent experiment in which we titrated APE1 with increasing concentrations of APX3330. Peak intensities for each concentration of APX3330 in the titration experiment gradually decrease with increasing concentration of APX3330 (SI Fig. S22). Variations in peak intensity are more pronounced than chemical shift perturbations showing both local and global effects. Specifically, peak intensities for residues within the small binding pocket identified for APX3330 have decreased intensities relative to neighboring residues consistent with the proposed binding site on APE1. These lowered intensities persist throughout the course of the titration while lowered peak intensities associated with binding to the endonuclease active site do not (SI Fig. 22). We suggest that global loss of intensities is potentially consistent with an alteration to the dynamical properties of APE1 in the presence of APX3330.

### Loss of APE1 chemical shifts is reversible

Having established that treatment with APX3330/acetonitrile results in a loss of amide cross peaks in the ^1^H-^15^N HSQC spectrum of APE1, we next addressed the issue of reversibility using three different approaches. In the first approach, we dialyzed the APE1-APX33330 sample to remove APX3330; dialysis was performed against a volume 1000 times that of the sample, which diluted the concentration of APX3330 to approximately 0.7 μM. No loss of APE1 was evident following dialysis as measured by UV; however, loss of color in the sample suggested that APX3330 was removed. APX3330 in powder form is bright orange; samples including an 8-fold molar excess are yellow orange in color. Following dialysis of APX3330 from the APE1-APX3330 sample, a small subset of amide resonances reappeared after 48 hours, and a few more after 96 hours in the ^1^H-^15^N HSQC spectrum (Fig. 10). This result is consistent with a very slow refolding process.

**Figure 10.**
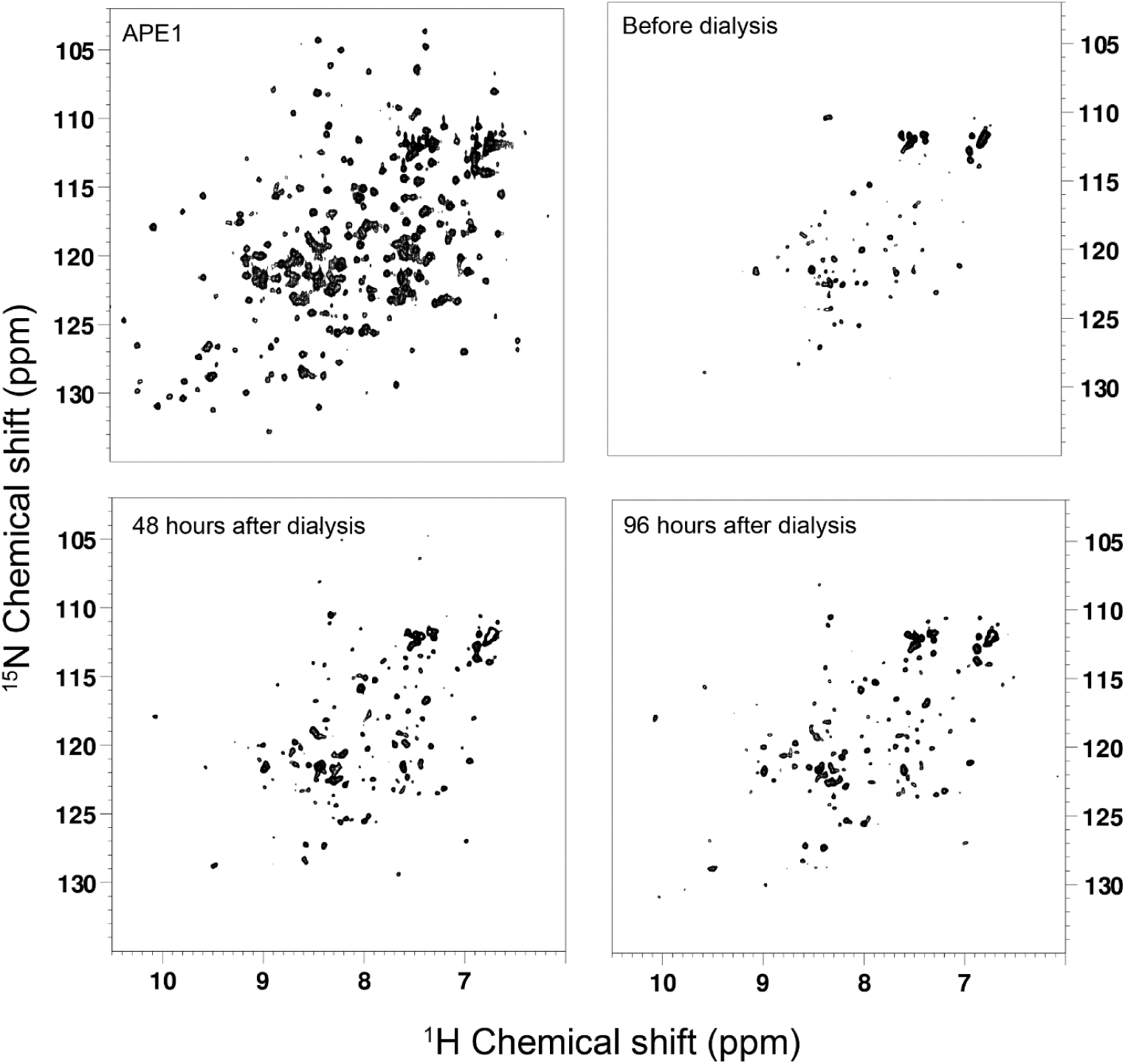
Chemically induced partial unfolding of APE1 is reversible in the presence of DNA following removal of APX3330. (a) Following a 2 day incubation with APX3330 at an 8-fold molar excess for which the ^1^H-^15^N HSQC spectrum of [*U*-^15^N]-APE1 is shown (black), DNA substrate mimic was added at a stochiometry of 1:1 (DNA:protein) (red); only subtle changes are observed. (b) The ^1^H-^15^N HSQC spectrum of [*U*-^15^N]-APE1 treated with APX3330, followed by addition of DNA, and then dialysis to remove APX3330 is shown in black. For comparison, the ^1^H-^15^N HSQC spectrum of APE1-DNA (1:1) complex is shown in red (close-up regions shown in SI Fig. S17). The comparison suggests that APE1 interacts with DNA following removal of APX3330 by dialysis, which in turn implies that APE1 is folded as the ^1^H-^15^N HSQC spectra obtained is similar to that of the APE1-DNA complex.

In the second approach, we tested the effect of adding a stoichiometric amount of DNA substrate mimic (15-mer duplex containing a tetrahydrofuran mimic of an abasic site) on the ^1^H-^15^N HSQC spectrum of APE1-APX3330. As a control for this experiment, we first established the effect of adding duplex DNA to [*U*-15N]-APE1. The ^1^H-^15^N HSQC spectrum was well-resolved and consistent with folded APE1 as expected based on crystallographic studies of APE1 bound to DNA (46; 20) (SI Fig. S23). A K_D_ of 14 μM for binding of the 15-mer duplex DNA substrate mimic to APE1 was determined by fitting the CSPs as a function of increasing DNA concentrations (SI Fig. S24). Consistent with specific binding to APE1, in the 1D ^1^H NMR spectra of the DNA, the imino resonances of the DNA duplex containing the target site exhibit large changes in the chemical shifts (SI Fig. S24). These differences are not only visible in the imino resonances but also in the aliphatic spectra regions. In the APE1 complex with substrate DNA containing a tetrahydrofuran abasic site mimic,(46) Arg177 and Met270 intercalate within the DNA stabilizing a bent conformation. The aliphatic region clearly shows upfield shifts of the methyl resonances consistent with protein-DNA interactions (SI Fig. S24). While it was possible to assign some of the ^1^H-^15^N HSQC cross peaks in the DNA titration experiment by visual inspection, many of the perturbations are too large to confidently assign without additional triple resonances experiments that are outside the scope of this study.

Addition of DNA to the APE1-APX3330 sample produced some changes in the ^1^H-^15^N HSQC spectrum of APE1 but failed to produce a spectrum equivalent to that obtained for APE1 with DNA (SI Fig. S25). In a third approach, we dialyzed the sample containing [*U*-^15^N]-APE1, APX3330, and DNA to remove APX3330 (as indicated by color loss) and obtained a well-resolved ^1^H-^15^N HSQC spectrum similar to that obtained for APE1 complexed with DNA. We estimate that at least 40% of APE1 is complexed to DNA in this sample (SI Fig. S25, S26).

The crystal structures of APE1 alone or bound to DNA are very similar with a rmsd of 0.5 Å. While side chain conformations are altered for some of the residues, there are no large conformational changes to the backbone of the protein upon binding DNA (46). Thus, we suggest that loss of ^1^H-^15^N HSQC resonances is induced by incubation with an 8-fold molar excess of APX3330 and results in partial unfolding of APE1 within the regions noted above. In the presence of an 8-fold molar excess of APX3330, APE1 cannot refold and bind DNA. However, once APX3330 is removed, the presence of DNA shifts the equilibrium in solution toward folded APE1 that can bind DNA.

## Discussion

### Partial unfolding required to expose Cys65

Prior to determination of the crystal structure of APE1(22), Cys65 was identified as the nucleophilic Cys residue required for redox activity involving a thiol-exchange reaction (64). Once the structure was available, it was clear that of the seven Cys residues present in APE1, two are solvent accessible Cys99 and Cys138, while the remaining five Cys residues are buried, including Cys65, which is located on a beta strand in the core of the protein. Thus, at least partial unfolding would be required to expose Cys65 for redox activity. Although APE1 is found in all organisms from bacteria to man as base excision repair activity is essential for survival, redox activity is dependent upon the presence of Cys65, which is conserved only in mammals. In zebrafish APE, the residue equivalent to Cys65 is Thr58, and this protein exhibits no redox activity. However, substitution of Thr58 with Cys confers redox activity to zebrafish APE confirming the important role of this protein (21). More recently, the MTS in APE1 was identified as the C-terminal residues (298–318) (35) located within the same beta sheet in the beta sandwich core as Cys65. In order to be transported to the inner mitochondrial space, the MTS must be exposed through partial unfolding of APE1.

At temperatures well below its Tm of ∼ 47 °C, APE1 behaves as a well-folded protein with high resolution crystal structures (24; 52) attesting to its ordered state. NEM footprinting experiments discussed in the Introduction provided the first evidence that APE1 could be partially unfolded following prolonged treatment with APX3330 in the presence of NEM. The present studies provide a molecular basis for understanding partial unfolding by APE1 initiated by separating two adjacent central strands, one comprising the N-terminal residues (62–66) and the second, the C-terminal residues (313–318) within the first β-sheet of the β-sandwich core.

### APX3330 mechanism of action and co-solvent effects

Determining a mechanism of action for APX3330 is complicated by the fact that use of different co-solvents produces different results for ^1^H-^15^N HSQC analysis of interactions with [*U*-^15^N] APE1 along with distinctly different time-dependent effects on the stability of [*U*-^15^N] APE1. Different results were also obtained for endonuclease assays in which APE1 was preincubated with APX3330 in different co-solvents; however, minimal co-solvent effects were apparent for addition of APX3330 added in the presence of DNA substrate mimic. Similarly, decreases in melting temperature were comparable independent of the solvent used. In considering these results, it should be noted that in endonuclease assays the concentration of APE1 used is subnanomolar (0.6 nM) and inhibition was measured for APX3330 at 10 μM (∼ 17,000-fold molar excess) with 50 nM substrate DNA. In NMR experiments, ∼ 80-100 μM [*U*-^15^N] APE1 was used with APX3330 concentrations up to 1.2 mM (a maximum of 12-fold molar excess). Melting temperature experiments were done using 20 μM APE1 with a 10-fold molar excess of APX3330. Further, given the differences in binding affinity measured by NMR for DNA (14 μM) versus APX3330 (500 μM), we would not expect APX3330 to compete effectively with DNA for binding to APE1.

The most likely difference between the co-solvents is desolvation effects on APE1. Although both ethanol and acetonitrile are polar solvents, ethanol can serve as both hydrogen-bond donor and acceptor, while acetonitrile can only serve as an acceptor. Thus, ethanol might be expected to compete more effectively with water molecules in hydrogen-bonding interactions to amino acid residues. Treatment of [*U*-^15^N] APE1 with ethanol-D6 resulted in a significant CSP for Asn174, located in the endonuclease active site pocket and hydrogen-bonded directly to a water molecule. This suggests that ethanol may alter the properties of the endonuclease active site and confer preferred binding to this site. While it is possible to mathematically subtract CSP value obtained for ethanol from those obtained for the complex (43), the biophysical effects of ethanol on APE1 cannot be subtracted. A second residue, whose chemical environment was perturbed by addition of ethanol to [*U*-^15^N] APE1 was Ser66. This residue is a buried residue that forms a water-mediated hydrogen bonding network with Gln95 and Asp283 (SI Fig. 7) that is potentially important for the integrity of the beta sandwich core. Significant CSPs were observed for Ser66 for interactions of APX3330 in acetonitrile with [*U*-^15^N] APE1 and for Asp283 for DMSO interactions.

The most striking difference in the co-solvent effects was observed in the time-dependent experiments with prolonged exposure of APE1 to a 10-fold molar excess of APX3330 in ethanol showing no evidence of loss of chemical shifts. The fact that we observed precipitation and an ^1^H-^15^N HSQC spectrum consistent with an unfolded protein when a 12-fold molar excess of APX3330 was added to 100 μM [*U*-^15^N] APE1 a (SI Fig. S16) suggests that at high enough concentrations, rather than unfolding slowly as was observed in acetonitrile, APE1 unfolds very rapidly. In contrast, use of APX3330 in acetonitrile provided a mechanism by which APE1 could be slowly and reversibly partially unfolded. Melting temperatures for APX3330 in all solvents tested (DMSO, acetonitrile, or DMSO) are lowered by ∼ 2 °C suggesting that APX3330 destabilizes the structure of APE1 rather than stabilizing it as might be expected from binding to the endonuclease active site.

Co-solvent effects were also observed in endonuclease assays in which APX3330 in different solvents was preincubated with APE1. Preincubation of APE1 with APX3330/acetonitrile or DMSO did not significantly reduce activity, while preincubation with APX3330/ethanol reduced activity by ∼70%. This finding is consistent with results previously reported for inhibition of APE1 endonuclease activity by APX3330/ethanol in which activity was reduced by 40% (43). In this latter report, APX3330/ethanol was not preincubated with the protein, but the final concentration of APX3330 was 100 μM with 2% ethanol as compared to 10 μM APX3330 and 0.1% ethanol in our experiment. One possible explanation is that preincubation results in binding of APX3330/ethanol to the endonuclease active site, which reduces binding of DNA substrate. A second explanation is that preincubation of APE1 with APX3330/ethanol results in precipitation of at least part of the protein resulting in decreased activity as was observed upon addition of a 12-fold molar excess of APX3330/ethanol to 100 μM [*U*-^15^N] APE1. Independent of the mechanism, the effect in our assays is specific for preincubation of APE1 with APX3330/ethanol.

In our study, an effect of increased time of incubation with APX3330 was first identified in titration experiments, which were completed over a 24 h period. This phenomenon was then confirmed by treating [*U*-^15^N]-APE1 with an 8-fold molar excess of APX3330 and monitoring the effects over time. CSPs calculated for the titration data and time-dependent loss of backbone resonances are consistent with interactions of APX3330 involving the core of the protein. Residues Ile64, and Ser66 (within strand 1) show significant CSPs and the entire strand (consisting of residues 63-68) exhibits loss of backbone resonances. This strand is the most N-terminal secondary structural element in the APE1 fold. This finding, combined with loss of backbone resonances for the C-terminal residues 312-314 in the adjacent strand 6 of the sheet, suggests a mechanism in which APX3330 contributes to partial unfolding of APE1 through interactions with centrally located strands composed of N- and C-terminal residues in the sheet. Further, this process is likely facilitated by interactions with the small pocket defined by CSPs (Figs. 3-5).

We suggest that binding of APX3330 molecules to the small pocket as shown in Figs. 4 and 5 over time perturb APE1 dynamics that affect the beta sandwich core. The effect is partially reversed approximately 48 h after removal of APX3330 potentially due to very slow refolding but can be substantially restored when a DNA substrate mimic is included during dialysis to remove APX3330 (SI Fig. S25). This finding is consistent with the more favorable state being the fully folded state that binds DNA.

### Implications for APE1 function and future drug development

Our studies provide the first evidence for a mechanism of partial unfolding of APE1 albeit mediated by the presence of APX3330, which is currently in Phase III clinical trials for treatment of diabetic retinopathy and continues to serve as a useful chemical tool in probing functional properties of APE1. Although APE1 remains fully folded and stable for at least 20 days in a phosphate buffered solution, in the presence of APX3330/acetonitrile, it begins to unfold over time. Strands including N-terminal and C-terminal residues in sheet 1 within the core of APE1 are implicated in partial unfolding. The N-terminal strand 1 includes Cys65, and the C-terminal strand 6, includes part of the mitochondrial targeting sequence (MTS) comprising residues 298-318. Alpha helix 2 adjacent to strand 1, which would normally shield Cys65, is also unfolded, as is adjacent strand 2. In the folded state, Cys65 is protected from oxidation. Partial unfolding would expose Cys65 and potentially result in increased intermolecular disulfide bond formation under oxidizing conditions. The most prevalent disulfide bond identified following treatment of APE1 with hydrogen peroxide was between Cys65 and Cys138 involving two APE1 molecules as previously reported (40). As Cys65 serves as the nucleophile, oxidation would affect its role as a redox factor and its role in mitochondrial trafficking (63).

In this study, binding of APX3330 to the small pocket defined by residues Arg136 and Gln137 stabilizes a partially unfolded state of APE1 and may affect mitochondrial trafficking, which under conditions of oxidative stress may impact survival of dividing cells. APE1 is thought to protect cells from oxidative stress potentially through repair of damaged mitochondrial DNA (23; 36) in addition to degrading damaged mitochondrial mRNA (4). APX3330 may alter the interaction of APE1 with mitochondrial transport proteins, Tom20,(35) or other cytoplasmic or mitochondrial chaperones such as Mia40 (5). In classifying redox inhibitors and their respective mechanisms of action, partial unfolding of APE1 is associated with APX3330 but not APX2009. These differences in interactions of redox inhibitors with APE1 may provide insights into the mechanism of action and efficacy of these compounds that guide future drug development.

## Materials and Methods

### Protein expression and purification

APE1 (39-318/C138A) was expressed in *E. coli* and purified as previously described.(24) ^15^N-APE1 (39-318/C138A) was expressed as a His-SUMO-fusion in BL21/Rosetta *E. coli*. 2L minimal media cultures were grown with ^15^N-ammonium chloride as the sole nitrogen source.(30; 56) Cells were grown at 37 °C until the culture reached an OD of 0.6; the cultures were then induced with 0.2 mM IPTG and grown for 4 hours at 37 °C. Cells were pelleted and stored at -80 °C prior to purification.

Cells were resuspended in 50 mM sodium phosphate, 0.3 M sodium chloride, 20 mM imidiazole pH 7.8 and lysed using a microfluidizer M110L. Following centrifugation at 35,000 rpm for 30 minutes, the supernatant was filtered and loaded on two 0.5 ml Ni-NTA (Qiagen) columns (one column/L growth) run at 4 °C. The column was washed with 10 column volumes of buffer and incubated with Ulp1 protease (10 μg/column) in 8 ml per column overnight. The flowthrough was then collected the next day. Labeled and unlabeled APE1 were assessed by SDS-PAGE and by QToF intact mass spectrometric analysis to confirm purity and mass.

### NMR spectroscopy experiments

#### ^1^H-^15^N HSQC experiments

2D ^1^H-^15^N heteronuclear single-quantum coherence (HSQC) spectra were acquired at 25 °C or as indicated on a 600 MHz Bruker AVANCE spectrometer equipped with a 5-mm triple-resonance cryoprobe (^1^H,^13^C, and ^15^N) and z-axis pulsed gradient. All NMR experiments were processed using TOPSPIN and analyzed with CCPN NMR(55). [*U*-^15^N] APE1 at a concentration of 80-90 μM was dialyzed against 20 mM sodium phosphate (pH 6.5), 0.1 M sodium chloride in 90%/10% H_2_O/D_2_O and placed in an NMR tube. The inhibitor APX3330 obtained from Apexian Pharmaceuticals was dissolved in acetonitrile-D3. For the inhibitor experiments, acetonitrile-D3 never exceeded 2% (v/v). The chemical shift perturbations (CSP) (Δδ) were calculated using eq. 1,(43) which reflects the total weighted change in ^1^H and ^15^N chemical shift for a given peak in the 2D spectra:

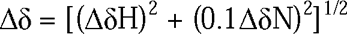

DNA oligonucleotides 5’-GCGTCCFCGACGACG (F indicates tetrahydrofuran serving as a mimic of an AP site) and 5’-CGTCGTCGAGGACGC were purchased from Integrated DNA Technologies, Inc., annealed, and then used without further purification as analysis by ion-exchange chromatography in 10 mM sodium phosphate (pH 8.0) with a KCl gradient indicated that the sample was greater than 95% duplex. DNA duplex was added to a final concentration of 80 μM to form a 1:1 (protein:DNA) complex with the protein along with calcium chloride yielding a final concentration of 90 μM, stoichiometric with the protein and DNA. While APE1 still cleaves DNA in the presence of calcium chloride, it does so much more slowly than with the preferred metal, magnesium (38).

#### WaterLOGSY experiments

WaterLOGSY experiments were conducted at 25 °C in 5 mm diameter NMR tubes with a sample volume of 500 μL. APX3330 was prepared as 50 mM stock solutions in CD_3_CN. Solutions were buffered in 20 mM sodium phosphate (pH .65), 0.1 M NaCl with 90% H_2_O and 10% D_2_O. WaterLOGSY experiments were done with 20 μM APE1 (39-318/C138A) and 200 μM compound for each experiment. The mixing time was set to 1.5 s and data were collected with 128 scans, a sweep width of 10 ppm, an acquisition time of 1.58 s, and a relaxation delay 5 seconds. Prior to Fourier transformation, the data were multiplied with an exponential function with a line broadening of 1 Hz.

#### DOSY experiments

The DOSY experiment was done using unlabled APE1 and APX3330 dissolved in acetonitrile. The ^1^H 2D DOSY experiment utilized a modified Bruker ^1^H 2D DOSY pulse sequence with water suppression to measure diffusion coefficients. NMR data were collected on a Bruker 600-MHz NMR instrument at 30 °C equipped with a triple-gradient TCI room temperature probe capable of generating a gradient field strength of 54 G/cm. The DOSY time interval (Δ) and gradient pulse duration (δ) were set at 600 ms and 1.2 ms, respectively. A total of 16 gradient increments, ranging linearly from 2% to 98%, were acquired with 1024 scans for each increment. Topspin 3.6.2 was employed for NMR data processing. For the diffusion experiments, 400 μM unlabeled APE1 sample was prepared in a buffer 20 mM sodium phosphate (pH 6.5), 0.1 M NaCl with 90% H_2_O, 50 μM DSS, and 10% D_2_O. APX3330 was added in an 8-fold molar excess, 3.2 mM, to the APE1 sample to collect DOSY with the inhibitor.

#### 1H-15N HSQC DNA titration

The DNA titration was done using serial dilutions of a stock DNA solution added to the APE1 protein sample. The final concentration of DNA was 90 μM, which makes protein to DNA ratio 1:1. The final concentrations of DNA in protein solutions ranged from 0 μM to 90 μM and were prepared so that a 9 μL addition of the DNA stock solution (including 1:1 DNA:CaCl_2_) to 470 μL of an APE1 solution resulted in final concentrations of 0, 9, 18, 27, 54, 72, and 90 μM DNA, respectively.

#### Characterization of APX3330 by light scattering

APX3330 diluted from a 50 mM stock solution in acetonitrile and then diluted into buffer was assessed using a refractive index detector (Optilab T-rEX, Wyatt Technology) followed by a multiple light scattering detector (Dawn HeleosII, Wyatt Technology). Each sample injection consisted of 50 μM, 500 μM of sample, and 6 mM decyl maltoside in buffer containing 20 mm sodium phosphate (pH 7.8), 50 mM NaCl. The flow rate was set at 0.5 ml/min and data were collected in a 1-s interval. Data processing and analysis were performed using the ASTRA software (Wyatt Technology). While decyl maltoside produced a peak, no signal was observed for APX3330 in this experiment or by measurement of dynamic light scattering on Malvern Zetasizer Nano-S dynamic light scattering instrument.

#### Modeling of APX3330 interactions with APE1

Structural modeling of APX3330 bound to APE1 in a small binding site near residues Arg136, Gln137, and Ser164 was performed with rigid-receptor docking using AutoDock Vina (ADV) and the CHARMM-based CDOCKER docking algorithm(65; 59). Prior to docking, the apo structure of APE1 was taken from PDBID: 4QHD (24) and prepared with the MolProbity webserver (14; 10) and PROPKA (49) to check for residue flips and determine appropriate protonation states for titratable residues at pH = 7.0. Receptor PDB coordinates were then formatted as a pdbqt file for ADV with the AutoDock Tools program;(2) pdb coordinates for CDOCKER were alternatively formatted with the CHARMM-GUI (31) and energy minimized with 200 steps of steepest decent minimization and the CHARMM36 protein force field (6; 7). APX3330 was then constructed using UCSF Chimera,(51) and subsequently formatted for ADV with AutoDock Tools or for CDOCKER via parameterization with the CHARMM Generalized Force Field (CGenFF)(60–62). Docking was performed in ADV using default settings and a cubic grid of 20 Å/edge centered on the binding site; however, 20 binding modes were generated using an exhaustiveness setting of 100, to account for the many degrees of freedom that could be sampled in the nonyl alkyl tail of APX3330. Similarly, default procedures were followed in CDOCKER (65; 17), which uses a combination of grid-based simulated annealing molecular dynamics and all-atom energy minimization for pose identification and scoring, respectively. A total of 2000 poses were generated with CDOCKER. Manual inspection of the top 10 rank ordered docking poses from both ADV and CDOCKER was preformed to identify the lowest energy consensus structure for APX3330 bound to APE1. Figures were created with PyMOL (1).

Docking of APX3330 to the endonuclease active site of APE1 was done using AutoDock4.2 and AutoDock Vina(59; 47; 19) with an exhaustiveness setting of 50 or 100, respectively. Docking grids for AutoDock 4.2 were created with AutoGrid. All other settings used default options. Two pockets were defined within the endonuclease active site (SI Fig. S11, pocket 1) or adjacent to it (SI Fig. S11, pocket 2) by residues Gly231, Met270, Met271, Asn272, Ala273, and Trp280. Lowest energy consensus structures for APX3330 were determined by manual inspection of the top 20 rank-ordered poses obtained from both programs, including ligand clustering commonly performed for AutoDock 4.2 in AutoDock Tools. Figures were created with PyMOL(1).

#### Circular dichroism experiments

Unlabeled APE1 (39–318) was used for CD experiments. The time dependence of APX3330 on APE1 secondary structure was evaluated by CD. Reactions included 80 μM APE1 (39–318) diluted from a 3 mM stock with either 1.3 % acetonitrile or 640 μM APX3330 (50 mM stock in acetonitrile) buffered in 20 mM sodium phosphate, 0.1 M NaCl pH 6.5 and incubated at either 25 ° or 30 °C. These are the conditions used for the time-dependent NMR experiments. Aliquots were removed from the reactions for CD measurements and diluted to give final concentrations of 20 μM APE1 and 120 μM APX3330. Controls showed that 1.3% acetonitrile had no effect on the CD spectrum of APE1. Data were measured on a Jasco J-1500 circular dichroism instrument with a Peltier temperature control device.

Thermal melting experiments were conducted using 20 μM APE1 buffered in 20 mM sodium phosphate pH 6.5, 100 mM NaCl in the presence or absence of 0.4% ethanol, 0.4% acetonitrile, 200 μM APX3330 dissolved in ethanol, or 200 μM APX3330 dissolved in acetonitrile. The temperature was gradually increased (1 °C/min) from 20 to 75 °C using the Peltier temperature controller on the Jasco J-1500 instrument with mdeg monitored at 210 nm.

#### Endonuclease activity assays

APE1 was diluted from a 3 mM stock solution into 50 mM HEPES pH 7.5, 50 mM NaCl, 0.01% IGEPAL to give final concentrations of 0.2-0.6 nM in the reaction mixtures. Substrate AP-DNA (100 μM) was diluted into the same buffer to yield a final concentration of 50 nM. Substrate DNA, two 30 bp complementary strands (5′**-**6-FAM-GAA-TCC-*CC-ATA-CGT-ATT-ATA-TCC-AAT-TCC -3′ and 5′**-**Q**-**GGA-ATT-GGA-TAT-AAT-ACG-TAT-GGT-GGA-TTC -3′), were obtained from Eurogentec Ltd. (San Diego, CA), with 6-FAM indicating fluorescein, * tetrahydrofuran mimicking an abasic site, and Q dabcyl. Oliogos subjected to HPLC and ion-exchange purification. Prior to cleavage of the tetrahydrofuran containing strand by APE1, dabcyl effectively quenches the fluorescent signal from 6-FAM. Reactions were initiated by injection of MgCl_2_ (5 μL making a final volume of 50 μL per reaction in wells within 384 well plates) resulting in a final concentration of 1 mM, and a kinetic fluorescent signal was measured following excitation at 485 nm and emission at 535 nm on the BioTek Synergy Neo 2 instrument using a synchronous plate reading mode. Reactions for 0.6 nM APE1 produced a linear response over a time period of 5 minutes. Rates were calculated as change in fluorescence over time. Experiments were repeated at least three times with three replicates for each experimental condition.

## Acknowledgements

We acknowledge the Chemical Genomics Core Facility at IU School of Medicine which houses the 600 MHz NMR instrument and supports operational costs. J.Z.V. acknowledges the Indiana University Pervasive Technology Institute for providing supercomputing and storage resources that have contributed to the research results reported within this paper. This work was supported by NIH grants R01CA231267 and R01CA167291 (M.R.K.), R01CA254110 (M.R.K. and M.M.G.), R35GM143054 (J.J.Z.), and P30CA082709 (IU Simon Comprehensive Cancer Center).

