## Supporting Information for "Chemically induced partial unfolding of the multifunctional Apurinic/apyrimidinic endonuclease 1"

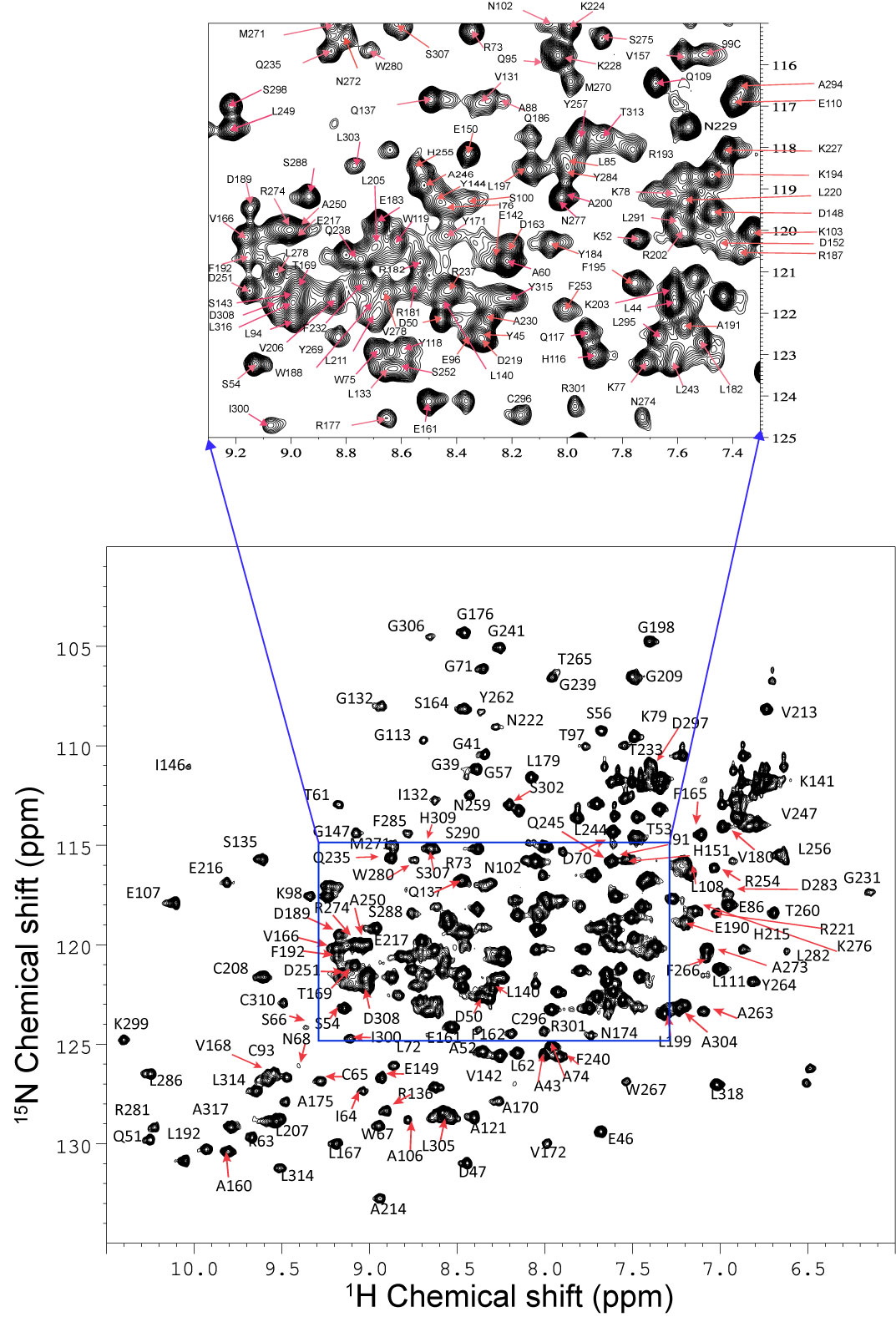

**Figure S1**. 2D ^1^H-^15^N NMR HSQC spectrum of APE1 at 25 °C collected using a 600 MHz Bruker AVANCE spectrometer equipped with a 5-mm triple-resonance cryoprobe. [*U*-^15^N]-APE1 at a concentration of 90 μM was dialyzed against 20 mM sodium phosphate (pH 6.5), 0.1 M sodium chloride in 90%/10% H_2_O/D_2_O and placed in an NMR tube. The 2D ^1^H-^15^N HSQC spectra were acquired with 2048 points in the direct F2 dimension (^1^H) and 256 points in the F1 dimension. (^15^N). Resonances were assigned by visual inspection using BMRB 16516 as a template.

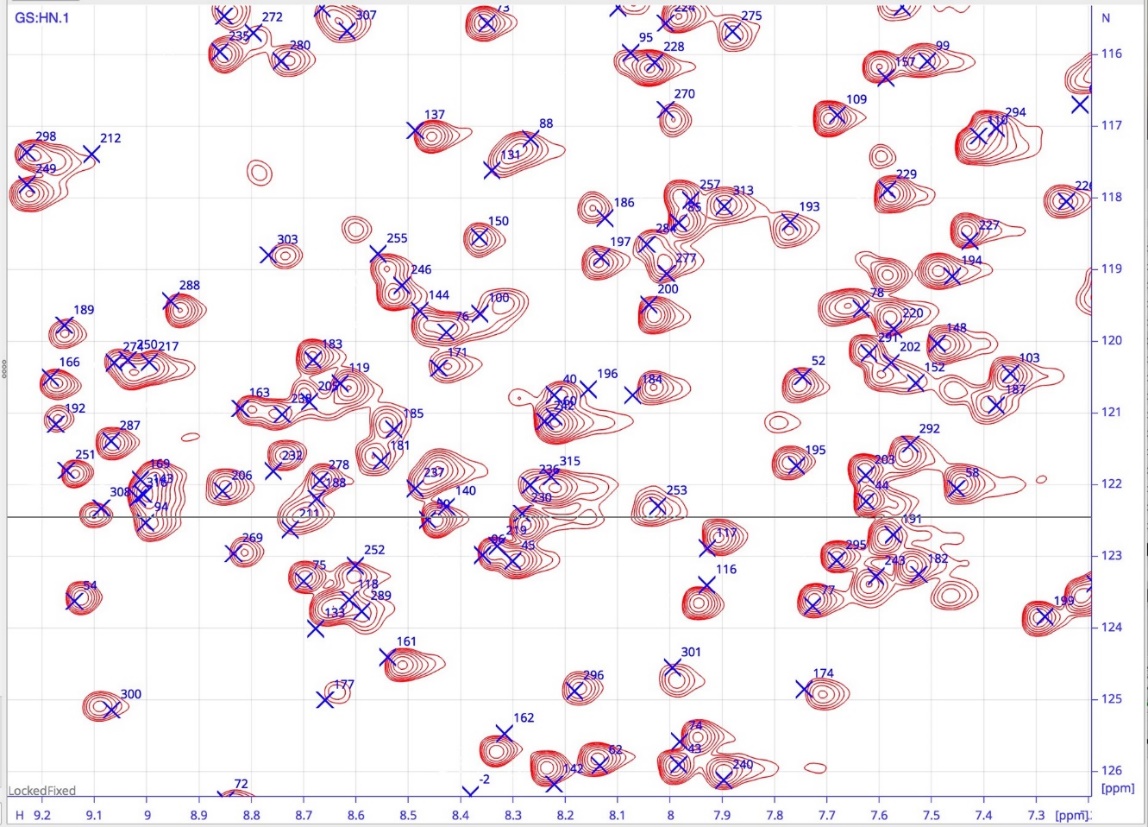

**Figure S2.** Close-up of central portion of the ^1^H-^15^N TROSY spectra collected for [*U*-^15^N] APE1 with BMRB assignments shown as blue crosses. The amide peaks in our spectra are very close to the original assignments.

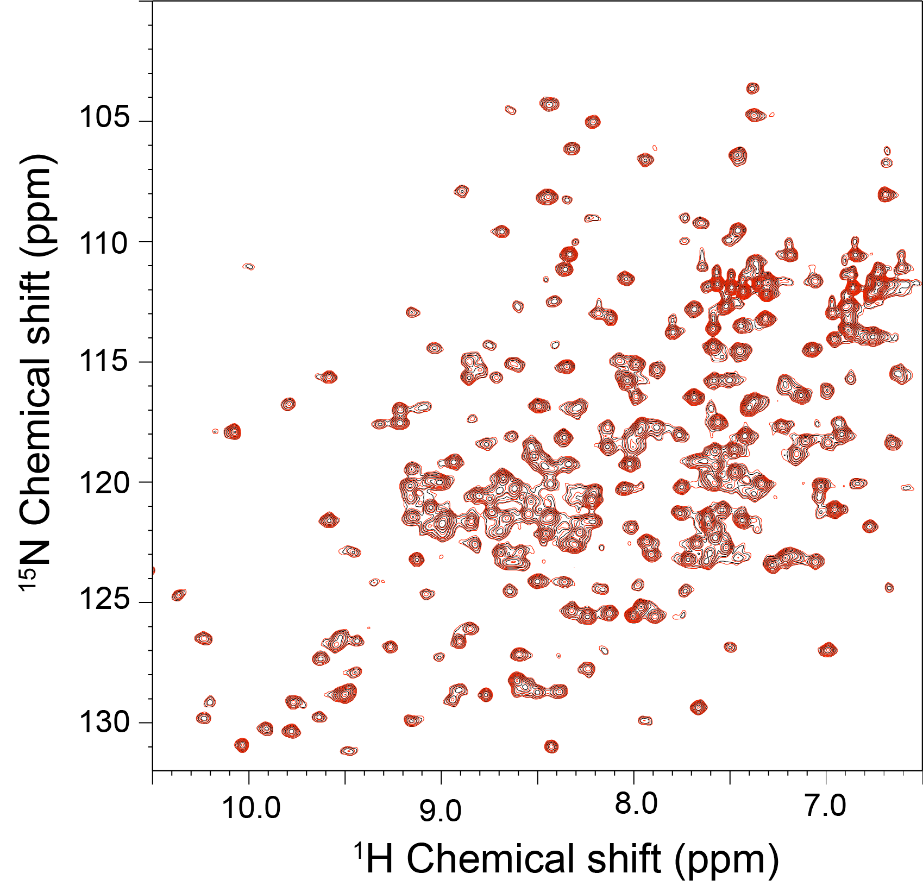

**Figure S3.** 2D ^1^H-^15^N NMR HSQC spectrum of APE1 at 25 °C collected using a 600 MHz Bruker AVANCE spectrometer equipped with a 5-mm triple-resonance cryoprobe. [*U*-^15^N]-APE1 at a concentration of 90 μM was dialyzed against 20 mM sodium phosphate (pH 6.5), 0.1 M sodium chloride, 0.5 mM DTT, 0.2 mM EDTA in 90%/10% H_2_O/D_2_O and placed in an NMR tube. The 2D ^1^H-^15^N HSQC spectra were acquired with 2048 points in the direct F2 dimension (^1^H) and 256 points in the F1 dimension. (^15^N). Resonances were assigned by visual inspection using BMRB 16516 as a template.

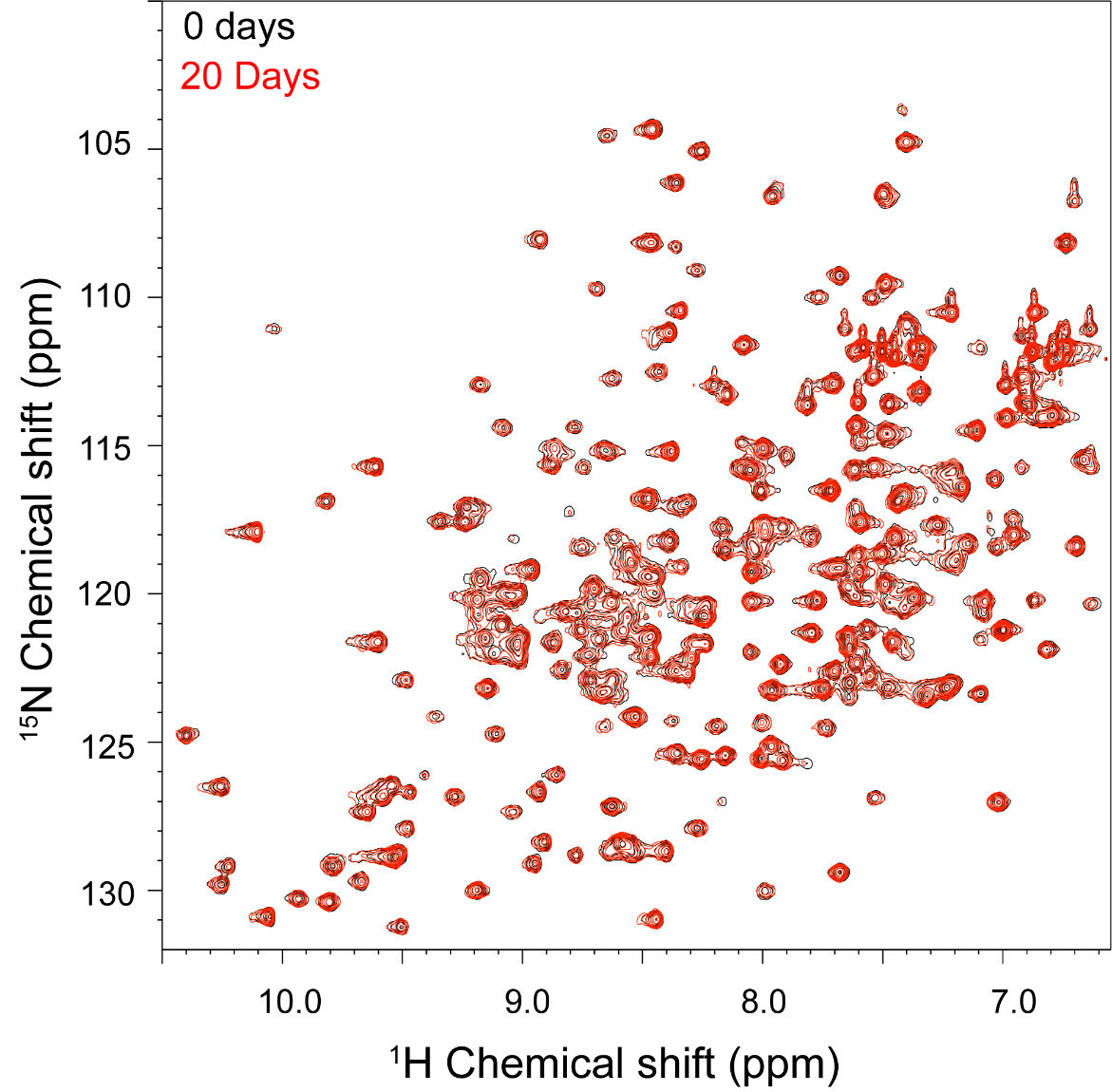
**Figure S4.** APE1is stable for at least 20 days. An overlay of 2D ^1^H-^15^N HSQC spectra was collected on Day 1 (black) and after 20 days (red). The sample was kept at room temperature throughout the experiment. The ^1^H-^15^N HSQC spectrum of [*U*-^15^N]-APE1 was collected using a 600 MHz Bruker AVANCE spectrometer equipped with a 5-mm triple-resonance cryoprobe. [*U*-^15^N]-APE1 at a concentration of 90 μM was dialyzed against 20 mM sodium phosphate (pH 6.5), 0.1 M sodium chloride in 90%/10% H_2_O/D_2_O and placed in an NMR tube. The 2D ^1^H-^15^N NMR HSQC spectra were acquired with 2048 points in the direct F2 dimension (^1^H) and 256 points in the F1 dimension. (^15^N).

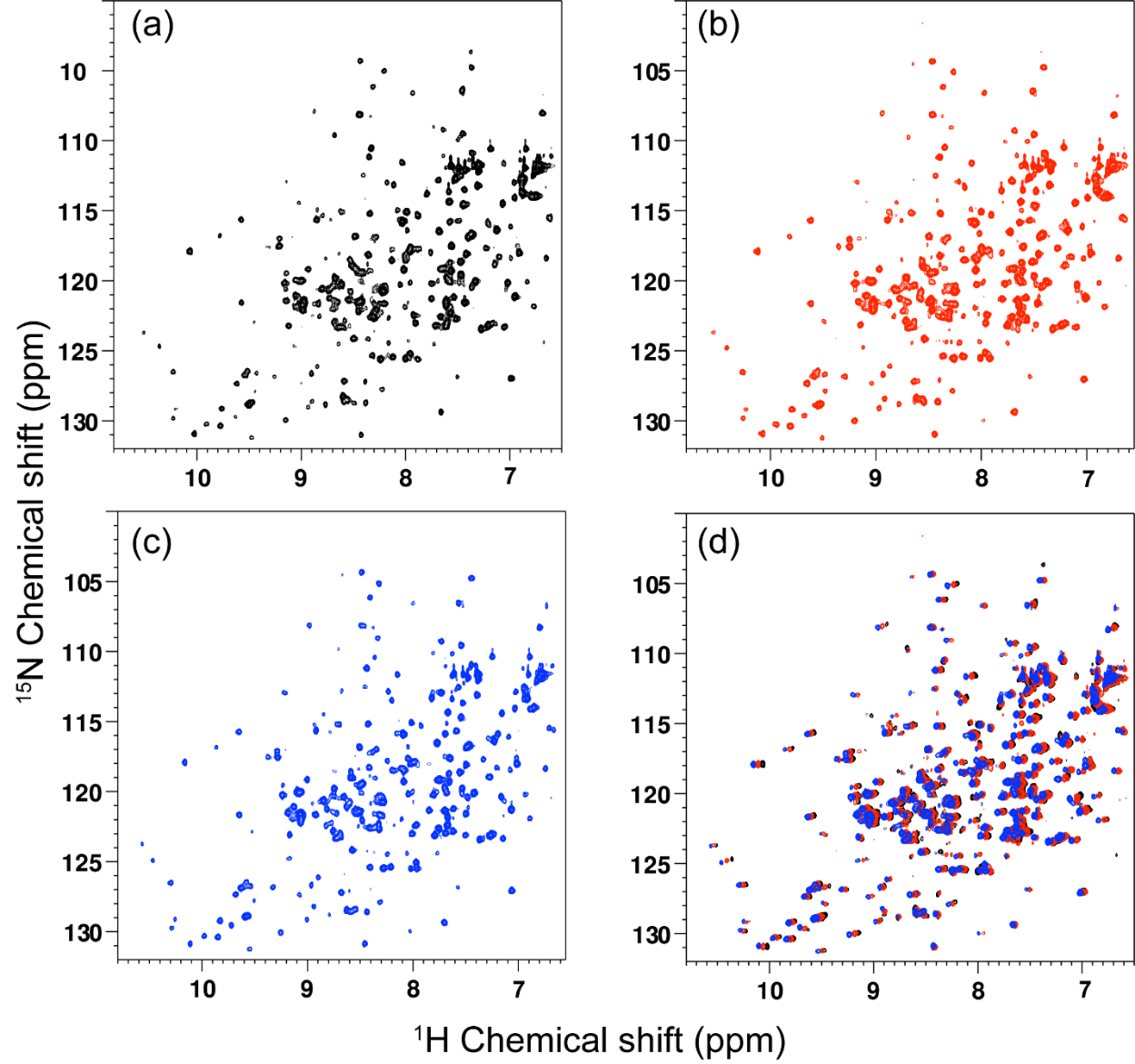

**Figure S5.** Temperature dependence of APE1. 2D ^1^H-^15^N HSQC spectra of [*U*-^15^N]-APE1 were collected using a 600 MHz Bruker AVANCE spectrometer equipped with a 5-mm triple-resonance cryoprobe. (a) 25 °C, (b) 30 °C, and (c) 35 °C. (d) The superimposition of APE1 ^1^H-^15^N HSQC spectra at 25 °C (black), 30 °C (red), and 35 °C (blue) indicate the protein is stable at high temperatures. All 2D ^1^H-^15^N NMR HSQC spectra were acquired with 2048 points in the direct F2 dimension (^1^H) and 256 points in the F1 dimension (^15^N).

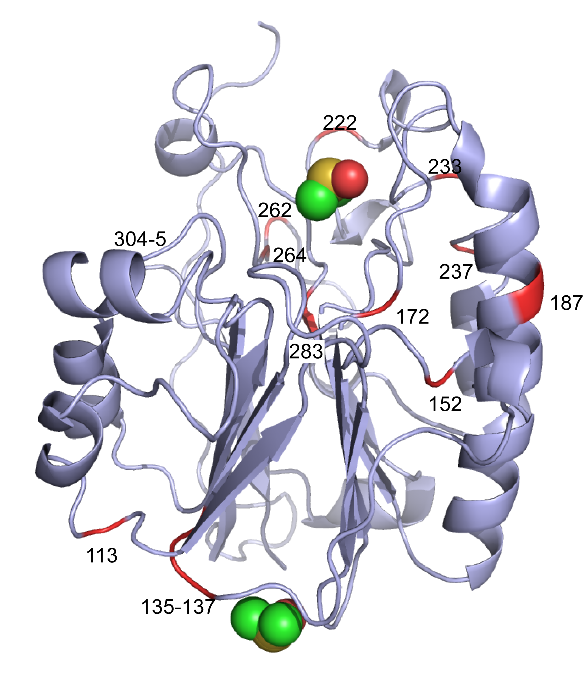

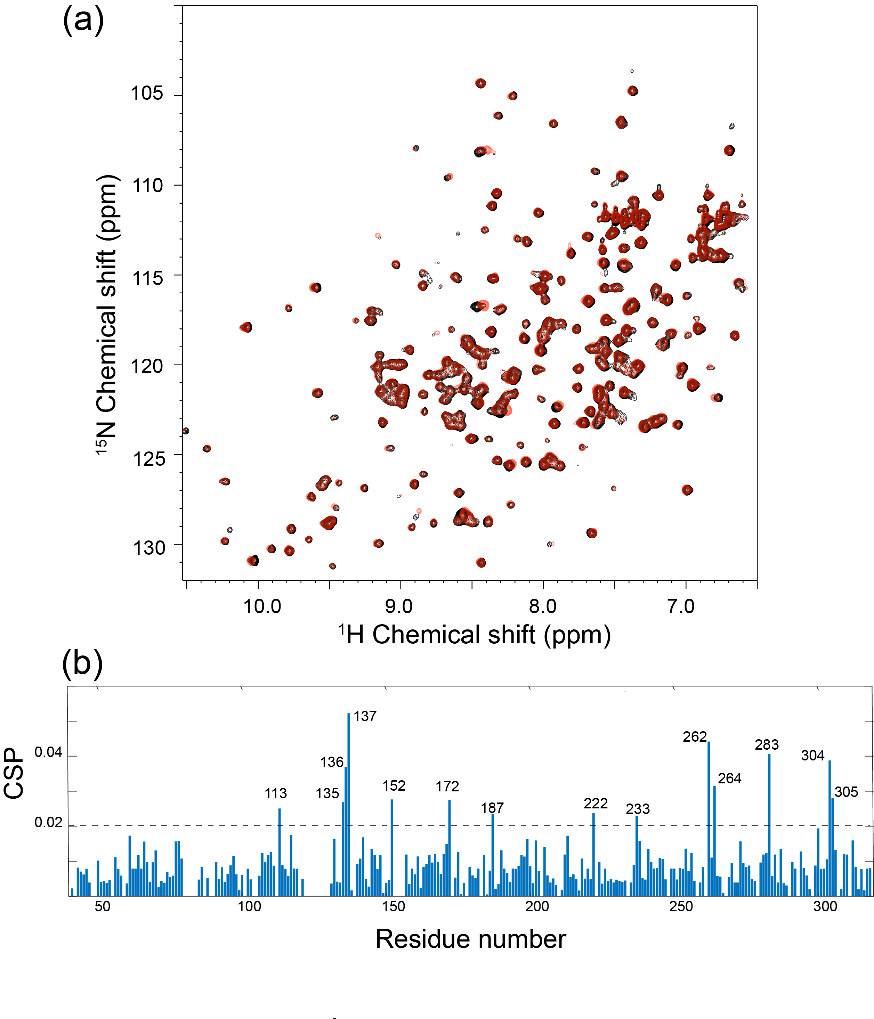

**Figure S6. DMSO-D6 binds APE1.** (a) To determine whether DMSO-D6 perturbs the APE1 2D ^1^H-^15^N HSQC spectrum, data were recorded with 0% (black) and 2% DMSO-D6 (red). (b) The chemical shift perturbation plot versus residue number indicates the binding of DMSO-D6 to APE1. All ^1^H-^15^N HSQC spectra of [*U*-^15^N]-APE1(90 μM) were collected using a 600 MHz Bruker AVANCE spectrometer equipped with a 5-mm triple-resonance cryoprobe at 25 °C. The 2D ^1^H-^15^N NMR HSQC spectra were acquired with 2048 points in the direct F2 dimension (^1^H) and 256 points in the F1 dimension. (^15^N). Residues with CSPs over 0.02 (Gly113, Ser135, Arg136, Gln137, Asp152, Val172, Arg187, Asn222, Thr233, Arg237, Tyr262, Tyr264, Asp283, Ala304, and Leu305) are mapped in red on a cartoon rendering of the structure of APE1 (light blue cartoon) with bound DMSO (space filling model) (PDB ID: 6MK3). The upper DMSO molecule is bound to the endonuclease active site. These results suggest significant binding to both the endonuclease active site and the small pocket. Other CSPs may reflect desolvation effects.

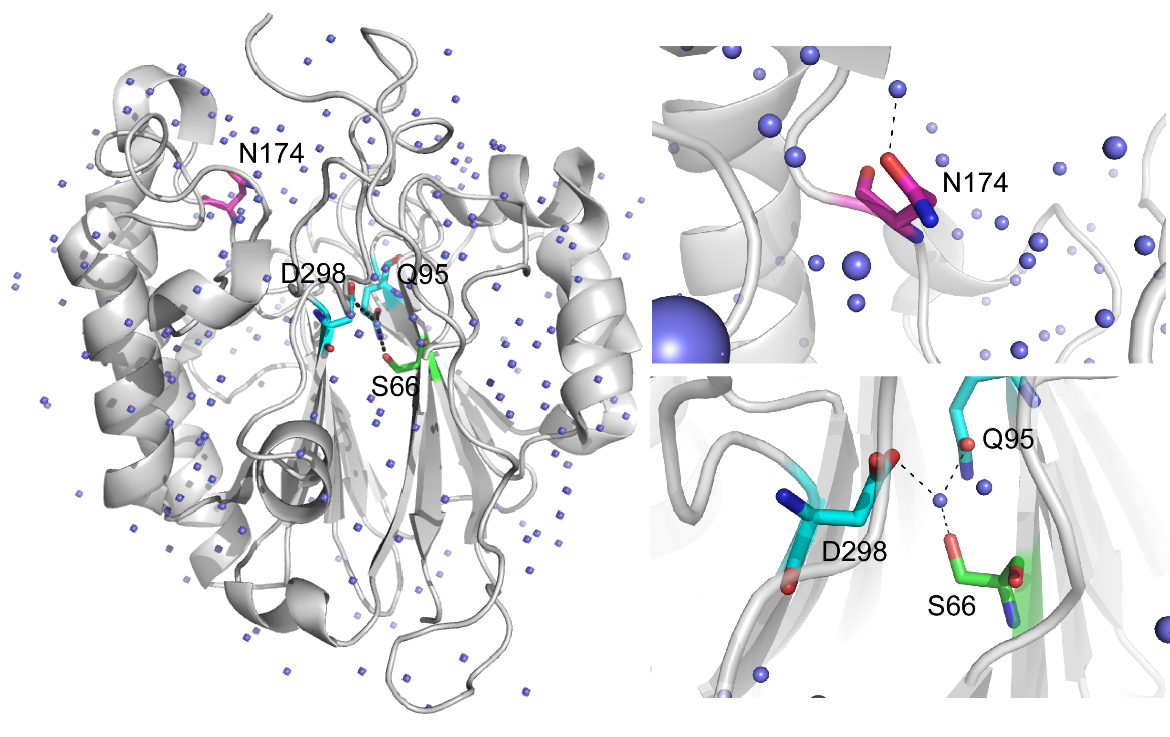

**Figure S7.** Residues Ser66, Gln95, and Asp298 hydrogen-bond to a water molecule in the structure of APE1 (PDB ID: 4QHD). Water molecules are shown as blue spheres, APE1 as a gray cartoon rendering, and residues as sticks with C, green for S66, magenta for N174, and cyan for D298 and Q95; O, red; and N, blue. N174 hydrogen bonds to a water molecule in the endonuclease active, while S66 hydrogen bonds to a water molecule that is also hydrogen-bonded to D298 and Q95. Addition of ethanol-D6 to [*U*-^15^N]-APE1 results in significant CSPs for S66 and N174 (see Fig. S8) in the 2D ^1^H-^15^N NMR HSQC spectrum. Addition of DMSO results in significant CSPs for numerous residues including D298.

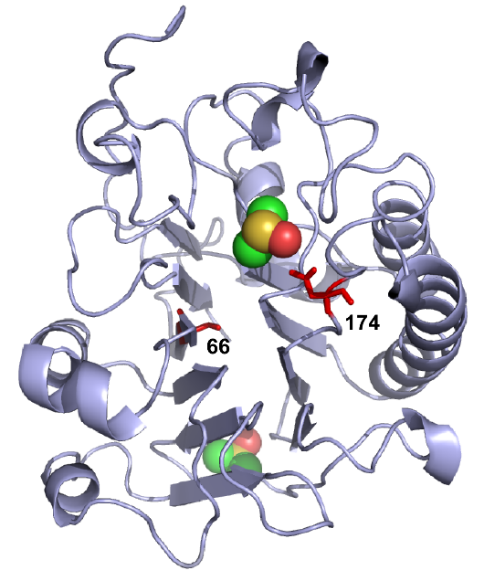

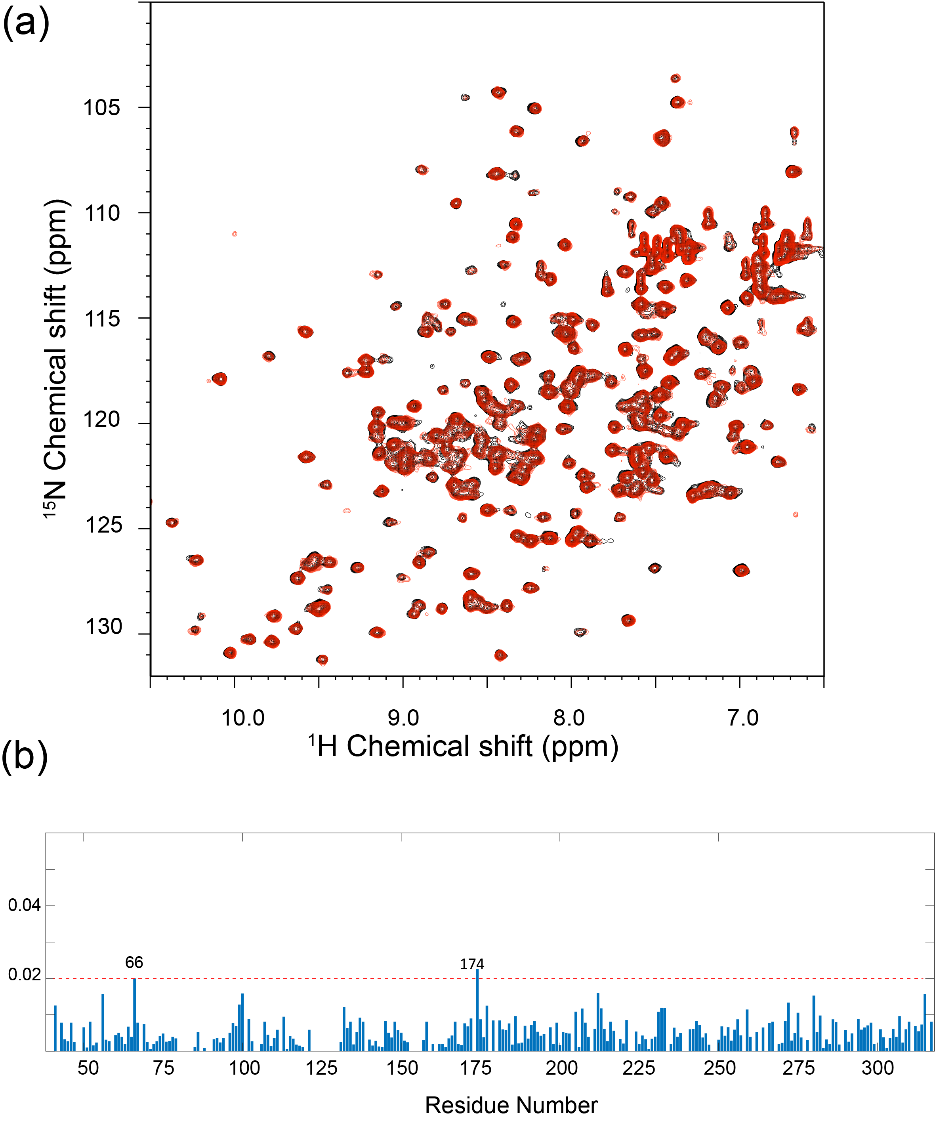

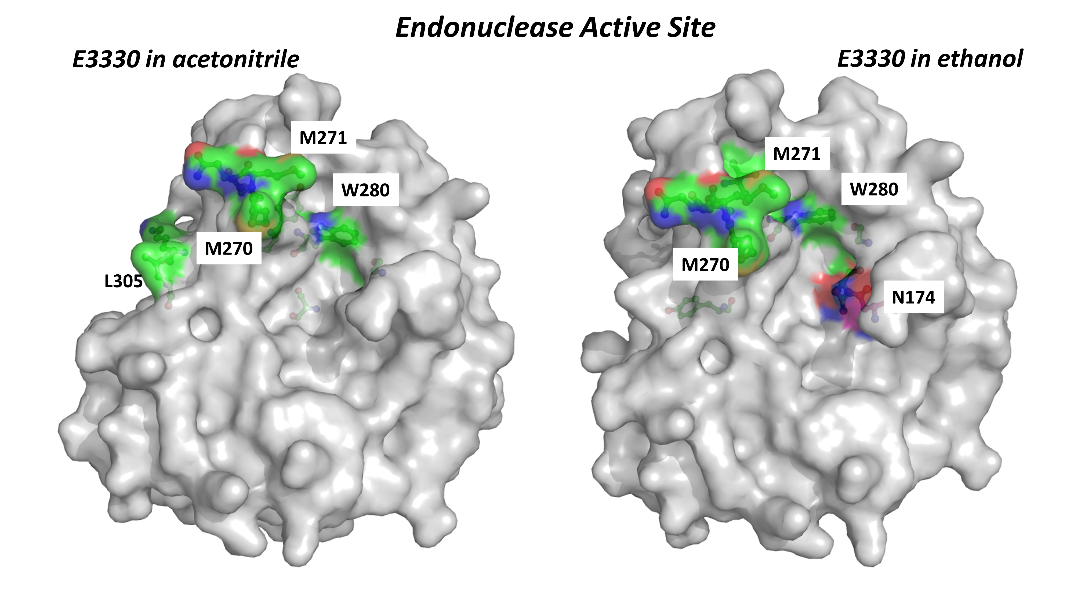

**Figure S8. Ethanol D-6 binds specifically to APE1.** (a) To determine whether ethanol-D6 perturbs the APE1 2D ^1^H-^15^N HSQC spectrum, data were recorded with 0% (black) and 2% methanol-D4 (red). (b) The chemical shift perturbation plot versus residue number indicates the specific binding of methanol-D4 to APE1. All ^1^H-^15^N HSQC spectra of [*U*-^15^N]-APE1(90 μM) were collected using a 600 MHz Bruker AVANCE spectrometer equipped with a 5-mm triple-resonance cryoprobe at 25 °C. The 2D ^1^H-^15^N HSQC spectra were acquired with 2048 points in the direct F2 dimension (^1^H) and 256 points in the F1 dimension (^15^N). Residues with CSPs over 0.02 (Ser66 and Asn174) are mapped in red on a cartoon rendering of the structure of APE1 (light blue cartoon) with bound DMSO (space filling model) (PDB ID: 6MK3). The upper DMSO molecule is bound to the endonuclease active site. A molecular surface rendering with APE1 in gray, active site residues M270, M271, and W280 (C, green) and N174 (C magenta) in ball-and-stick models. N174 borders the active site pocket that binds the abasic site within substrate DNA.

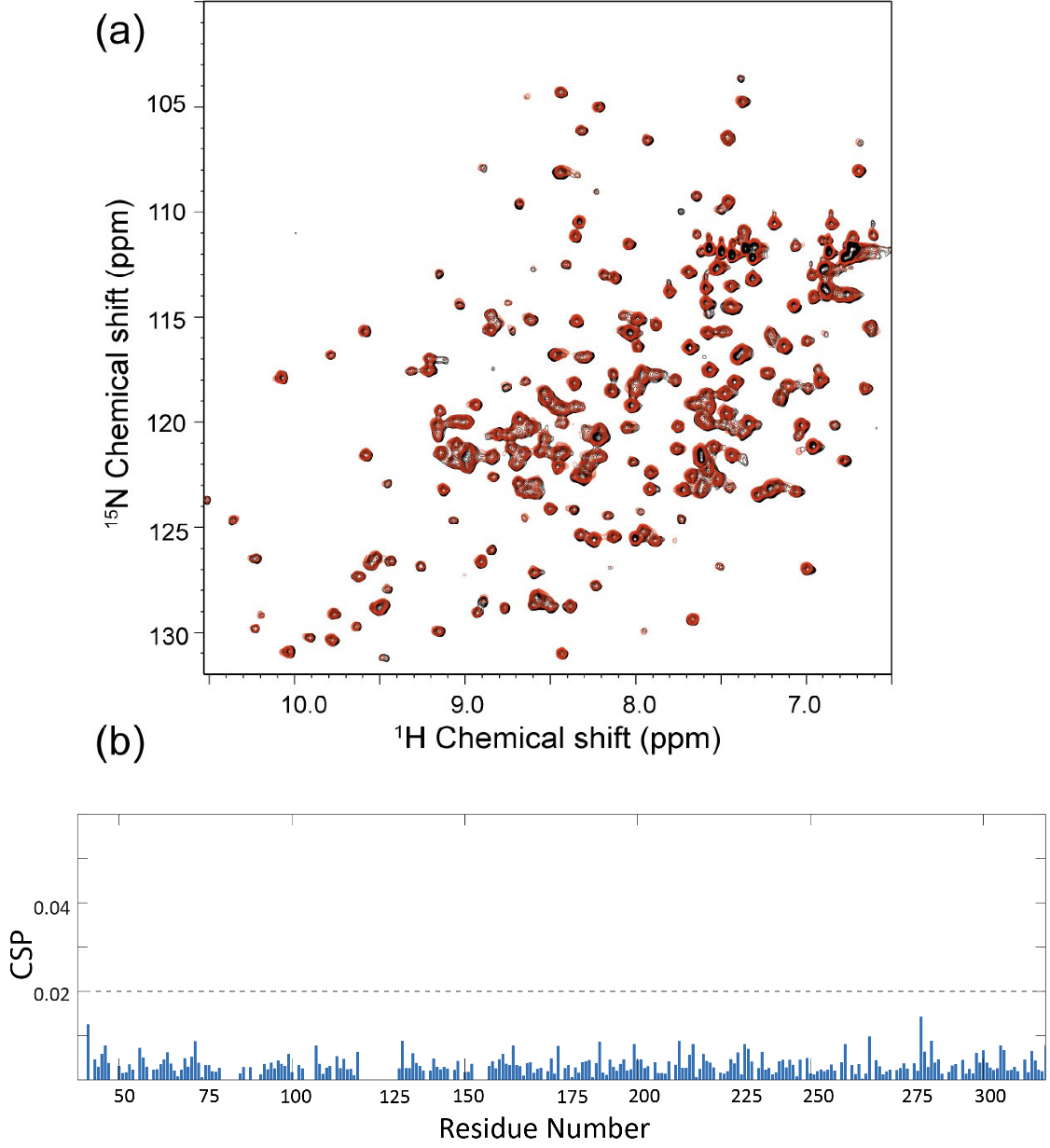

**Figure S9.** **Acetonitrile-D3 does not specifically bind APE1.** (a) In contrast to DMSO-D6, addition of 2% acetonitrile-D3 solvent (red) minimally perturbs the APE1 2D ^1^H-^15^N HSQC spectrum (black). (b) The chemical shift perturbation plot versus residue number indicates minimal impact from the addition of acetonitrile-D3 with no residues that have CSPs over 0.02. All ^1^H-^15^N HSQC spectra of [*U*-^15^N]-APE1(90 μM) were collected on a 600 MHz Bruker AVANCE spectrometer equipped with a 5-mm triple-resonance cryoprobe at 25 °C. The 2D ^1^H-^15^N NMR HSQC spectra were acquired with 2048 points in the direct F2 dimension (^1^H) and 256 points in the F1 dimension (^15^N).

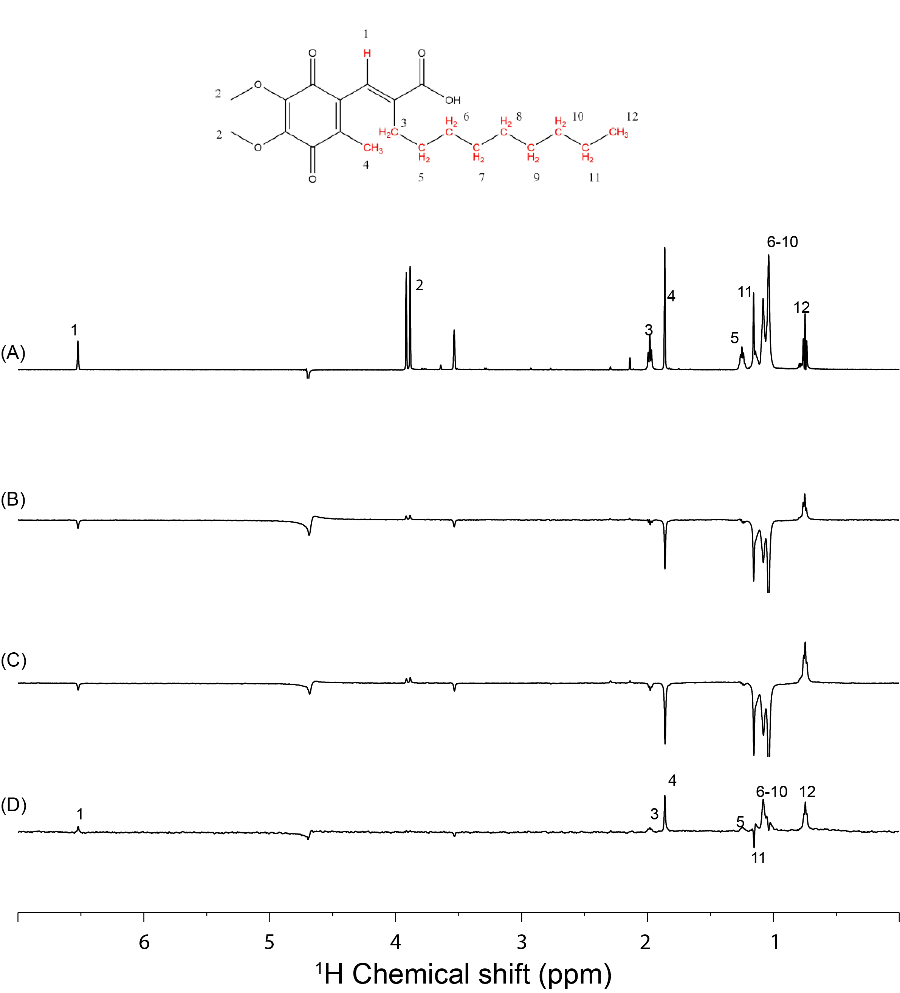

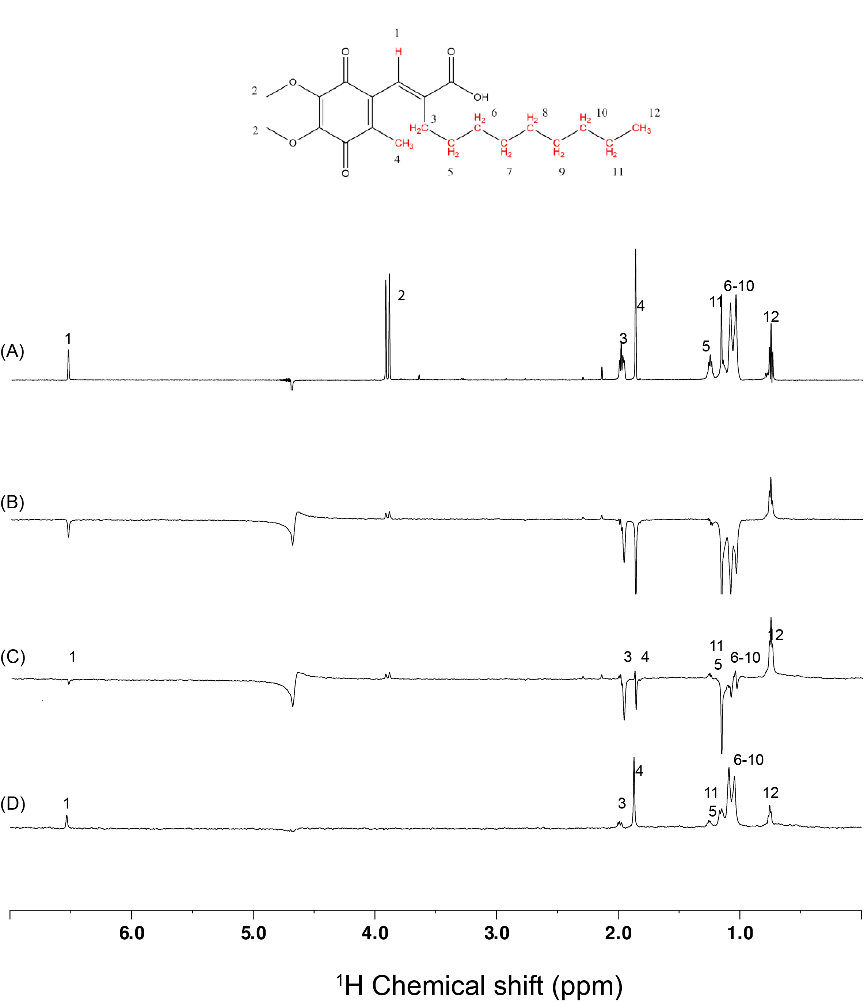

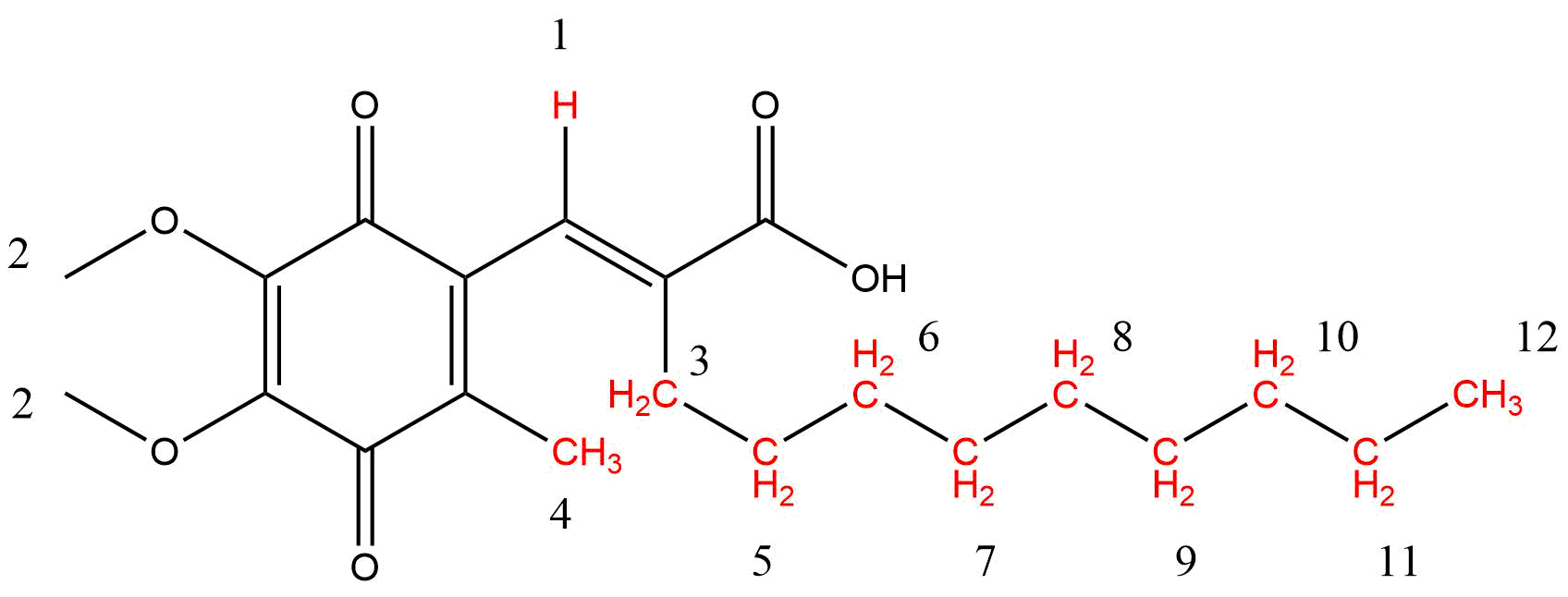

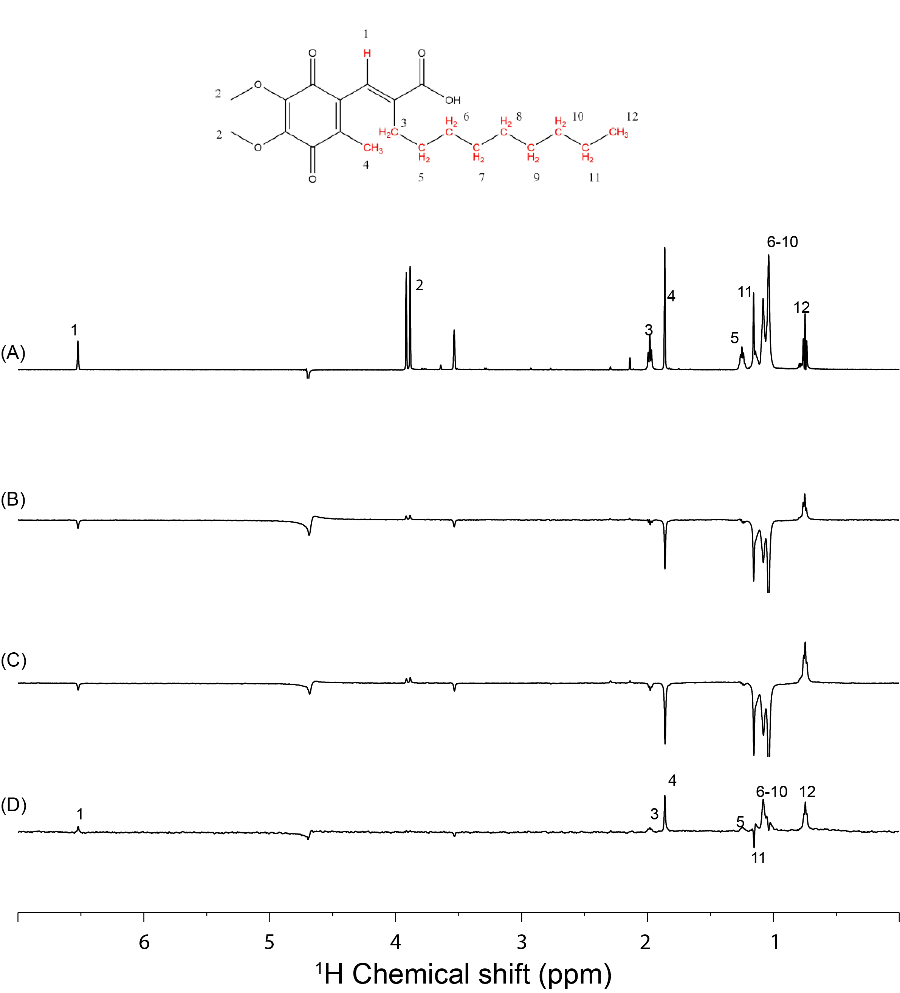
**Figure S10.** Interactions of APX3330 dissolved in acetonitrile-D3 (top panel) or ethanol-D6 (bottom panel) with APE1 were probed using a ligand-based WaterLOGSY NMR experiment. (A)The 1D ^1^H-spectrum was measured for 200 μM APX3330 (stock solution 50 mM dissolved in acetonitrile-D3 or ethanol-D3) in 20 mM sodium phosphate, 0.1 M NaCl pH 6.5, 0.5 mM DTT, 0.2 mM EDTA. WaterLOGSY experiments were done in the absence and presence of 20 μM APE1. (B) WaterLOGSY spectrum in the absence of APE1. (C) WaterLOGSY spectrum in the presence of APE1. (D) WaterLOGSY difference spectrum (spectrum C – spectrum B) indicates changes in the water accessibility for all assigned protons except the methoxy protons, labeled 2 in the spectra (See numbering scheme on structure of APX3330). The waterLOGSY spectrum was collected on a 600 MHz Bruker AVANCE spectrometer using a TCI cryoprobe with a mixing time of 1.5 s. The data were collected with 128 scans, a sweep width of 10 ppm, an acquisition time of 1.58 s, and a relaxation delay of 5 seconds, and processed using Topspin 3.6.2.

APX3330/ethanol-D6

APX3330/acetonitrile-D3

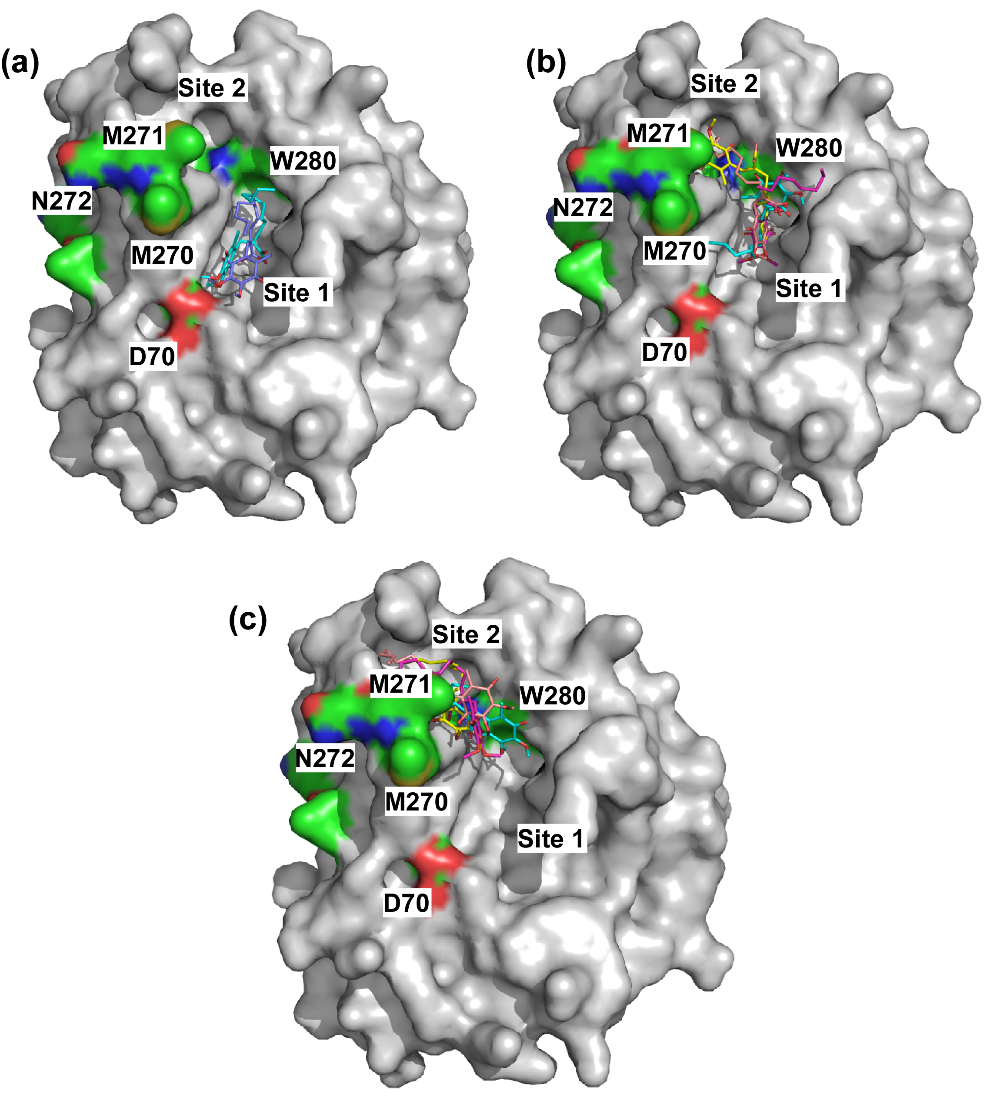

**Figure S11.** Docking of APX3330 to the endonuclease active site reveals several different poses. A gray surface rendering of APE1 (4QHD) is shown with residues within or near the endonuclease active site highlighted with C (green), oxygen, (red), nitrogen (blue), and sulfur (yellow). Selected low energy poses of APX3330 bound to APE1 obtained from AutoDock Vina and AutoDock 4.2 are shown. (a) In site 1, the nonyl alkyl tail of APX3330 poses, shown in blue (AutoDock Vina) and cyan (Autodock 4.2), is positioned close to Trp280 and the dimethoxyquinone points out into the rest of this large pocket. The blue pose is very similar to that previously reported. (b) Alternate poses of APX3330 from AutoDock 4.2 (shown in cyan, and magenta) place the nonyl alkyl tail outside site 1. AutoDock Vina poses (yellow and orange) bridge an adjacent site 2 and site 1 with the nonyl alkyl tail pointing either into either site. (c) Other alternate poses from AutoDock 4.2 (yellow and cyan) and Autodock Vina (orange and magenta) position the nonyl alkyl tail within site 2 and the dimethoxyquinone ring adjacent to W280 and M270. Energetic differences between docked poses in site 1 versus site 2 are very close (within 0.5-0.6 kcal/mol) for results from both programs. This range is too close to definitively say APE3330 only binds in one site or the other. Energetically, if you only took the result with the lowest binding energy, Autodock Vina would slightly prefer site 1 by 0.6 kcal/mol while AutoDock 4.2 would prefer site 2 by 0.3 kcal/mol.

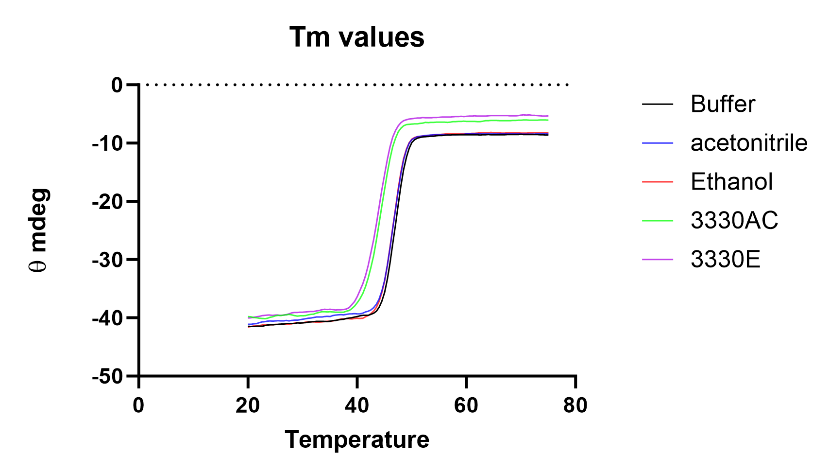

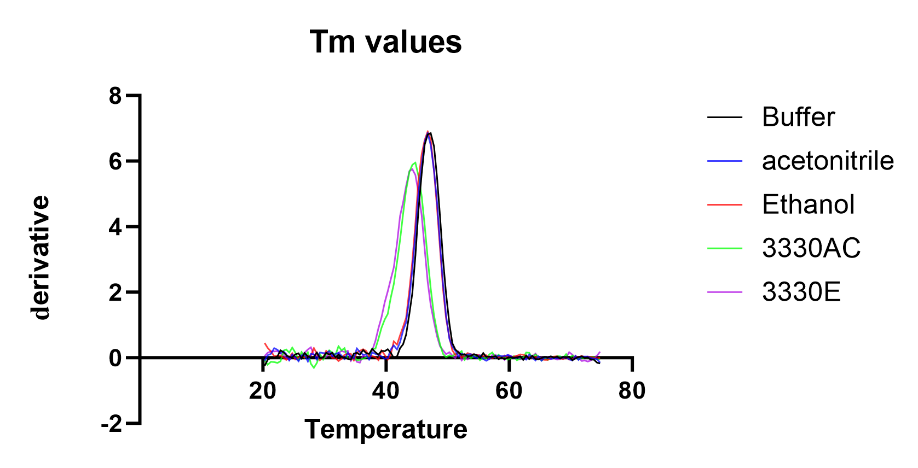
**Figure S12.** Thermal melting curves for APE1 were determined using 20 μM APE1 buffered in 20 mM sodium phosphate pH 6.5, 100 mM NaCl in the presence or absence of 0.4% ethanol, 0.4% acetonitrile, 200 μM APX3330 dissolved in ethanol, or 200 μM APX3330 dissolved in acetonitrile. The temperature was gradually increased (1 °C/min) from 20 to 75 °C using the Peltier temperature controller on the Jasco J-1500 instrument with mdeg monitored at 210 nm. Addition of acetonitrile lowered the melting temperature of APE1 from 47.3 °C by 0.5° and 1 °C, respectively. Addition of APX3330 dissolved in either acetonitrile or ethanol lowered the melting temperature by 2 ° and 2.5 °C, respectively.

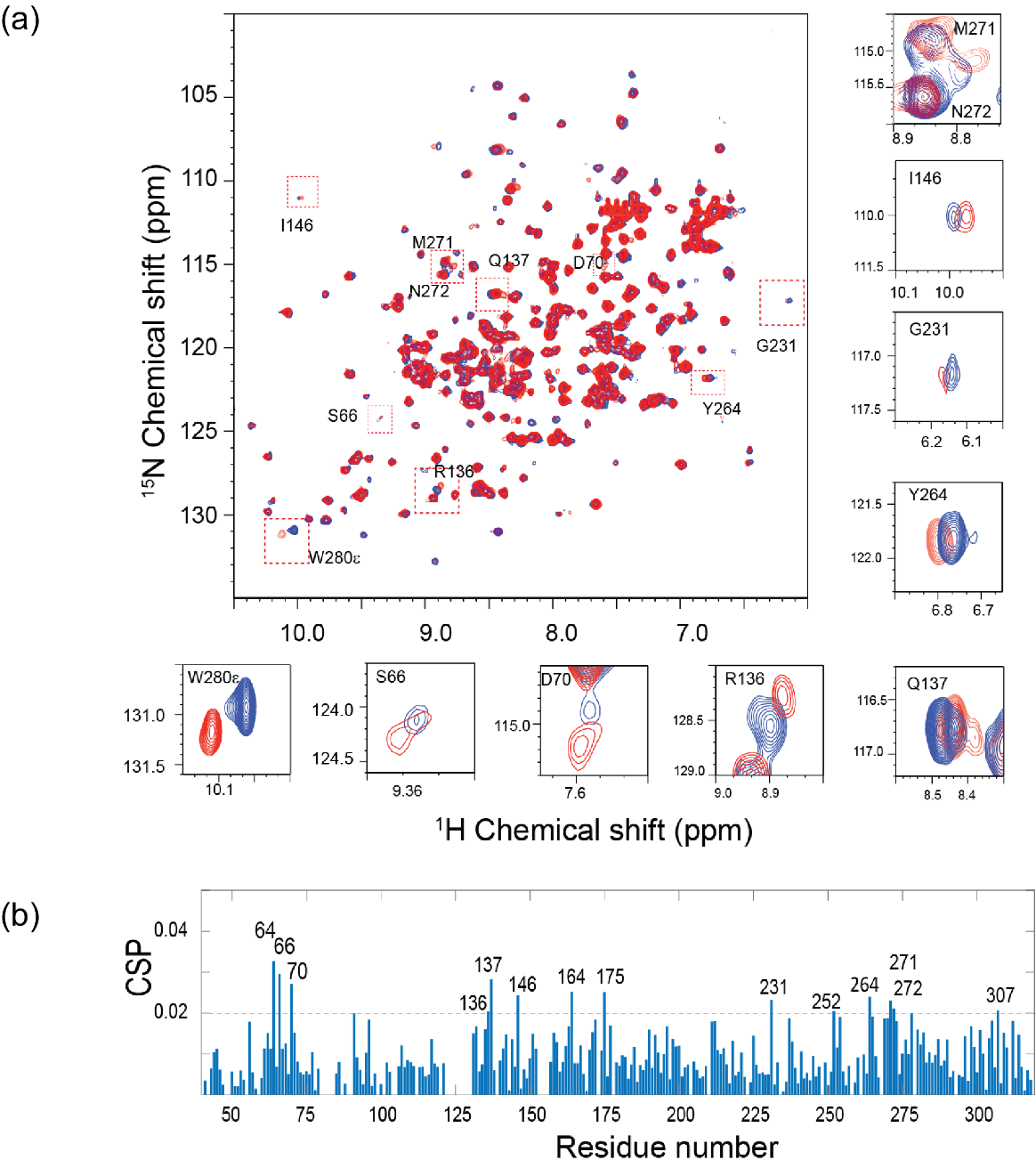

**Figure S13.** Interaction of a 10-fold molar excess of APX3330 (co-solvent acetonitrile) to [*U*-^15^N]-APE1 from titration experiment. (a) 2D ^1^H-^15^N HSQC spectral overlay of the APE1 (85 μM) spectrum (blue) and with 850 µM APX3330 (red). Each cross peak corresponds to one backbone or side chain amide. Specific chemical shift perturbations are shown in small boxes. (b) The chemical shift perturbations versus residue number reveal widespread interactions with APE1. All 2D ^1^H-^15^N NMR HSQC spectra were acquired with 2048 points in the direct F2 dimension (^1^H) and 256 points in the F1 dimension (^15^N).

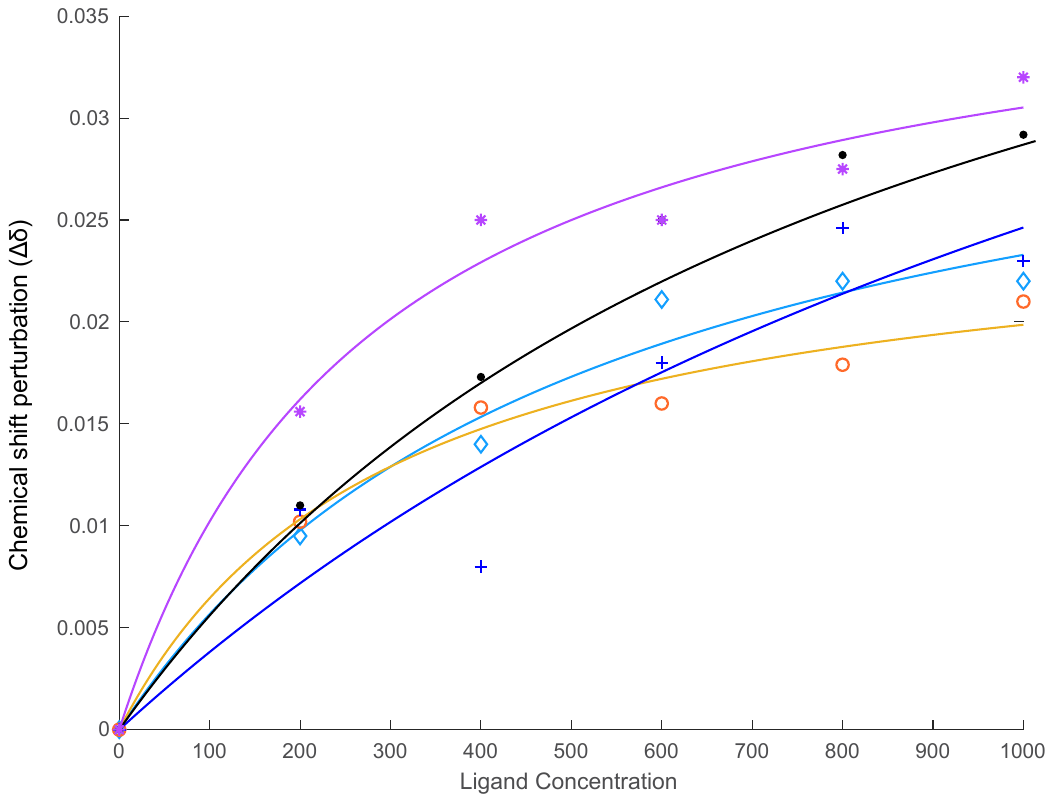

**Figure S14.** The binding affinity of APE1^ΔN40^ for APX3330. The APX3330 binding affinity was determined from the dependence of shift perturbation (Δδ) on APX3330 concentration using MATLAB for multiple residues giving a K_d_ = 500± 250 μM. Data for Ile64 (*), Gln137 (●), Ala175(+), Met271(◇), and Asn272 (o) were used to determine a K_d_.

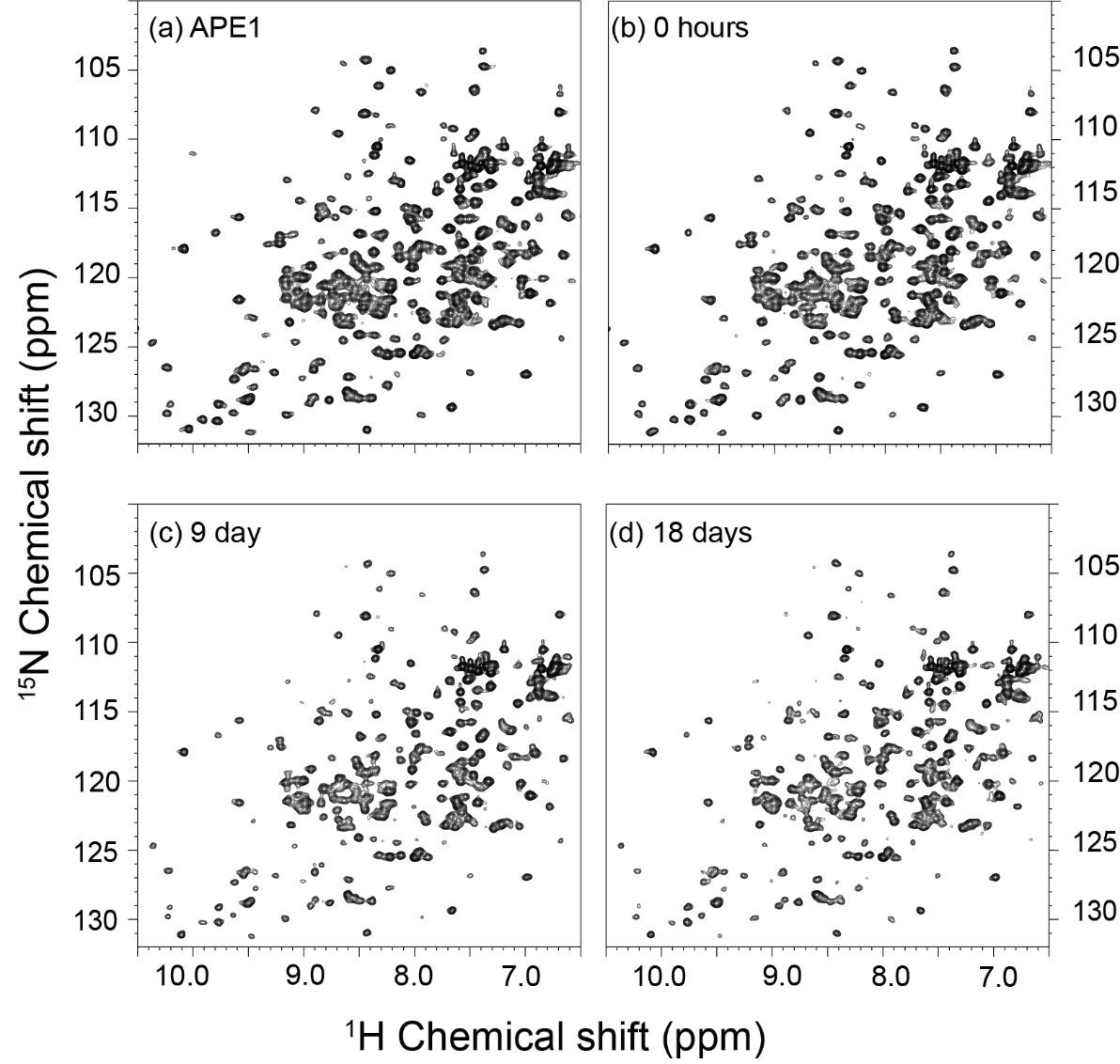
**Figure S15.** Time dependent effects of APX3330/ethanol-D6 on [*U*-^15^N]-APE1. A time course of ^1^H-^15^N HSQC spectra were collected at 25 °C following addition of a 10-fold molar excess of APX3330. (a) APE1 spectrum prior to addition of APX3330/ethanol-D6 (b) 0 hours, (c) 9 days after addition, and (d) 18 days after addition of a 10-fold molar excess of APX3330/ethanol-D6. No loss of chemical shifts is evident up to 18 days following the addition of APX3330 using ethanol as a co-solvent in contrast to the distinct chemical shift losses observed for APX3330 using acetonitrile as a co-solvent. All 2D ^1^H-^15^N NMR HSQC spectra were acquired with 2048 points in the direct F2 dimension (^1^H) and 256 points in the F1 dimension (^15^N).

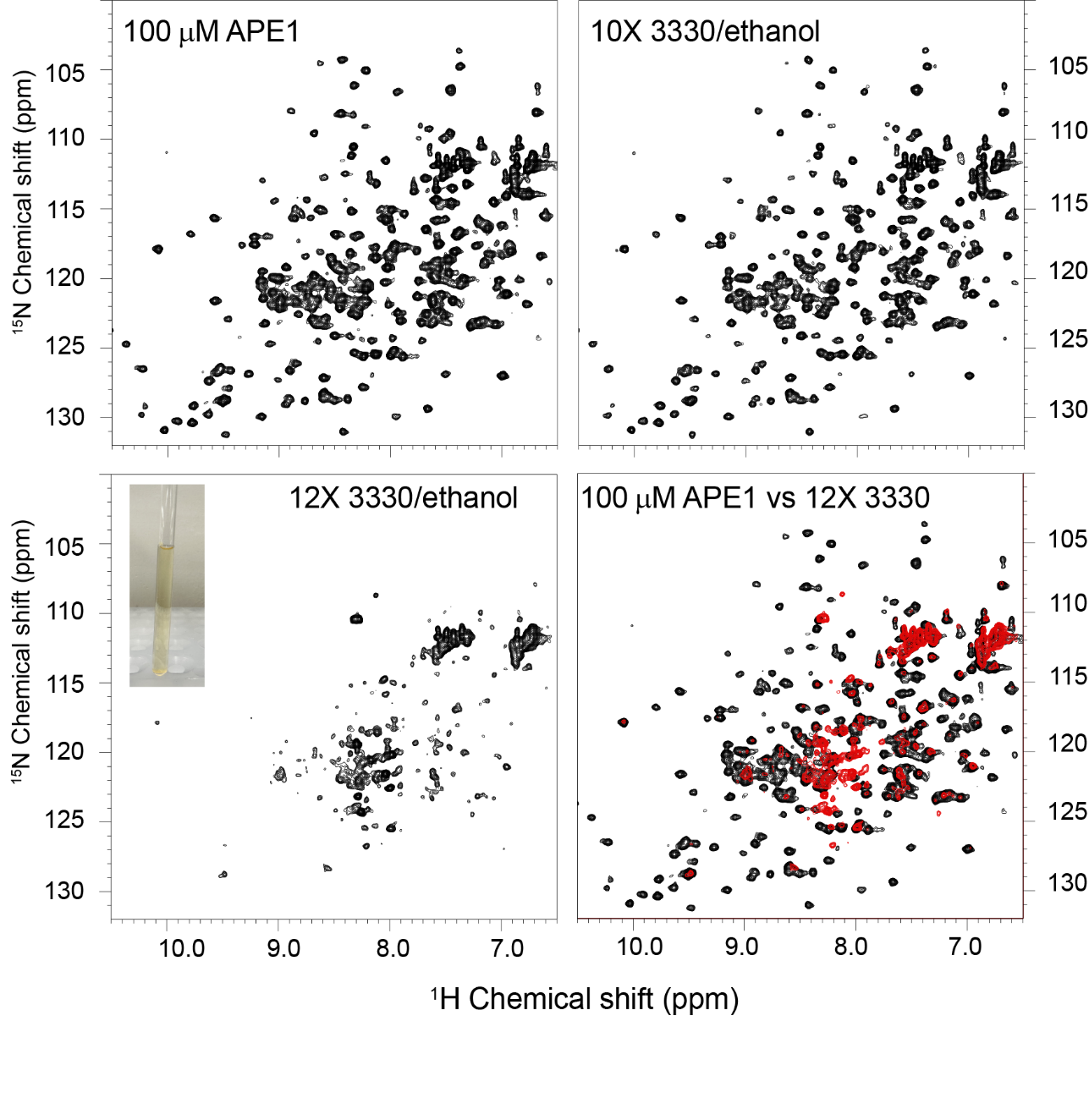

**Figure S16.** Effect of APX3330/ethanol-D6 on [*U*-^15^N]-APE1 (100 μM) (top left panel) at a 10-fold (top right panel) and 12-fold molar excess (bottom left panel) is shown for ^1^H-^15^N HSQC spectra were collected at 25 °C. An image of the precipitated [*U*-^15^N]-APE1 in the bottom of the NMR tube following addition of a 12-fold molar excess of APX3330/ethanol-D6 is shown in the panel inset for bottom left panel. An overlay of the ^1^H-^15^N HSQC spectrum for [*U*-^15^N]-APE1 (100 μM) in black and 12-fold molar excess of APX3330/ethanol in red (bottom right panel) highlights a collapse of the cross peaks toward the central diagonal consistent with the behavior of an unfolded protein. This spectrum is clearly distinct from that of the protein following addition of a 12-fold molar excess of APX3330/acetonitrile-D3. All 2D ^1^H-^15^N NMR HSQC spectra were acquired with 2048 points in the direct F2 dimension (^1^H) and 256 points in the F1 dimension (^15^N).

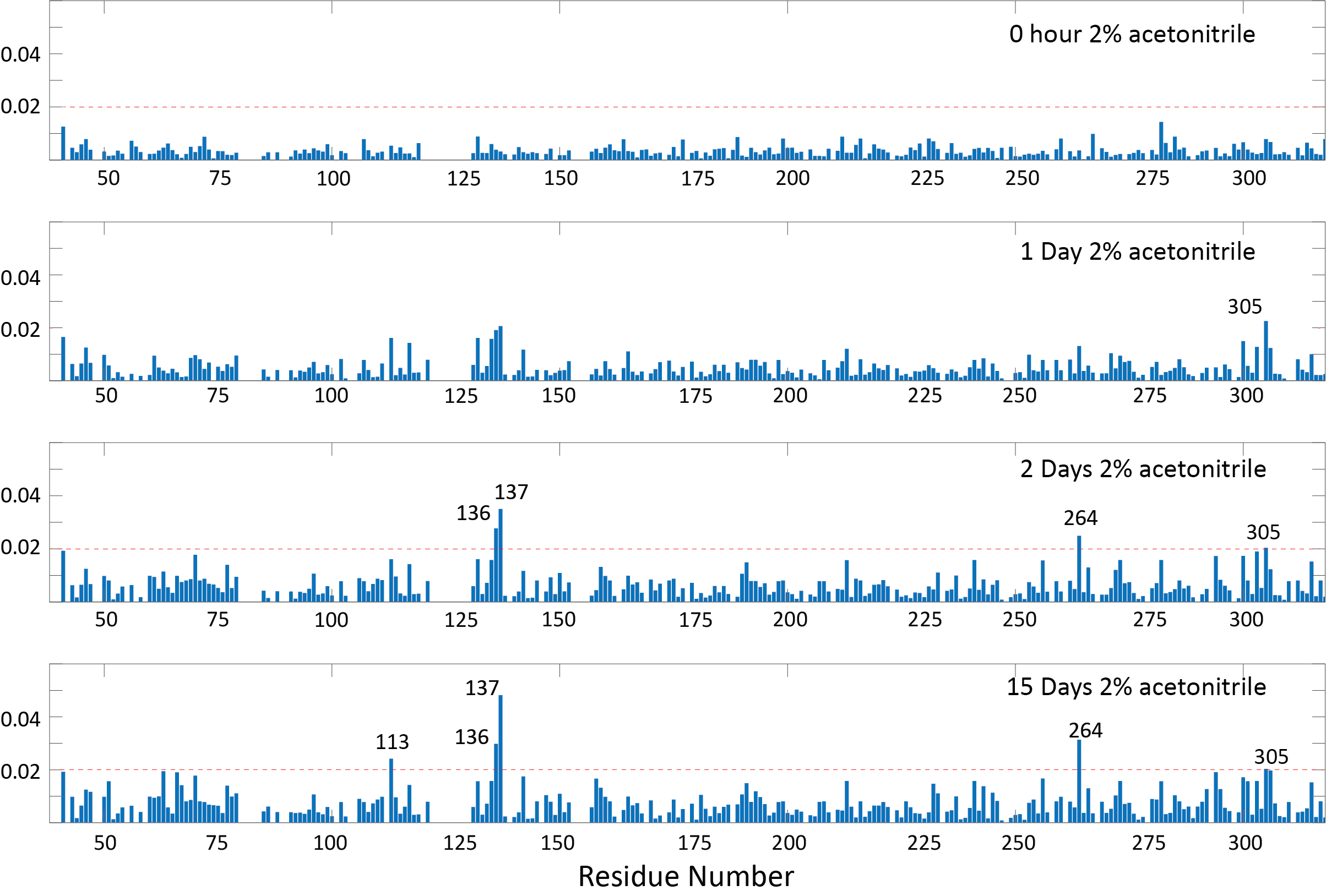
**Figure S17**. Effect of acetonitrile-D3 on [*U*-^15^N]-APE1 over time. CSPs are shown for time point 0 hours, 1 day, 2 days, and 15 days after addition of acetonitrile-D3 to APE1. No loss of chemical shifts was observed over this time period. After 48 hours, CSPs over 0.02 for residues within the small pocket Arg136 and Gln137 appeared. Other residues with significant CSPs include 264 and 305; these residues do not define a specific pocket on the surface of APE1. All 2D ^1^H-^15^N NMR HSQC spectra were acquired with 2048 points in the direct F2 dimension (^1^H) and 256 points in the F1 dimension (^15^N).

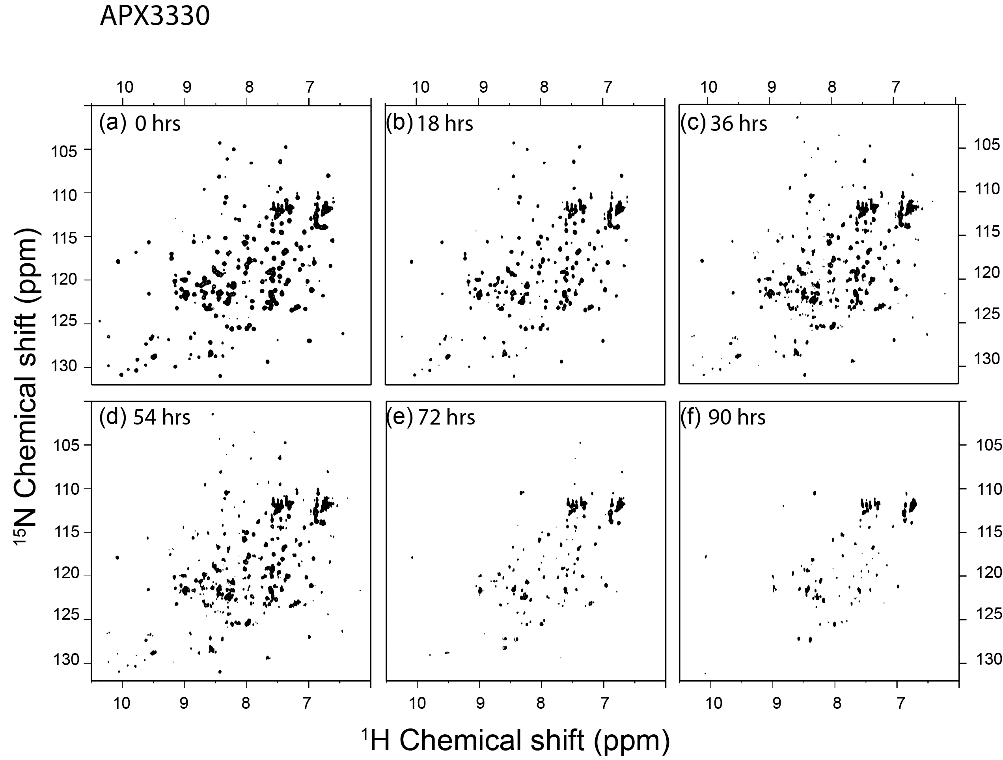

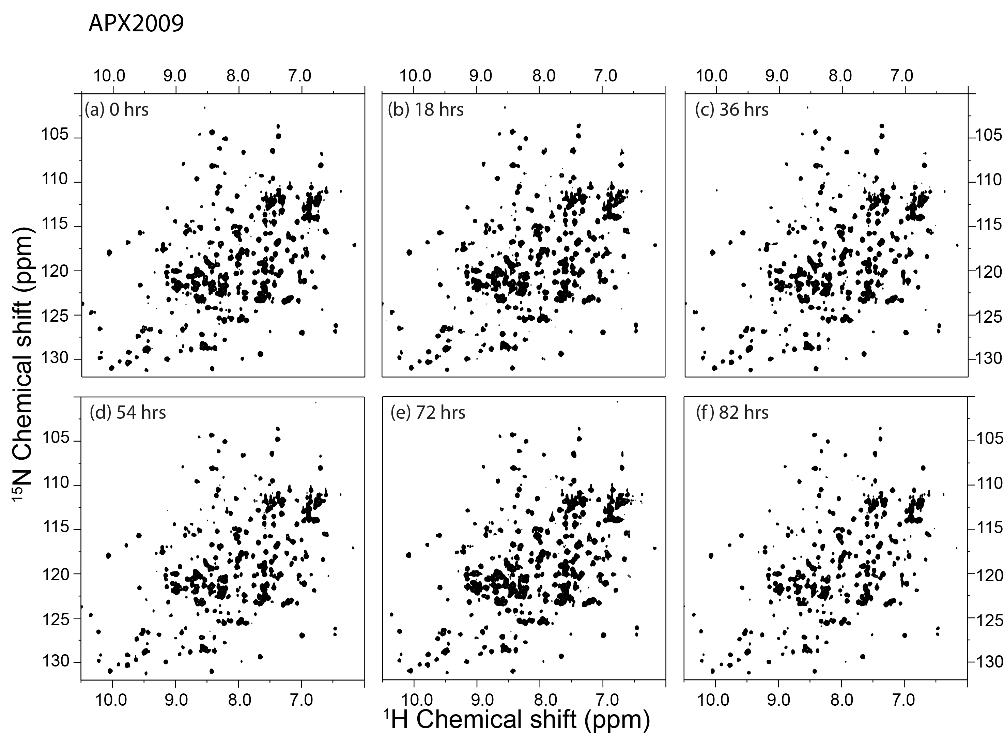
**Figure S18.** Time-dependent effects of APX2009 (top) and APX3330 (bottom) on [*U*-^15^N]-APE1 at 25 °C. ^1^H-^15^N HSQC spectra were collected at 25 °C for APE1 following the addition of compound at (a) 0 hours (b) 18 hours (c) 36 hours (d) 54 hours (e) 72 hours and (f) 82 or 90 hours. There is no loss of cross peak intensities up to 82 hrs for incubation with APX2009, while for APX3330, the cross peak intensities decrease as time progresses. After about 72 hours, approximately 35% of the backbone resonances are lost. All ^1^H-^15^N HSQC spectra of [*U*-^15^N]-APE1(90 μM) were collected on a 600 MHz Bruker AVANCE spectrometer equipped with a 5-mm triple-resonance cryoprobe. The 2D ^1^H-^15^N NMR HSQC spectra were acquired with 2048 points in the direct F2 dimension (^1^H) and 256 points in the F1 dimension. (^15^N).

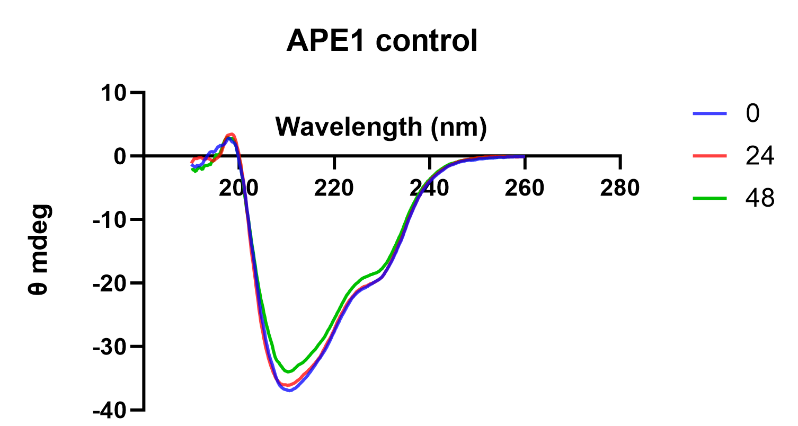

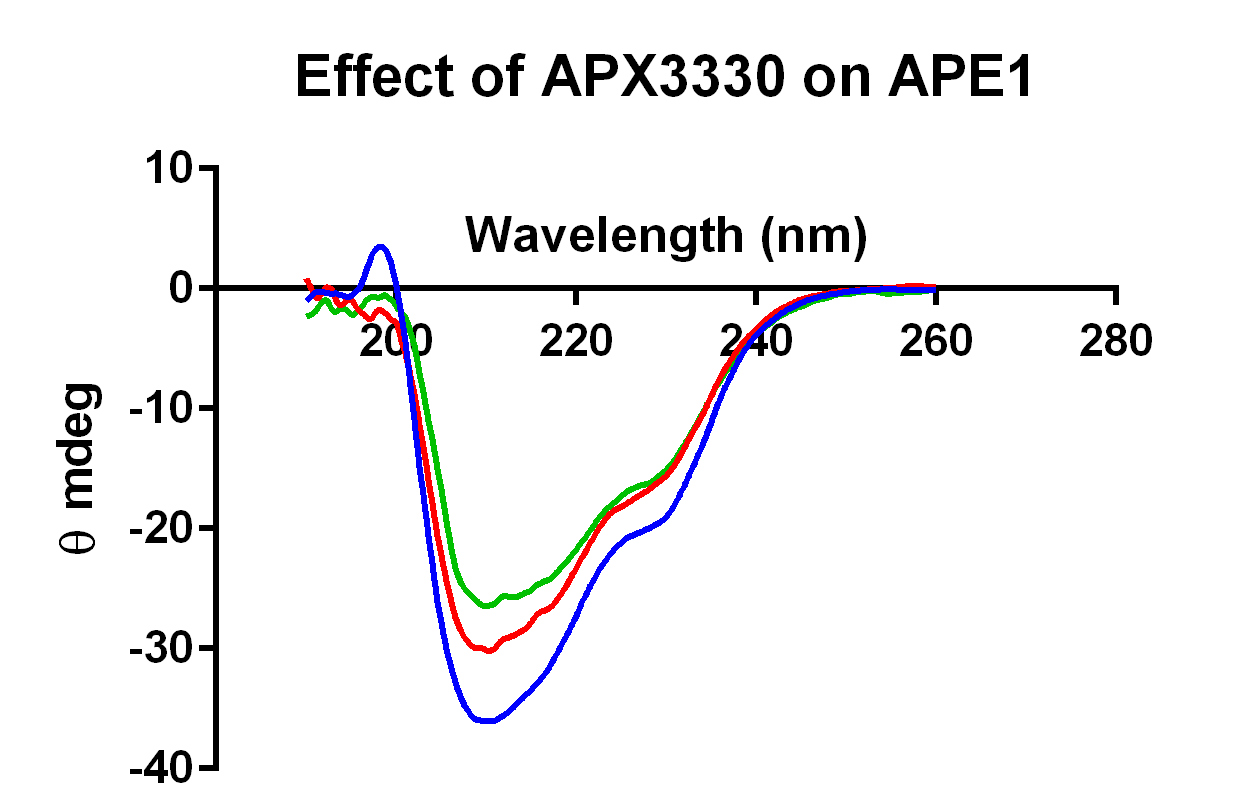

**Figure S19.** Effect of APX3330 on the circular dichroism (CD) spectrum of APE1 measured at 30 °C. Samples were prepared identically to those used for NMR studies in 20 mM sodium phosphate, 0.1 M sodium chloride pH 6.5. APX3330 was dissolved in acetonitrile and added at an 8-fold molar excess. CD spectra for APE1 (blue), APE1 with APX3330 after 24 hours (red) or 48 hours (green) were measured on a Jasco J-1500 circular dichroism instrument with Pelletier temperature control (left panel). Control samples for 0, 24, and 48 hr time points with added acetonitrile and no APX3330 are shown on the right for comparison. Within 48 hours there is a small effect on the CD signal at 30 °C. Two negative bands were observed, one at approximately 210 nm consistent with alpha helical structure and a second band at approximately 230 nm that includes contributions from both beta and alpha helical secondary structural elements.

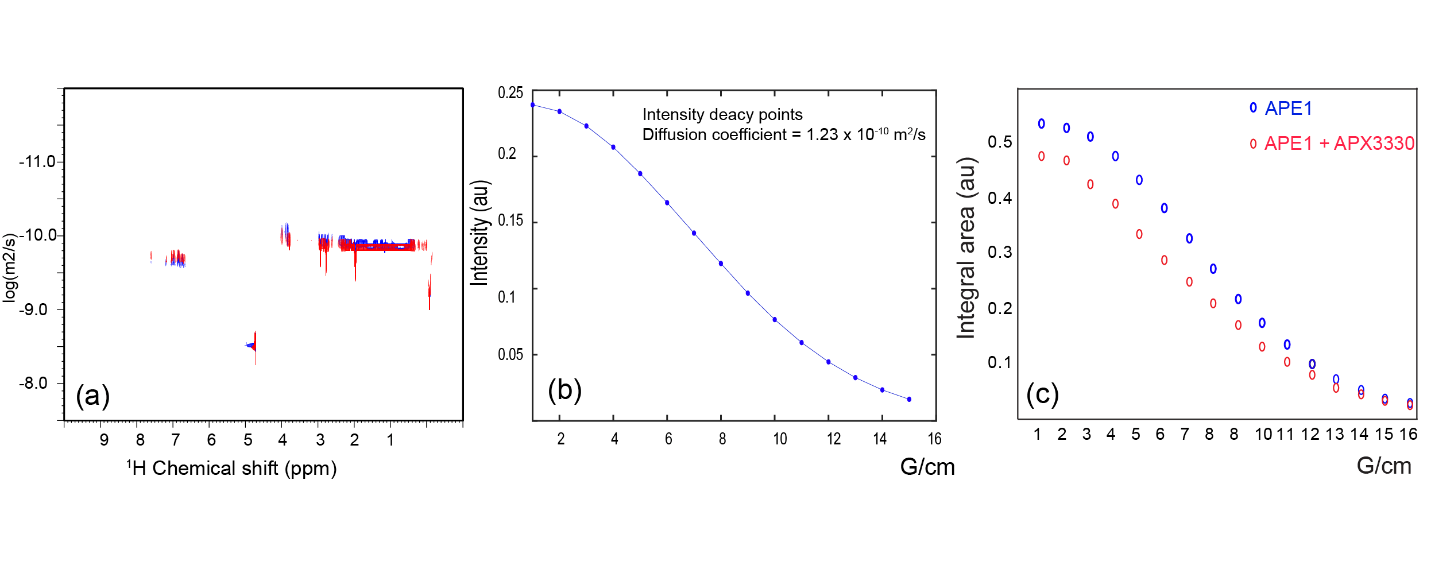

**Figure S20.** Characterization of APE1 oligomerization/aggregation in the presence or absence of APX3330 by DOSY NMR. (a) The DOSY NMR spectra are plotted for APE1 (blue) and APE1 plus an 8-fold molar excess of APX3330 (red). The diffusion coefficient (D) and molecular weight were calculated by DOSY NMR using DSS (D=8.216x10^-10^ m^2^/s) as the reference. (b) The diffusion coefficient calculated for APE1 was (D=1.123 x 10^-10^ m^2^/s) which corresponds to 32.4 kDa and with an 8-fold molar excess of APX3330 (D=1.124 x 10^-10^ m^2^/s), which corresponds to 31 kDa. (c) The integral of the peak for APE1 (blue) and APE1 plus an 8-fold molar excess of APX3330 (red). DOSY spectra were collected on a 600 MHz Bruker AVANCE spectrometer at 30 °C using a TCI cryoprobe with a relaxation time of 5 s. The DOSY time interval (Δ) and gradient pulse duration (δ) were set at 600 ms and 1.2 ms, respectively. A total of 16 gradient increments, ranging linearly from 2% to 98%, were acquired with 1024 scans for each increment. Topspin 3.6.2 was employed for NMR data processing.

**Figure S21.** APE1 was incubated with an 8-fold molar excess of APX3330 as in our time-dependent NMR experiments for 1 week (in this case at room temperature, comparable to the experiment shown in SI Fig. S9). The sample was subjected to chromatographic analysis on a 24 ml Superdex 200 column equilibrated in 20 mM sodium phosphate, pH 6.5, 0.1 M NaCl (the same buffer that was used for the NMR experiments). As shown by the chromatographic measurements in blue, a large peak for APE1 elutes at 16 ml, exactly the same elution volume as the control APE1 sample shown in orange. There is a small peak of APE1 that elutes at approximately 14 ml, which would correspond to approximately a dimer of APE1. This peak is very small relative to the main peak. The broad peak eluting at 30 ml corresponds to APX3330, which separates from APE1 during the chromatographic separation as expected for a complex with an apparent K_D_ over 1 μM.

SEC Analysis

|  | APE1+3330 | APE1 |
| --- | --- | --- |
| 0 | 0 | -0.001 |
| 0.0372904 | 0 | 0 |
| 0.0745808 | 0.005 | 0 |
| 0.111871 | 0.006 | 0.003 |
| 0.149162 | 0.001 | -0.001 |
| 0.186452 | 0.003 | -0.003 |
| 0.223742 | 0.003 | -0.003 |
| 0.261033 | 0.004 | 0 |
| 0.298323 | 0 | -0.001 |
| 0.335614 | -0.001 | -0.006 |
| 0.372904 | 0.008 | -0.001 |
| 0.410194 | 0.012 | -0.001 |
| 0.447485 | 0.008 | 0 |
| 0.484775 | 0.002 | 0 |
| 0.522066 | 0.001 | 0 |
| 0.559356 | 0.003 | -0.006 |
| 0.596646 | 0.005 | -0.005 |
| 0.633937 | 0.004 | -0.004 |
| 0.671227 | 0.005 | -0.008 |
| 0.708518 | 0.005 | -0.003 |
| 0.745808 | 0.006 | 0.002 |
| 0.783098 | 0.004 | 0.002 |
| 0.820389 | 0.006 | 0 |
| 0.857679 | 0.007 | -0.002 |
| 0.89497 | 0.008 | -0.002 |
| 0.93226 | 0.01 | 0 |
| 0.96955 | 0.004 | 0.001 |
| 1.00684 | 0.004 | 0.002 |
| 1.04413 | 0.008 | 0.005 |
| 1.08142 | 0.007 | 0.012 |
| 1.11871 | 0.012 | 0.013 |
| 1.156 | 0.014 | 0.015 |
| 1.19329 | 0.007 | 0.018 |
| 1.23058 | 0.009 | 0.015 |
| 1.26787 | 0.007 | 0.025 |
| 1.30516 | -0.001 | 0.03 |
| 1.34245 | 0.001 | 0.031 |
| 1.37974 | 0.002 | 0.037 |
| 1.41704 | -0.003 | 0.042 |
| 1.45433 | 0 | 0.042 |
| 1.49162 | 0.002 | 0.042 |
| 1.52891 | -0.001 | 0.048 |
| 1.5662 | 0 | 0.052 |
| 1.60349 | 0.003 | 0.055 |
| 1.64078 | -0.002 | 0.06 |
| 1.67807 | -0.004 | 0.07 |
| 1.71536 | -0.006 | 0.073 |
| 1.75265 | -0.006 | 0.075 |
| 1.78994 | -0.006 | 0.08 |
| 1.82723 | -0.004 | 0.084 |
| 1.86452 | -0.004 | 0.093 |
| 1.90181 | -0.01 | 0.1 |
| 1.9391 | -0.011 | 0.106 |
| 1.97639 | -0.013 | 0.11 |
| 2.01368 | -0.021 | 0.104 |
| 2.05097 | -0.019 | 0.097 |
| 2.08826 | -0.016 | 0.086 |
| 2.12555 | -0.013 | 0.073 |
| 2.16284 | -0.01 | 0.05 |
| 2.20013 | -0.011 | 0.036 |
| 2.23742 | -0.015 | 0.022 |
| 2.27471 | -0.011 | 0.002 |
| 2.31201 | -0.011 | -0.014 |
| 2.3493 | -0.012 | -0.028 |
| 2.38659 | -0.013 | -0.035 |
| 2.42388 | -0.011 | -0.043 |
| 2.46117 | -0.01 | -0.05 |
| 2.49846 | -0.013 | -0.06 |
| 2.53575 | -0.014 | -0.065 |
| 2.57304 | -0.012 | -0.066 |
| 2.61033 | -0.013 | -0.078 |
| 2.64762 | -0.017 | -0.089 |
| 2.68491 | -0.017 | -0.096 |
| 2.7222 | -0.018 | -0.101 |
| 2.75949 | -0.017 | -0.109 |
| 2.79678 | -0.014 | -0.118 |
| 2.83407 | -0.011 | -0.128 |
| 2.87136 | -0.009 | -0.135 |
| 2.90865 | -0.006 | -0.142 |
| 2.94594 | -0.006 | -0.137 |
| 2.98323 | -0.005 | -0.141 |
| 3.02052 | -0.002 | -0.153 |
| 3.05781 | -0.005 | -0.152 |
| 3.0951 | -0.007 | -0.156 |
| 3.13239 | -0.005 | -0.158 |
| 3.16968 | -0.003 | -0.161 |
| 3.20697 | -0.002 | -0.165 |
| 3.24427 | 0 | -0.17 |
| 3.28156 | 0.003 | -0.178 |
| 3.31885 | 0.009 | -0.179 |
| 3.35614 | 0.01 | -0.181 |
| 3.39343 | 0.007 | -0.187 |
| 3.43072 | 0.005 | -0.188 |
| 3.46801 | 0.013 | -0.194 |
| 3.5053 | 0.019 | -0.2 |
| 3.54259 | 0.017 | -0.201 |
| 3.57988 | 0.014 | -0.196 |
| 3.61717 | 0.017 | -0.2 |
| 3.65446 | 0.027 | -0.204 |
| 3.69175 | 0.031 | -0.206 |
| 3.72904 | 0.034 | -0.21 |
| 3.76633 | 0.039 | -0.212 |
| 3.80362 | 0.046 | -0.207 |
| 3.84091 | 0.048 | -0.212 |
| 3.8782 | 0.038 | -0.215 |
| 3.91549 | 0.038 | -0.216 |
| 3.95278 | 0.039 | -0.217 |
| 3.99007 | 0.042 | -0.216 |
| 4.02736 | 0.045 | -0.224 |
| 4.06465 | 0.054 | -0.225 |
| 4.10194 | 0.264 | -0.222 |
| 4.13923 | 1.235 | -0.227 |
| 4.17653 | 1.127 | -0.229 |
| 4.21382 | 0.54 | -0.236 |
| 4.25111 | 0.361 | -0.243 |
| 4.2884 | 0.299 | -0.233 |
| 4.32569 | 0.28 | -0.238 |
| 4.36298 | 0.287 | -0.245 |
| 4.40027 | 0.307 | -0.246 |
| 4.43756 | 0.332 | -0.238 |
| 4.47485 | 0.358 | 0.371 |
| 4.51214 | 0.38 | 1.093 |
| 4.54943 | 0.397 | 0.448 |
| 4.58672 | 0.4 | 0.069 |
| 4.62401 | 0.4 | -0.05 |
| 4.6613 | 0.392 | -0.104 |
| 4.69859 | 0.373 | -0.134 |
| 4.73588 | 0.342 | -0.159 |
| 4.77317 | 0.307 | -0.181 |
| 4.81046 | 0.28 | -0.193 |
| 4.84775 | 0.268 | -0.196 |
| 4.88504 | 0.261 | -0.204 |
| 4.92233 | 0.264 | -0.215 |
| 4.95962 | 0.272 | -0.219 |
| 4.99691 | 0.279 | -0.221 |
| 5.0342 | 0.282 | -0.227 |
| 5.07149 | 0.276 | -0.23 |
| 5.10879 | 0.258 | -0.235 |
| 5.14608 | 0.24 | -0.243 |
| 5.18337 | 0.215 | -0.248 |
| 5.22066 | 0.184 | -0.248 |
| 5.25795 | 0.161 | -0.254 |
| 5.29524 | 0.139 | -0.261 |
| 5.33253 | 0.119 | -0.263 |
| 5.36982 | 0.1 | -0.264 |
| 5.40711 | 0.086 | -0.266 |
| 5.4444 | 0.073 | -0.269 |
| 5.48169 | 0.065 | -0.274 |
| 5.51898 | 0.055 | -0.281 |
| 5.55627 | 0.048 | -0.282 |
| 5.59356 | 0.041 | -0.287 |
| 5.63085 | 0.041 | -0.289 |
| 5.66814 | 0.031 | -0.285 |
| 5.70543 | 0.025 | -0.29 |
| 5.74272 | 0.016 | -0.297 |
| 5.78001 | 0.008 | -0.3 |
| 5.8173 | 0.005 | -0.298 |
| 5.85459 | 0.001 | -0.296 |
| 5.89188 | -0.006 | -0.301 |
| 5.92917 | -0.013 | -0.304 |
| 5.96646 | -0.013 | -0.302 |
| 6.00375 | -0.015 | -0.308 |
| 6.04105 | -0.021 | -0.31 |
| 6.07834 | -0.025 | -0.305 |
| 6.11563 | -0.03 | -0.305 |
| 6.15292 | -0.035 | -0.309 |
| 6.19021 | -0.039 | -0.312 |
| 6.2275 | -0.04 | -0.311 |
| 6.26479 | -0.039 | -0.311 |
| 6.30208 | -0.045 | -0.313 |
| 6.33937 | -0.047 | -0.316 |
| 6.37666 | -0.051 | -0.316 |
| 6.41395 | -0.051 | -0.316 |
| 6.45124 | -0.052 | -0.318 |
| 6.48853 | -0.054 | -0.319 |
| 6.52582 | -0.055 | -0.315 |
| 6.56311 | -0.062 | -0.316 |
| 6.6004 | -0.067 | -0.315 |
| 6.63769 | -0.07 | -0.314 |
| 6.67498 | -0.074 | -0.314 |
| 6.71227 | -0.078 | -0.311 |
| 6.74956 | -0.086 | -0.316 |
| 6.78685 | -0.083 | -0.309 |
| 6.82414 | -0.079 | -0.307 |
| 6.86143 | -0.083 | -0.299 |
| 6.89872 | -0.089 | -0.295 |
| 6.93602 | -0.093 | -0.29 |
| 6.97331 | -0.098 | -0.284 |
| 7.0106 | -0.099 | -0.272 |
| 7.04789 | -0.101 | -0.263 |
| 7.08518 | -0.107 | -0.255 |
| 7.12247 | -0.112 | -0.241 |
| 7.15976 | -0.115 | -0.23 |
| 7.19705 | -0.117 | -0.222 |
| 7.23434 | -0.116 | -0.22 |
| 7.27163 | -0.118 | -0.204 |
| 7.30892 | -0.124 | -0.191 |
| 7.34621 | -0.12 | -0.184 |
| 7.3835 | -0.118 | -0.177 |
| 7.42079 | -0.118 | -0.172 |

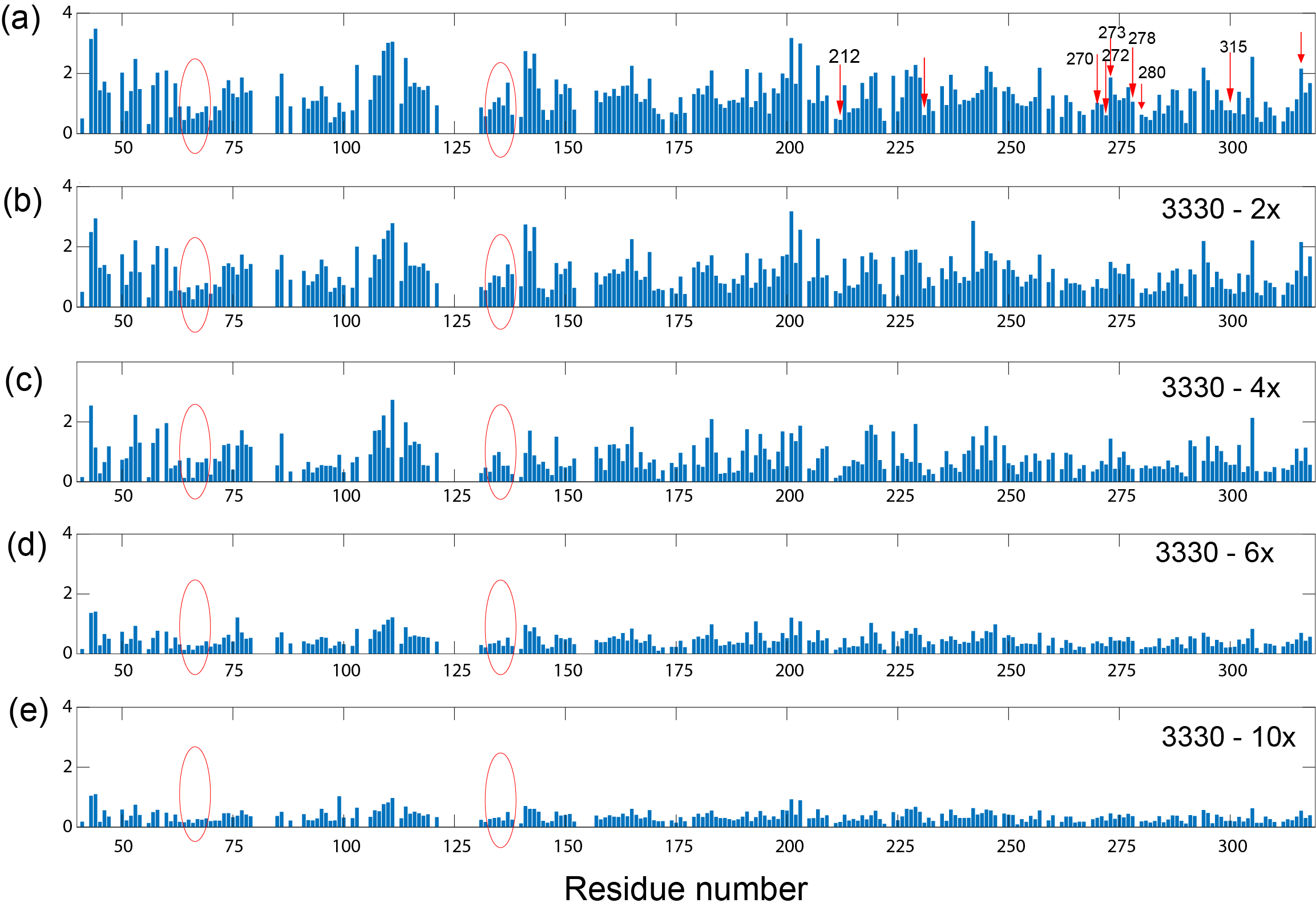

**Figure S22.** 2D ^1^H-^15^N HSQC peak intensities at varying concentrations of APX3330. Peak intensity with increasing APX3330 concentration (a) native protein (b) 2-fold APX3330 (c) 4-fold APX3330 (d) 6-fold APX3330 and (e) 10-fold APX3330. Although several peaks show decreased intensity and/or a chemical shift variation, the variations in peak intensity affect more peaks and are more pronounced than chemical shift variations. The peak intensities indicate that APX3330 shows local as well as global effects; local effects include the small binding pocket of APE1 for which peak intensities are significantly affected by APX3330 as indicated by the red ovals. In contrast, binding interactions with the endonuclease active site for which residues with significant CSPs are indicated with red arrows in panel (a) do not persist throughout the course of the titration. By the 10X point, the intensities of the peaks are similar to those of the surrounding peaks. All 2D ^1^H-^15^N NMR HSQC spectra were acquired with 2048 points in the direct F2 dimension (^1^H) and 256 points in the F1 dimension (^15^N).

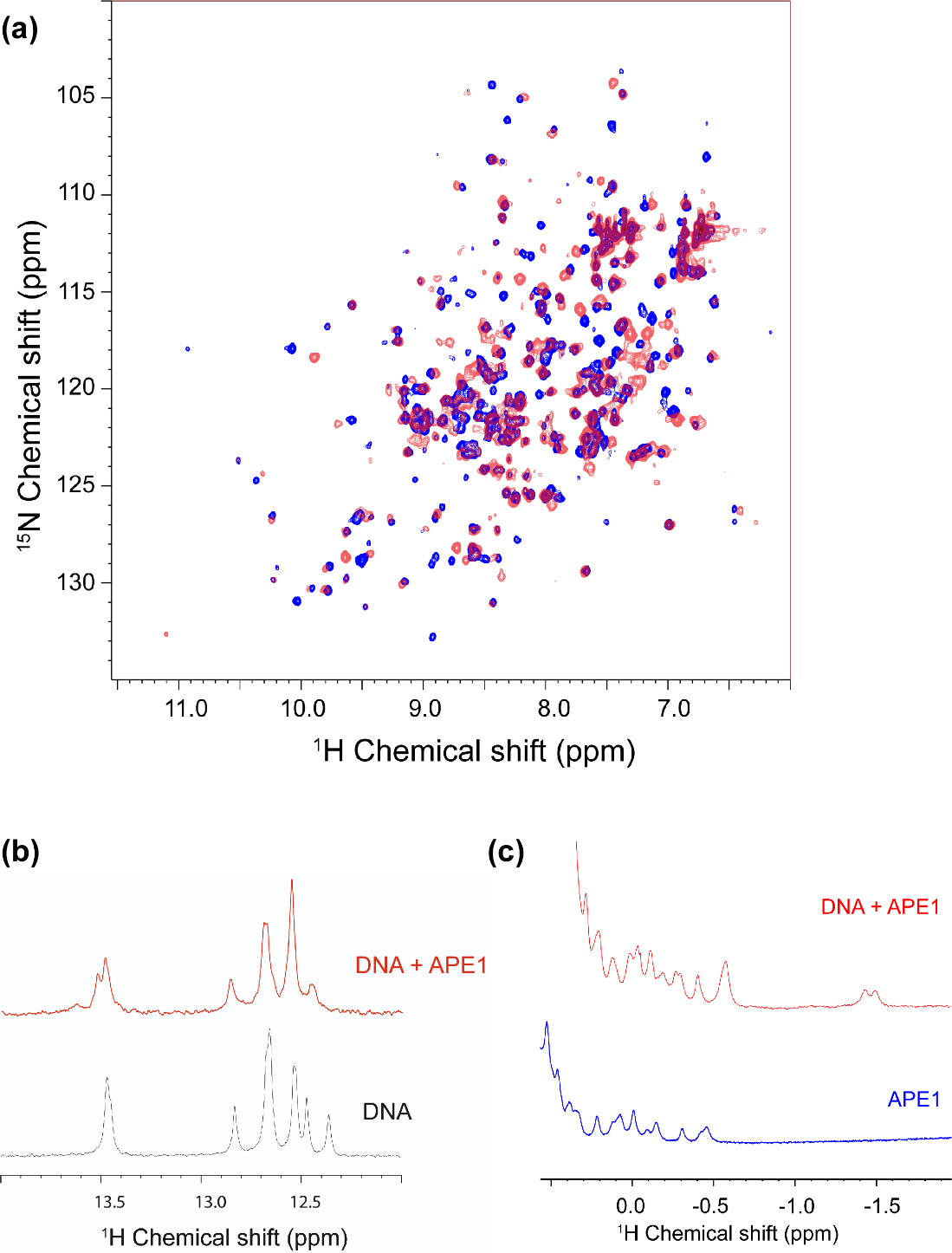

**Figure S23.** The binding of a DNA substrate mimic to APE1 produces a well-resolved ^1^H-^15^N HSQC spectrum. (a) Overlay of 2D ^1^H-^15^N HSQC spectra for free APE1 (blue) with APE1 in the presence of an equimolar amount of DNA (red). Many of the perturbations are too large to confidently assign without additional triple resonances experiments. All spectra were acquired at 25 °C. (b) The imino region of a 1D spectrum of the DNA in the presence and absence of APE1 suggests specific changes in chemical shifts in the DNA consistent with binding. (c) The methyl region of the 1D ^1^H spectrum of the protein shows the emergence of a new peak in the presence of the DNA consistent with a specific interaction between APE1 and the DNA. In the crystal structure of APE1 bound to DNA, R177 and M270 intercalate at the abasic site within the DNA duplex. All of the experiments were collected at 600 MHz Bruker AVANCE spectrometer in TCI cryoprobe.

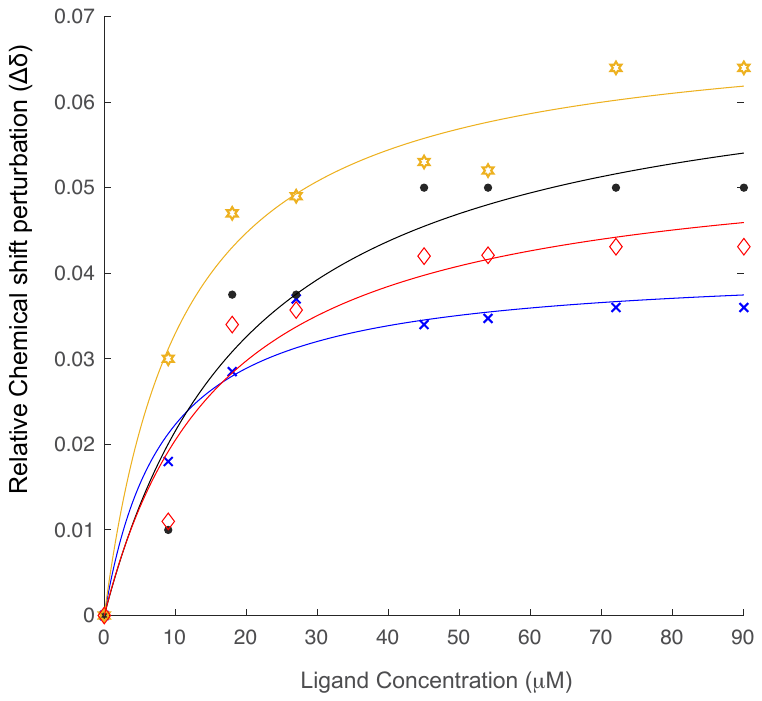

**Figure S24.** The binding affinity of APE1 for DNA was determined using CSPs. The DNA binding affinity was determined from the dependence of shift perturbation (Δδ) on the DNA concentration using MATLAB for multiple residues giving a Kd = 14± 6 μM. Fitting included data for Leu134 (●), Lys299(◇), Leu92(*), and Gly241(X).

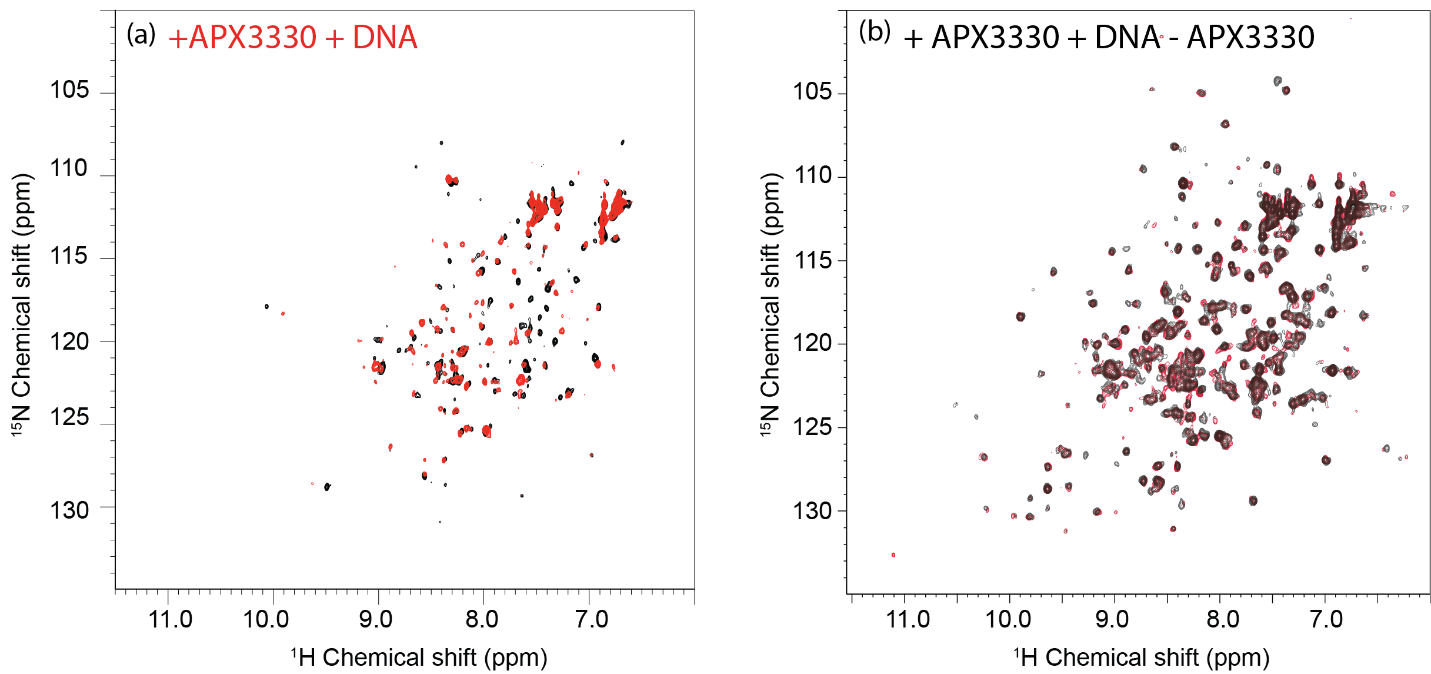

**Figure S25.** Chemically induced partial unfolding of APE1 is reversible in the presence of DNA following removal of APX3330. (a) Following a 2 day incubation with APX3330 at an 8-fold molar excess for which the ^1^H-^15^N HSQC spectrum of [*U*-^15^N]-APE1 is shown (black), DNA substrate mimic was added at a stochiometry of 1:1 (DNA:protein) (red); only subtle changes are observed. (b) The ^1^H-^15^N HSQC spectrum of [*U*-^15^N]-APE1 treated with APX3330, followed by addition of DNA, and then dialysis to remove APX3330 is shown in black. For comparison, the ^1^H-^15^N HSQC spectrum of APE1-DNA (1:1) complex is shown in red (close-up regions shown in SI Fig. S17). The comparison suggests that APE1 interacts with DNA following removal of APX3330 by dialysis, which in turn implies that APE1 is folded as the ^1^H-^15^N HSQC spectra obtained is similar to that of the APE1-DNA complex.

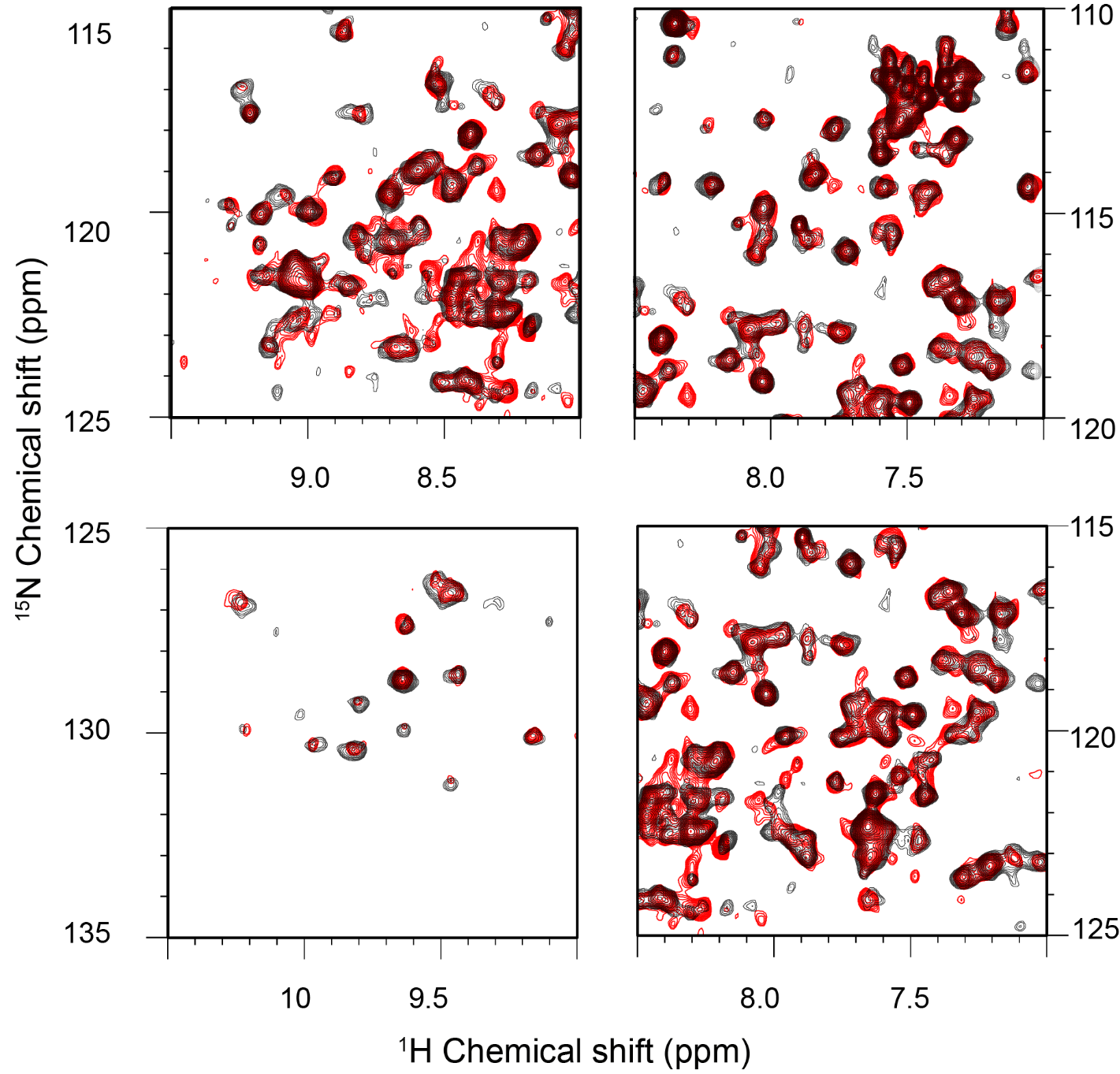

**Figure S26.** The expanded ^1^H-^15^N HSQC spectrum of [*U*-^15^N]-APE1 treated with APX3330, followed by the addition of DNA, and then dialyzed to remove APX3330 is shown in black. For comparison, the HSQC spectrum of the APE1-DNA (1:1) complex is shown in red. The comparison illustrates that under the conditions of the experiment, APE1 folds as in a native state to interact with DNA.  All ^1^H-^15^N HSQC spectrum of [*U*-^15^N]-APE1(90 μM) was collected at 600 MHz Bruker AVANCE spectrometer is equipped with a 5-mm triple-resonance cryoprobe at 25 °C. The 2D ^1^H-^15^N NMR HSQC spectra were acquired with 2048 points in the direct F2 dimension (^1^H) and 256 points in the F1 dimension (^15^N).
